# The Vertebrate Genomes Project Phase I: A global reference genome resource

**DOI:** 10.64898/2026.06.24.732306

**Authors:** Giulio Formenti, Dominic E. Absolon, Linelle Ann L. Abueg, Francine Ackerman, Farooq O. Al-Ajli, Alexandre Aleixo, Agostinho Antunes, Dahiana Arcila, Jean-Nicolas Audet, Jennifer R. Balacco, Alan J.S. Beavan, Katherine Belov, Ricardo Betancur-R, Shelby L. Bidwell, Matthew T. Biegler, Iliana Bista, Mark Blaxter, Kevin Boyer, Ingo Braasch, Ian R. Bradbury, Nadolina Brajuka, Anne M. Bronikowski, Tom Brown, George Butler, Shuo Cao, Eduardo Charvel, Ying Chen, Claudio Ciofi, Hiram M. Clawson, Joanna C. Collins, Solenne Correard, Shannon Corrigan, Mélanie Couture, Andrew J. Crawford, Nathália D. D’Alessandre, Helena B. da Conceicao, Rute R. da Fonseca, Minoli Daigavane, Jeronymo Dalapicolla, Mark de Bruyn, Emma de Jong, Diego N. De Panis, H. William Detrich, Carol DeWeese Scott, Mark Diekhans, Erick Duarte, Richard Durbin, Merly Escalona, Michel Eschenbrenner, Olivier Fedrigo, Jonathan Fenn, Patrick Flannery, Niamh Forde, Simone M. Gable, Pedro A. Galante, Rajee Ganesan, Amanda Gardiner, Erik Garrison, Neil J. Gemmell, M Thomas P Gilbert, François Giudicelli, Tássia M. Gonçalves, Gabriela D.A. Guardia, Andrea Guarracino, Anshu Gupta, Bettina Haase, Delson Hays, Glenn Hickey, Michael Hiller, Paul M. Hime, Carolyn J. Hogg, Kathleen M. Horan, Kerstin Howe, Danny Jackson, Nivesh Jain, Lin Kang, Sergei Kliver, Byung June Ko, Klaus-Peter Koepfli, Aleksey Komissarov, Lisa M. Komoroske, Bonhwang Koo, Sergey Koren, Jonas Korlach, Iva Kovacic, Ksenia Krasheninnikova, Lukas F. Kuderna, Ryan A. Langlois, Delphine Lariviere, Denis M. Larkin, Chul Lee, Young Ho Lee, Harris A. Lewin, Daven Lim, Runyang Nicolas Lou, Sarah S. Mak, Kateryna D. Makova, Laia Marín-Gual, Tomas Marques-Bonet, Fergal J. Martin, Thomas C. Mathers, Camila J. Mazzoni, Kirsty McCaffrey, Shane A. McCarthy, Jack A. Medico, Rafael L.V. Mercuri, Axel Meyer, Pawel Michalak, Siavash Mirarab, Phillip A. Morin, Jacquelyn Mountcastle, Robert W. Murphy, Terence D. Murphy, Eugene W. Myers, Gavin J. Naylor, Thomas Near, Anton Nekrutenko, Zemin Ning, Ben J. Novak, Michelle F. O’Brien, Mary J. O’Connell, Brian P. O’Toole, Sadye Paez, Benedict Paten, Michael Paulini, Sarah E. Pelan, Matt Pennell, Andreas R. Pfenning, Adam M. Phillippy, Brendan Pinto, Martin Pippel, Marina Popova, Maksym Prylutskyi, Arang Rhie, Hugues Roest Crollius, Aurora Ruiz-Herrera, Oliver Ryder, Yana Safonova, Camilla A. Santos, Michael C. Schatz, Simona Secomandi, H. B. Shaffer, Lauren Shalmiyev, Beth Shapiro, Jeramiah J. Smith, Marco Sollitto, Cibele G. Sotero-Caio, Cynthia C. Steiner, Peter H. Sudmant, Deacon J. Sweeney, Samuel C. Talbot, Christopher H. Taylor, Emma C. Teeling, Françoise Thibaud-Nissen, Tatiana Tilley, Nataliya Timoshevskaya, Patrick Traore, Yatish Turakhia, Marcela Uliano-Silva, Chris Venditti, Byrappa Venkatesh, Sonja C. Vernes, Wesley C. Warren, Conor V. Whelan, Melissa A. Wilson, Sylke Winkler, Jonathan M. Wood, Guojie Zhang, Guoyan Zhao, Erich D. Jarvis, the Vertebrate Genomes Project Consortium Phase I

**Author notes:** Contact authors: Giulio Formenti, Erich D. Jarvis. See Extended Author List Table (the Vertebrate Genomes Project Consortium Phase I).

## Abstract

The Vertebrate Genomes Project (VGP) aims to produce complete and near-error-free reference genomes for all ∼70,000 extant vertebrate species^1^. Organized in four phases, it progressively targets all vertebrate orders, families, genera, and eventually all species. Here we present the completion of VGP Phase I, delivering reference genomes for ∼95% of vertebrate orders, along with additional lineages within those orders, totaling 816 species and 1.6 trillion base pairs of main haplotype sequence. These genomes were assembled and annotated over an 8-year period (2018-2026) of rapid advances in genome sequencing, assembly, and annotation methods^2–4^, alongside the growth of associated consortium initiatives and international collaborations^5–9^. They represent some of the highest-quality vertebrate genomes currently available, and most have become the primary reference for their respective species in public databases. Comparative analyses across a subset of 579 species when we reached a threshold of 85% of orders allowed us to reconstruct the genome of the last common ancestor of all vertebrates 500 million years ago, identify diverse modes of sex chromosome evolution, reveal clade-specific three-dimensional genome architecture, discover methylated epigenetic landscapes across vertebrates, and provide a framework for studying gene and pseudogene evolution, immune loci, cancer-associated genes, and other trait-associated loci. Approximately a quarter of this subset are listed as Vulnerable to Critically Endangered by the IUCN Red List of Threatened Species, and have enabled more advanced genomic investigations of extinction risk. VGP Phase I delivers a reference backbone for vertebrate genomics, enabling discoveries that would otherwise remain out of reach across evolution, conservation, and medicine.

**Talking points:**

1. The flagship paper of VGP Phase I
2. The highest quality collection of genomes within the eukaryotic domain of life
3. Evolution of genome sequencing technology quality throughout VGP Phase I
4. A driver project that has been a model for multiple large-scale, high-quality reference genome projects
5. Releases all currently unpublished genomes in Phase I from scientific study embargoes
6. Multiple biological discoveries across the vertebrate tree of life

## Introduction

The Vertebrate Genomes Project (VGP) was established in 2017 with the mission to generate high-quality, near-complete, and near error-free reference genomes for all extant vertebrate species, an estimated 70,000 species spanning over 500 million years of evolutionary history^1^. The initiative is organized into four progressive, phylogenetically inclusive phases: beginning with at least one representative species per order (Phase I), followed by representatives at the family (Phase II) and genus (Phase III) levels, and ultimately all species (Phase IV). The overarching goal is to establish a comprehensive and accurate genomic resource for biology, supporting comparative and evolutionary genomics, biodiversity conservation, and biomedical research.

The VGP evolved from the Genome 10K (G10K) Consortium established in 2009^10,11^, as the first large-scale initiative of its kind for multicellular organisms, with the goal of sequencing 10,000 vertebrate species. During the first iteration of the project, 100 species were sequenced, mostly with short 100-250 bp reads. Although these genomes enabled important discoveries^12–15^, it soon became apparent that their assembly quality was insufficient for other scientific goals, like determining gene functions, gene family expansions, and chromosome evolution^1,16–18^. In the second iteration, as the VGP, focus was placed on developing genome sequencing technologies and algorithms that could achieve near-complete and near-error-free assemblies. Lessons learned by 2021 from a hummingbird and 24 other vertebrate species demonstrated that: long reads exceeding 10 kbp were necessary to achieve highly contiguous vertebrate assemblies, especially through repetitive regions; sequencing chemistries capable of traversing GC-rich regions were required; and that phasing paternal and maternal haplotypes before or during assembly was essential to prevent false haplotype duplications and related errors^1,19^. Based on these lessons, the VGP set minimum metrics for high-quality genome assemblies, including contig N50 ≥ 1 Mbp, scaffold N50 ≥ 10 Mbp, haplotype phasing, base-call quality values (QV) ≥ 40, chromosome-level assignments, and manual curation of structural errors^1^. These metrics and the VGP protocols were adopted by additional consortia that formed with overlapping interests, most under the Earth Biogenome Project (EBP) umbrella^5^, whose mission is to coordinate a network of efforts to produce high-quality reference genomes of all 1.8 million named eukaryotic species, including vertebrates. Concurrent advances in sequencing technology and assembly algorithms further enabled the generation of telomere-to-telomere (T2T) complete diploid vertebrate genomes, initially in humans^20,21^ and subsequently in other species^22–24^.

Here, we report the completion of VGP Phase I, delivering one or more high-quality reference genome assemblies for 816 species, representing 95% of a targeted 271 VGP that includes all orders, lineages defined by divergence times, and four sister deuterostome invertebrate outgroups, spanning the vertebrate tree of life, covering more than 500 million years of vertebrate evolution^25^ and 635 million years of animal evolution^26^ (**Supplementary Table 1**). These assemblies were generated between 2018 and 2026, meet VGP^1^ and EBP^27^ quality standards (Howe et al., in preparation), and range from partially phased^1,28^, fully phased diploid assemblies^28–30^, to complete T2T diploid assemblies^20,22^. Genomes were generated by a global collaborative effort of over 26 organizations including: the VGP^1^, Bat1K^9^, Cetacean Genomes^31^, Darwin Tree of Life (DToL)^7^, European Reference Genome Atlas (ERGA)^8^, Denmark Yggdrasil^32^, Genomics of the Brazilian Biodiversity (GBB)^33^, Minderoo Australian Oceanomics^34^, French ATLASea^35^, African BioGenome Project (AfricaBP)^36^, and T2T consortia^20^, among others (**Supplementary Table 1, Columns AJ and AK**). With a datafreeze subset of 579 species completed by April 2025 (**Supplementary Table 2**), we illustrate how this VGP Phase I genomes can be used to address broad biological questions, including reconstruction of the ancestral vertebrate genome, comparative analyses of gene families and specialized traits, and investigations in conservation genomics.

## Results

### A high-quality phylogenomic resource

#### Species selection

VGP Phase I species selection was determined by a combination of phylogenetic breadth across orders, value to the scientific community, benefits to society and human health, extinction risk, and availability of high-quality samples (**Supplementary Table 3**). For phylogenetic breadth, because classification systems are often not equally applied across groups of species and vary according to different criteria, we sought an additional quantitative measure of species selection that could be applied uniformly. We noted that for fossil-dated family trees of birds^15^ and mammals^37^ inferred from nuclear genomic sequences, most clades ranked as orders had stem divergence ages between 70 to 50 million years ago (MYA), around the Cretaceous-Paleogene mass extinction; for ray-finned fish, most orders had stem divergences between 120 to 50 MYA^38^. We thus applied an additional selection criterion for Phase I clades, as lineages that diverged from their nearest extant sister lineage between 100 to 50 MYA, inferred from timetree phylogenies based on nuclear genomic sequences, available as of 2018 (**Supplementary Table 4**)^15,37–50^. This resulted in the expansion from 136 to 267 ordinal equivalent lineages of vertebrates according to divergence times (**Supplementary Table 2, Column I**): from 30 to 39 mammal groups, mostly due to expansion in rodents; 40 to 52 bird groups, mostly due to expansions in Pelecaniformes and Caprimulgiformes; 4 to 33 non-avian reptile groups, mostly due to expansions in turtle and lizard families; 3 to 30 amphibian groups, mainly due to expansions in frog and salamander families; 72 to 93 fish groups, based on previous ordinal definitions for bony fish, sharks, and rays^51,52^. Nearly all (98%) previous taxonomically defined orders had inferred divergence times with their closest sister order of ∼50 MYA or earlier (4 exceptions had 40-45 MYA divergence times, all birds: Woodpeckers vs Bee eaters; Hummingbirds vs Swifts; Tinamous vs Emus; Kiwis vs Rheas; **Supplementary Table 2**, **Column I**). Splitting primates into catarrhines (humans, other apes, and old world monkeys) and platyrrhines (new world monkeys), Passeriformes into oscines (vocal learning songbirds) and suboscines (mostly non-vocal learners), and adding four Deuterostome invertebrate sister outgroup lineages (lancelets/amphioxus, tunicates, sea stars, acorn worms) brought the total to 271 targeted VGP Phase 1 lineages.

Over an 8-year period, we generated genome assemblies for ∼95% (257 of 271) of the targeted VGP Phase I lineages (**Fig. 1a**, **Supplementary Table 2**). A similar proportion was achieved when considering formally recognized vertebrate orders in the current NCBI taxonomy^53^ (∼96%, 154/161, including the outgroups). Of these 257, 29 are monotypic and 9 are ditypic lineages, with just 1 or 2 living species, respectively, that diverged from their nearest living relative species 50 MYA or more. For the 14 missing targeted lineages without a high-quality reference genome yet, 7 are mono- or ditypic, and thus it is more difficult to obtain a representative sample. The next most principled obstacle to generating high-quality reference genomes was difficulty in obtaining sufficient sample quality or quantity, or the length of time it took to obtain the necessary permits (**Supplementary Table 5**).

**Fig. 1.**
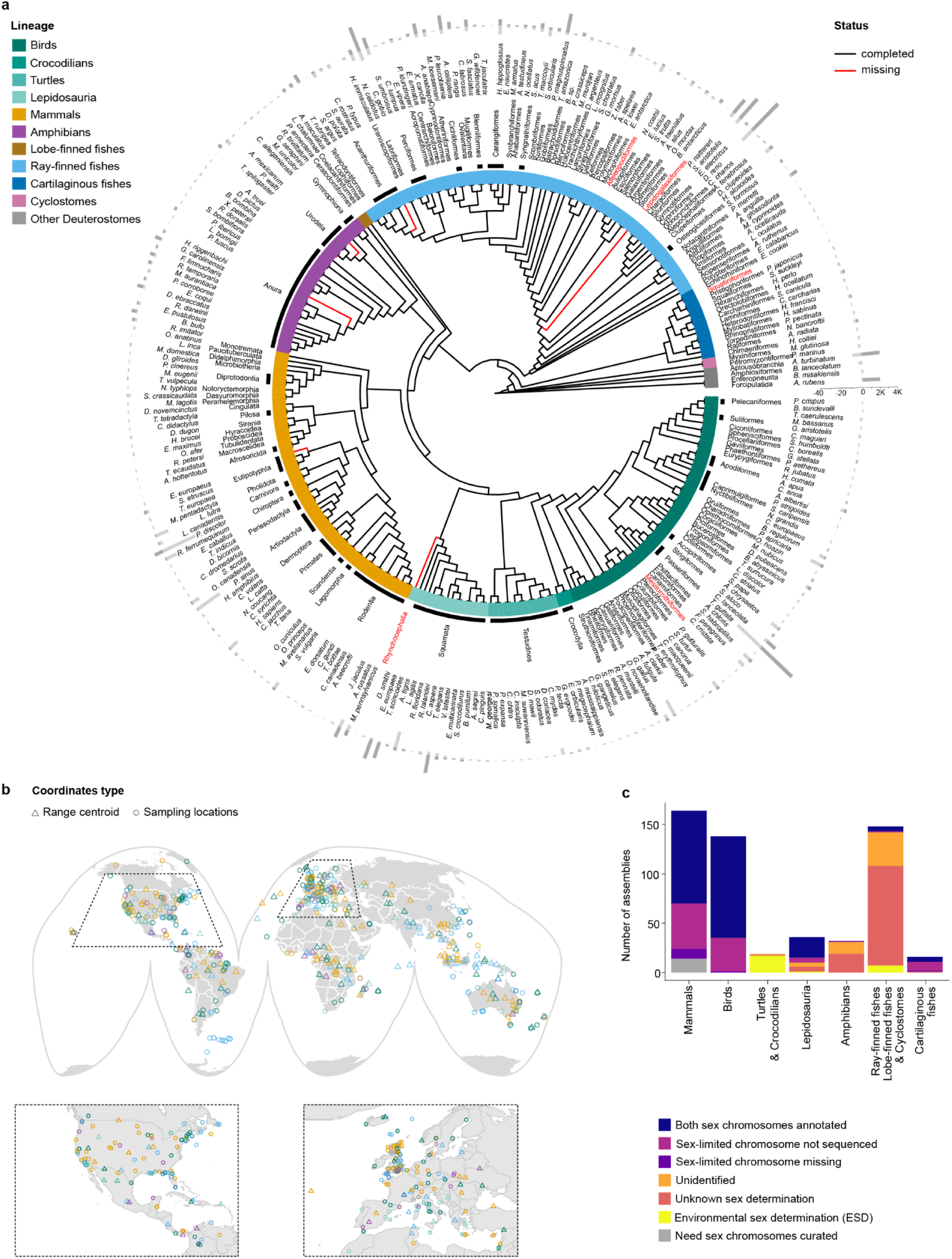
Ordinal-level lineages and their species included in the VGP Phase I dataset. **a**, Targeted lineages according to the NCBI ordinal taxonomy as of 2025 (inner labels; ∼96% sequenced, 154/161) and VGP criterion of 50-100 MYA divergence time window (outer representative species labels; ∼95%, 257/271). Red branches and text, missing target lineages. Outer circle contains bar plots that represent the estimated number of extant species per targeted lineage (outer bars) and number sequenced in the VGP Phase I dataset (inner bars). The phylogenetic tree is from Open Tree of Life^69^; a more updated tree will be presented in a separate study on phylogeny. **b,** Geographic sampling locations of VGP phase I 579 species. Circle, collection sites from wild populations. Triangle, range centroid for species without reported sampling locations, or obtained from zoos, aquaria, or laboratories. Colors indicate major clades as in (**a**). North America and Europe (delineated by dashed lines and expanded below) contained the highest densities of sample locations. **c,** Proportions of sex chromosomes or sex determining systems among the VGP 579 datafreeze assemblies (y-axis and color code) among 7 major vertebrate groupings (x-axis).

We also included additional species within any order or temporally-defined lineage from collaborating projects that followed similar genome sequencing and assembly protocols to reach the VGP metric thresholds. Selection of additional species was driven in part by communities of scientists focused on specific projects and questions. We also reached out to labs that followed our protocols and generated genomes of new species, to request adding their genomes to the VGP Phase I collection, where we sometimes improved the assembly further. The highest number of species per targeted lineage includes: 28 cetaceans and 65 bats among mammals, spearheaded by the Cetacean and Bat1K genome projects, respectively; 45 songbirds and 25 waterfowl, spearheaded by the vocal learning and Darwin Tree of Life initiatives, respectively; 15 Lacertidae wall lizards among reptiles, by the Wellcome Sanger Institute Tree of Life Programme; and 25 perciforms and 21 wrasses among fishes, spearheaded by the ichthyological science community. The combined result is 816 species in the VGP Phase I collection (**Extended Data Fig. 1**, **Supplementary Table 1**), sourced from all seven continents, multiple islands, and all five oceans (**Fig. 1b**).

In past sequencing efforts, the heterogametic sex was often avoided due to challenges in assembling highly divergent sex chromosomes, such as lower sequencing coverage relative to the autosomes^54^. In the VGP and collaborating projects, we focused on improving methods for sequencing and assembling sex chromosomes to high quality^1,28,55^, and thereby purposely selected the heterogametic sex when available (e.g. a male for XX/XY species; a female for ZZ/ZW species). Of the 816 species, this resulted in sequencing both sex chromosomes for 144 XY and 147 ZW heterogametic individuals, and four species with 3 or more sex chromosomes (**Fig. 1c**, **Supplementary Table 1, Column T**; Gable et al., to be submitted to bioRxiv).

Most analyses in this study were conducted on a subset of the 816 species, a VGP data freeze in April 2025 comprising 579 species, when the consortium reached a threshold of 85% of the targeted 271 lineages (**Supplementary Table 2**). Analyses incorporating some or all of the species are indicated where applicable. Metadata associated with analyses subsets can be retrieved in resources described in the companion studies (Sotero-Caio et al., to be submitted to bioRxiv; Clawson et al., in preparation). More details on species selection is in **Supplementary Information section 1**.

### Genome assembly quality

The genomes included in the VGP Phase I data set were generated during a period of rapid evolution of sequencing technologies (**Extended Data Fig. 2**), all with the criteria of using long reads as their basis for generating contigs, scaffolding of contigs into chromosomes, haplotype phasing (partial to complete), manual curation of structural errors, chromosome naming, and removal of contaminating sequences from other species. Chromosome-level continuity was achieved with Hi-C data^56,57^; Hi-C data was also later used for haplotype phasing^19,29^. Assembly types fell into five categories of increasing quality, as technology and methods improved over the course of the VGP Phase I project^1,28,58–60^ (**Supplementary Table 2**):

1. Single haploid assemblies using Pacific Biosciences (PacBio) Continuous Long Reads (CLR) >10 kbp, partial haplotype phasing, scaffolding with Hi-C^56,57^ and Bionano optical maps^1^, and base call polishing with Illumina short reads^1^, with the resulting haplotypes labelled as a high-quality primary (.pri) and a lower quality alternate (.alt) assembly (n = 140 species; 128 using FALCON unzip assembler, 12 using Canu^61^, IPA^62^, or mercat2^63^ assemblers);
2. Single haploid assemblies using PacBio High Fidelity (HiFi) long reads > 10 kbp for contigs and Hi-C for scaffolding^19,56,57,64,65^, with haplotypes labelled also as high-quality primary (.pri) and lower quality alternate (.alt) assembly (n = 181 species; using HiFiasm^64^ assembler, with 5 species including ONT data);
3. Diploid assemblies in which parental short read data were used to completely phase child long reads into their respective parental haplotypes, coupled with the above approaches to obtain two nearly complete assemblies per individual^1,61^, with haplotypes named paternal (.pat) and maternal (.mat) (n = 28 species, using TrioCanu^30^ or HiFiasm^64^ trio assemblers);
4. Diploid assemblies similar to type 3, but using Hi-C data to chromosomally phase haplotype contigs^29^, with haplotypes per individual labelled as haplotype 1 (.hap1) and haplotype 2 (.hap2) (n = 207 species; using HiFiasm^29^ Hi-C phasing assembler).
5. T2T diploid assemblies using a combination of PacBio HiFi and Oxford Nanopore Technologies (ONT) Ultra-long (≥ 100 kbp) reads, along with parental short read or individual Hi-C data for phasing and/or scaffolding^20,22,23,66^, and haplotypes also labelled as in assembly types 3 or 4 (n = 19 species; using Verkko^60,67,68^ or HiFiasm Hi-C Phasing ONT integrated^59^ assemblers).

Each approach required different assembly algorithms, with the most common used for generating contigs among the Phase I dataset being HiFiasm on PacBio HiFi data in Hi-C phasing mode^29^. These assembly protocol types are set up as pipelines in the Galaxy platform^28^ (https://vgp.usegalaxy.org) (Lariviere et al., to be submitted to bioRxiv) or in nextflowhub for Tree of Life@Sanger assemblies (https://workflowhub.eu/collections/15/programmes). We selected the highest quality haplotype that met or surpassed the VGP metrics (labelled it hap1 for the Hi-C phased diploid assemblies), added the sex chromosomes when present in the other haplotype of diploid assemblies of heterogametic individuals, and included the mitochondrial genome when recovered. We designated this high-quality combination as the main haplotype (**Supplementary Table 1, Column Q**), for alignments and analyses that use only one haplotype per species .

Sizes of assembled genomes fell within the expected ranges for each vertebrate clade^70^ (**Fig. 2a**). VGP genomes were typically larger than those obtained by non-VGP protocols for the same or closely related species (**Fig. 2b**, **Supplementary Table 6**). Among the 579 species, ray-finned fishes had the smallest genomes but exhibited a broad range of sizes (0.3-3.5 Gbp), whereas birds possessed compact genomes with more limited size variation (1.0-1.4 Gbp), and amphibians showed the greatest variation (1-30 Gbp). The greater pipefish (*Syngnathus acus*, ray-finned fish) possessed the smallest genome (0.324 Gbp), and the hellbender salamander (*Cryptobranchus alleganiensis*) possessed the largest (48.8 Gbp), followed by the West African lungfish (*Protopterus annectens*, Dipnoi; 40.5 Gbp). The average across the vertebrates sequenced herein was 2.14 Gbp (SD 2.88 Gbp). For the invertebrate deuterostome outgroups, genome size range was smaller (0.1-1.0 Gbp). In total, main haplotype sequences summed to ∼1.265 trillion base pairs (Tbp) for the 579 species data freeze and ∼1.600 Tbp for the expanded list of 816 species; the latter figure is equivalent to ∼530 human genomes (3 Gbp size) or ∼3,900 insect genomes (∼0.4 Gbp average size).

**Fig. 2.**
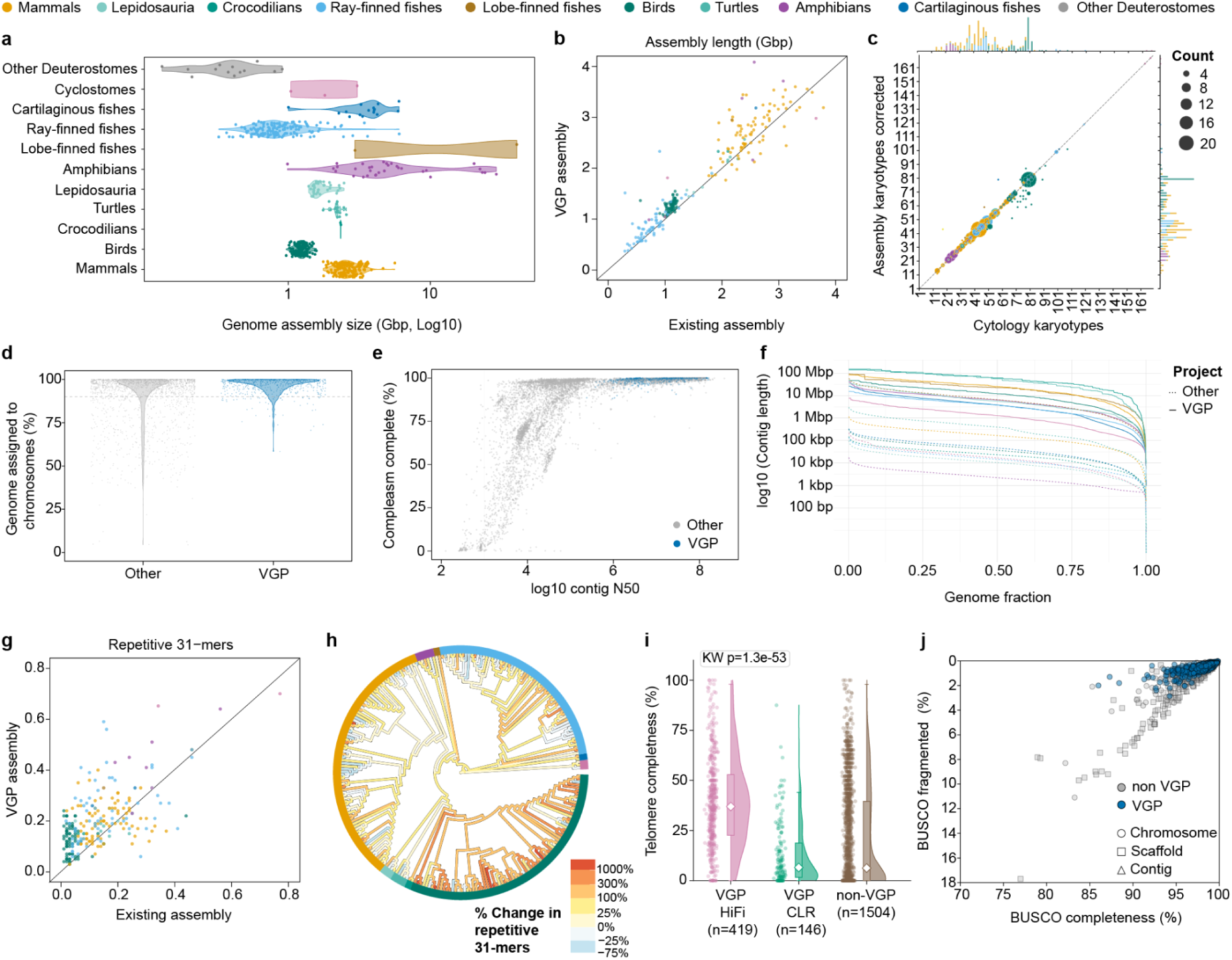
Relative quality of VGP Phase I genomes. **a**, Genome assembly sizes grouped by 10 vertebrate clades and deuterostome invertebrate outgroups. **b,** Total length of the genome assemblies in the VGP data freeze (y-axis) for species that also have an alternative assembly available in GenBank (x-axis). **c,** Concordance between chromosome counts obtained from cytological karyotypes and assembly-derived diploid chromosome numbers. Dot size corresponds to the number of species represented by each paired data point. The frequency distribution of diploid chromosome counts per lineage is shown as histograms. Chromosome counts are corrected for sex, sex determination system, and sex chromosomes in hap1 versus hap2. **d,** Percentage of assembly assigned to chromosomes for non-VGP chromosome-level assemblies in GenBank and in the VGP 579 species data freeze. **e,** Comparison of gene completeness (compleasm %) vs contig N50 for all vertebrate genome assemblies in GenBank, colored-coded blue for VGP 579 data freeze genomes and gray for all others. **f,** Median contig Nx values for the same genome datasets. **g,** Fraction of each assembly composed of 31-mers that occur at least twice in the genome for assemblies with a corresponding genome in the comparator set (VGP, y-axis; non-VGP, x-axis in Genbank). **h,** Percent change in 31-mer repetitiveness in VGP genomes relative to the existing comparator set, with values above zero indicating higher repeat representation in VGP genomes. Colors along internal branches represent ancestral state reconstructions on the ROADIES tree. The hellbender salamander (*Cryptobranchus alleganiensis*) and West African lungfish (*Protopterus annectens*) were excluded owing to their exceptionally large genome assembly sizes (48.8 and 40.5 Gbp, respectively). **i**, Telomere completeness, measured as the percentage of expected chromosome ends with terminal telomere calls, across VGP HiFi, VGP CLR and non-VGP assemblies; Kruskal-Wallis test was used to test for significant differences in distributions. **j**, Percent BUSCO completeness versus BUSCO fragmentation for VGP and non-VGP genomes.

Across the 579 species, 15,009 chromosome-level scaffolds were assembled, curated manually for errors, and named. Of these, 335 species had prior cytogenetically determined karyotype data, and of which 277 (83%) showed complete agreement with assembled chromosome numbers and 25 (7%) showed differences of only +/-2 chromosomes, most of which were among birds (**Fig. 2c**, **Supplementary Table 7**). The remaining 33 (10%) species exhibited larger differences (-26 to +20) between published karyotype and assembled chromosomes, mostly in birds and fishes. These discrepancies may reflect the limited resolution of cytological karyotyping, particularly in birds, certain fishes, and reptiles (taxa which possess numerous microchromosomes that are difficult to distinguish in karyotyping), discrepancies in the identification of sex chromosomes (especially common among fishes), intraspecific variation in chromosome number, or residual assembly errors. Nevertheless, the 90% concordance between cytological karyotypes and assembled chromosomes demonstrates remarkable genome structure accuracy in VGP assemblies.

Of the 579 genomes, 91% had more than 90% of their sequence assigned to chromosomes, compared with 73% of the other chromosome-level vertebrate assemblies in GenBank^71^ (**Fig. 2d**). The VGP genomes also exhibited fewer frameshift errors, affecting 2.2% of BUSCO genes compared with 4.9% in other genomes, largely reflecting the higher accuracy of HiFi reads (**Extended Data Fig. 3a,b**, **Supplementary Table 8**). A total of 543 (96.4%) met the contig (N50 > 1 Mbp), compared with only 30.7% of other vertebrate genomes currently available in GenBank using short reads or other non-VGP protocols (**Fig. 2e,f**). The VGP genomes have gene completeness in the 95-100% range, except cyclostomes (**Extended Data Fig. 3c,d**), whereas many of the other non-VGP genomes have completeness in the near 0 to 95% range; these differences in gene completeness are correlated with contig N50 size (**Fig. 2e; Extended Data Fig. 3e,f**), and reflect different project standards (**Extended Data Fig. 3g**). Among the VGP genomes, mammals and birds showed particularly high gene completeness, exceeding 98% on average, likely due to the better resolution of their BUSCO gene sets rather than assembly quality (**Extended Data Fig. 3c,d**). The larger and/or more repetitive the genome, the harder it was to reach or surpass the standard metrics. The VGP genomes also had a greater proportion of repetitive sequences compared to their short read counterparts when available (**Fig. 2g**), presumably reflecting read length, quality of the HiFi reads, and haplotype phasing. The enhanced representation of repeats was up to 1,000-fold greater, particularly evident in birds, sharks, some ray-finned fishes, and some reptiles (**Fig. 2h**). Indeed, when we annotated canonical telomeric repeats (TTAGGG) across chromosomes using a new software tool, Teloscope (Medico et al., to be submitted to bioRxiv), VGP assemblies using HiFi showed a higher representation of telomeric sequences at the end of chromosome scaffolds relative to earlier VGP PacBio CLR-based assemblies and non-VGP genomes in general (**Fig. 2i**). Among the 579 species, we found a total of 10.5K telomeres in 15K chromosome-level scaffolds; over half (53%) of these chromosome-level scaffolds possessed at least one telomere, but only ∼20% had telomeres at both ends. Complete mitochondrial genomes were also generated for a subset of Phase I species (**Supplementary Table 1, Column Y**); this required suitable long mtDNA reads, whose availability varied with tissue type and fragment size cut offs in library preparations^72^. Overall, use of the VGP sequencing and assembly strategies has produced the largest and most phylogenetically expansive set of high-quality vertebrate genomes to date.

### Gene annotations

We generated comparative, evidence-driven gene models for 97.3% of the main assemblies in the 579 species data freeze, using both the well-established NCBI RefSeq Eukaryotic Genome Annotation Pipeline (EGAP)^4^ (298 genomes) and its now public version EGAPx (Tvedte et al, in preparation) (269 genomes), using transcriptome data sets generated by the VGP consortium for 242 species and non-VGP transcriptomes for an additional 312 species at least at the genus level (Sollitto et al., in preparation; **Supplementary Table 9**). For 13 genomes it was not possible to obtain transcriptomic data even at the genus level, and therefore these were annotated with EGAPx using only related protein sequences aligned with the more accurate ProSplign^73^ instead of Miniprot^74^. In addition, the Ensembl annotators generated annotations for 341 VGP species using their vertebrate annotation pipeline. The annotations of the VGP genomes clustered at very high (90-100%) BUSCO completeness with minimal fragmentation relative to non-VGP assemblies (**Fig. 2j; Supplementary Table 10**), demonstrating that improved genome continuity and completeness enable higher quality gene annotations. Mammalian, avian, and chelonian (turtle) genomes were also annotated with the homology-based method TOGA2 using 2 - 5 reference species (Malovichko et al., to be submitted to bioRxiv). These annotations provided the basis for downstream comparisons of protein-coding gene content, pseudogene and retrogene repertoires, immunoglobulin loci, cancer-associated genes, and gene family birth-and-death dynamics.

### Initial phylogeny and whole-genome alignments

While we leave a comprehensive phylogenetic study of the VGP Phase 1 dataset to future work (Mirarab et al., in preparation), to support downstream analyses, we constructed a tree using a modified version of the ROADIES software tool^75^ (**Extended Data Fig. 4**, **Supplementary Table 11, rows 3-15**). ROADIES takes random loci of a genome in a species, finds and aligns that locus in all other species when present, and infers a tree iteratively until the topology stabilizes. With random loci of 500 bp in length, it took 123,141 loci iterations to reach a stabilized tree on the 579 species. A more detailed explanation is presented in **Supplementary Information section 2**.

The ROADIES tree was used as a guide tree for a whole-genome reference-free Progressive Cactus alignment^76^. Progressive Cactus had to be adapted to handle such a large number of species spanning ∼635 million years of phylogenetic divergence. Salamanders and African lungfish genomes were excluded because their genome sizes were too large for efficient Progressive Cactus processing. The commonly used human GRCh38 and mouse GRCm39 references were added as additional human and mouse haplotypes. For the resultant 577 species (**Supplementary Table 11, rows 19 to 596**), it took 67 CPU years to generate the full genome alignment. We also performed an all-vs-all alignment of the 579 species data freeze, using wfmash^77–79^ and FastGA^80^, generating 336,980 pairwise alignment files in PAF format per method (**Supplementary Table 11, row 17**). Each alignment was indexed using impg, and per-base alignment depth on the human T2T-CHM13v2.0 reference was computed using impg depth. Together, these three alignment resources provide complementary ways to interrogate vertebrate genome evolution at multiple scales: the ROADIES tree provides a common phylogenetic framework, the Progressive Cactus alignment enables reference-free comparison across hundreds of assemblies, and the all-vs-all alignments and impg depth profiles provide locus-level views of conservation and assembly representation on a chosen reference. The alignment also constitutes a vertebrate-scale reference framework for future pan-genomic analyses as intraspecific diversity data become available across VGP species. We next used the generated assemblies, annotations, and alignments to examine broader patterns of genomic content across vertebrates.

### Genomic content across vertebrates

#### Protein-coding gene content

Across vertebrates and the deuterostome outgroups, the number of annotated protein-coding genes was relatively constrained (∼20K on average), but exhibited lineage-specific differences (**Fig. 3a, right; Supplementary Table 12**). Birds consistently harboured the lowest amount of protein-coding genes (median = 16,191 genes [IQR: 15,578–16,804]; range: 14,185–18,621), whereas amphibians and ray-finned fishes harboured the largest numbers, also with broader ranges (amphibian median = 22,117 genes [IQR: 20,947–23,717]; range: 18,296–35,513; ray-finned median = 23,205 genes [IQR: 22,027–24,808]; range: 18,398–45,073). The increase in amphibians was in part driven by elevated gene counts in *Xenopus petersii* (35,513), due to its allotetraploid genome architecture, in which both L and S subgenomic components are represented within the assembly. For ray-finned fishes, the increase was also in part driven by several outlier taxa, Salmoniformes (e.g. salmon and char), Acipenseriformes (e.g. sturgeons), and Cyprinidae (e.g. carps) containing the highest gene counts (range 38,165-45,073) among the VGP Phase 1 dataset, suggestive of either not losing complements of most genes or further expansion after the whole-genome duplication (WGD) event in teleost ray-finned evolution^81^. Cartilaginous fishes (sharks and rays) exhibited a more constrained range of protein-coding gene counts (median 19,806 genes [IQR:18,868–21,957]; range: 14,085–22,886). Non-avian reptiles exhibited an intermediate number of protein-coding genes relative to other groups (16-21K genes; median 19,872), and were similar between crocodilians, turtles, and lizards (**Fig. 3a**). Mammals exhibited a relatively narrow distribution of protein-coding genes (median = 20,078 genes [IQR: 19,236–21,166]; range: 17,643–23,549), indicating a high degree of conservation in gene repertoire size across the clade.

**Fig. 3.**
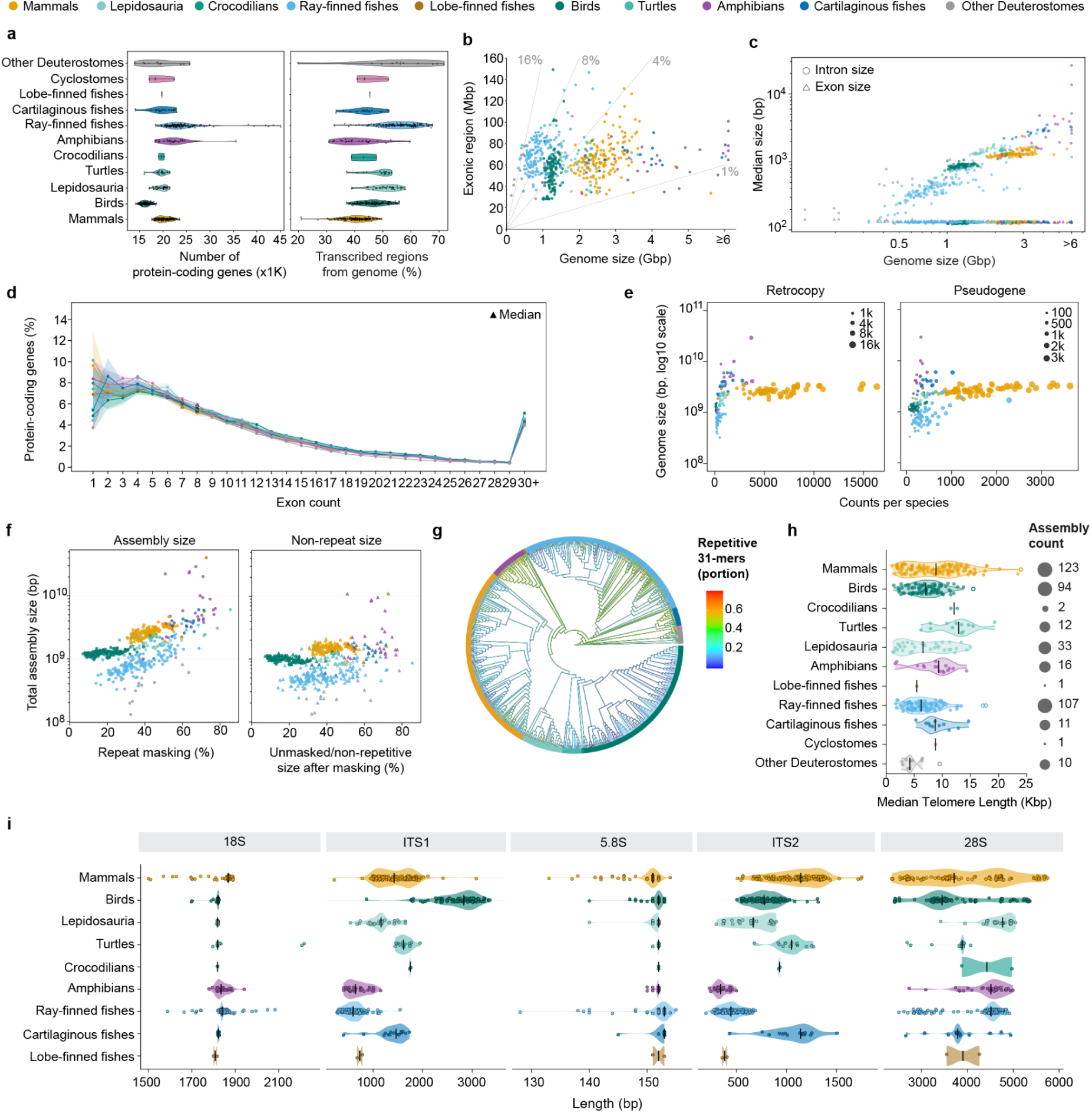
Genomic content across vertebrates across vertebrates. **a**, Total protein-coding gene counts (left) and percentage of the genome transcribed (right) across major vertebrate clades. **b**, Relationship between genome size and total exonic content (coding sequences plus UTRs). Diagonal dashed lines indicate the percentage of the genome occupied by exonic sequences. **c**, Relationship between genome size and median intron and exon lengths per species. **d**, Percentage of protein-coding genes (y-axis) with 1 to 30+ exon counts (x-axis), across clades (color). Circular markers show the mean percentage of genes assigned to each exon-count class for each lineage, with shaded areas representing the standard deviation. The median exon-count class for each lineage is indicated with a triangular marker (most at 7-8 exons). **e**, Processed pseudogene (retrocopies) counts (left) and unprocessed pseudogene counts (right) versus genome size across vertebrate clades. Circle size is proportional to the number of elements. Note different x-axis scales between panels. **f**, Genome assembly sizes with (left) and without (right) repetitive regions included. **g**, ROADIES phylogenetic tree of 579 species, with each branch colored by the portion of 31-mers in an assembly that appear at least twice (i.e., are repetitive); colors of internal branches are based on ancestral state reconstruction. **h**, Median telomere length of HiFi-based assemblies in each major vertebrate clade. **i**, Distributions of species-level median lengths for 18S, ITS1, 5.8S, ITS2 and 28S within the 45S ribosomal RNA gene clusters across vertebrate clades. Violin plots show clade-level density, points indicate individual species, and vertical bars mark clade medians.

On average, the fraction of the genome encompassed by transcribed regions (including exons, introns, and UTR, considering only protein-coding genes) was 47.3% (**Fig. 3a, left**), varying between 20.8% in the golden-mantled ground squirrel (*Callospermophilus lateralis*) to 67.6% in the sea stickleback (*Spinachia spinachia*). The percentage of the genome occupied by transcribed protein-coding regions showed substantial variation across vertebrate lineages. Ray-finned fishes and deuterostome invertebrates displayed the highest median values of 56.1% each. Both groups also showed wide ranges, especially the invertebrates, which varied from 19.8% to 71.8%, suggesting strong heterogeneity. Among reptiles, turtles and Lepidosauria (snakes and lizards) showed relatively high median values, at 50.5% and 50.6%, respectively, while crocodilians had a lower median of 42.0%. Birds showed an intermediate median value of 46.4%, with values ranging from 37.1% to 55.9%. Mammals (median 40.4%), amphibians (median 39.0%), and cartilaginous fishes (median 44.1%) with larger genomes had lower fractions transcribed.

The total exonic sequence (including CDS and UTRs) varied less extensively than genome size, spanning approximately 27.8–149.2 Mbp across species. However, the fraction of the genome occupied by exonic sequence differed markedly among lineages, reflecting differences in genome size and genome architecture (**Fig. 3b**, **Supplementary Table 12**). For example, amphibians had the lowest exonic representation (median 63.7 Mbp; 1.4%) and ray-finned fishes showed the highest (median 69.6 Mbp; 8.2%).

Median intron size scaled positively with increasing genome size across species and showed a strong clade-specific pattern, whereas median exon size of the same genes was highly stable regardless of genome size across vertebrates (**Fig. 3c**, **Supplementary Table 12**). Ray-finned fishes generally had the shortest introns (median 419.0 bp; range 0.2–2.6 kbp), with the reedfish *Erpetoichthys calabaricus* representing a notable outlier, consistent with its large 3.6-Gbp genome. Median intron lengths were also relatively short in deuterostomes invertebrates (median 476.0 bp; 0.17-0.78 kbp) and birds (median 872 bp; 0.74-1.07 kbp), but higher in mammals (∼1.2 kbp) and amphibians (2.5 kbp).

Examining exons further, protein-coding genes displayed comparable exon-number distributions across lineages, with lineage-specific median exon counts consistently centered around 7-8 exons per gene (**Fig. 3d**, **Supplementary Table 12**). However, the proportion of single-exon protein-coding genes varied substantially. Mammals showed the highest mean proportion of single-exon genes (9.6%, partly driven by olfactory receptor genes that in some species can account for up to 60% of single exon genes), followed by amphibians (8.4%), whereas ray-finned fishes and cyclostomes showed the lowest proportion of single-exon genes (4.4 and 3.8% respectively). The deuterostome outgroups showed an even higher proportion of single-exon genes (10.1%; **Fig. 3d**, **Supplementary Table 12**). Together, these results indicate that genomic expansions and contractions among vertebrates were primarily driven by intronic and intergenic changes, but not exons.

### Pseudogene content

The greater resolution of duplicated regions in the VGP assemblies provided an opportunity to quantify pseudogene diversity, conservation, and functional potential across the vertebrate phylogeny, as these regions are often fragmented or collapsed in draft-quality genomes (Mercuri et al., to be submitted to bioRxiv). Pseudogenes can undergo further genetic changes to acquire functionality^82^. In RefSeq annotated VGP genomes, we searched for the two major pseudogene categories: unprocessed pseudogenes, often formed by partial gene duplication and translocation to another locus, and processed pseudogenes (hereafter retrocopies), formed by spliced mRNAs that were reverse-transcribed and inserted back into the genome^83^. We identified retrocopies and pseudogenes in all assemblies of the 579 species data freeze, and found were clear lineage-specific patterns: mammals harboured markedly higher counts of both, especially of retrocopies, and a wider range of counts per species, with some species exceeding 10,000 retrocopies (**Fig. 3e, Supplementary Table 13**). All other lineages had lower counts and narrower ranges, independent of genome size (**Fig. 3e**). Despite the differences in abundance, the total exonic content of retrocopies and unprocessed pseudogenes (typically 1–8 Mbp), was at least an order of magnitude lower than the total protein coding exonic content (27.8-149.2 Mbp; **Extended Data Fig 5a**), with mammals showing the greatest lineage-specific variation (**Extended Data Fig. 5b**).

While most retrocopies decay rapidly, a minority is retained over deep evolutionary distances as functional retrogenes^84,85^. To quantify this balance across vertebrates, we compared species-specific and conserved retrocopies per lineage and found that ∼50% of retrocopies in mammals, birds, and turtles were species-specific, compared to ∼70% to 98% in all other lineages (**Extended Data Fig. 5c**). These results suggest that mammals, birds, and turtles retain a substantially larger fraction of ancestral, lineage-shared retrocopies, while other clades are dominated by recent and/or poorly retained insertions. Across lineages, most retrocopies (∼45-90%) lacked intact ORFs (**Extended Data Fig. 5d**), presumably reflecting rapid mutational decay and truncation inherent to the retroduplication process^86^. Of the remaining retrocopies (∼10-50%) that retained coding potential, a fraction (∼15-35%) displayed significant signatures of selection (conservative dN/dS > 1.2, positive selection; FDR corrected p-value < 0.05 per retrocopy; **Extended Data Fig. 5d**). Ray-finned fishes, birds, and amphibians exhibited the highest proportions of retrocopies under selection, consistent with elevated rates of retrogene recruitment in these lineages. Species-specific retrocopies in birds, squamates (e.g. lizards), amphibians, and fishes, exhibited lower dN/dS values, which clustered below 0.5 (**Extended Data Fig. 5e**), consistent with purifying selection. In agreement with previous reports^84,87,88^, mammals showed similar distributions of dN/dS values between conserved and species-specific retrocopies. Together, these results illustrate that pseudogenes (in particular, retrocopies) constitute a major, lineage-specific feature of vertebrate genome diversity. Mammals emerge as outliers both in retrocopy abundance and in long-term retention, whereas other clades exhibit younger, selection-shaped retrocopy landscapes consistent with active retrogene recruitment. More details are in **Supplementary Information section 3**

### Repeat content

The higher completeness of VGP genomes provides the opportunity to investigate previously inaccessible regions of the genomes. It also provides the opportunity for improvement and benchmarking of repeat annotation tools, such as EDTA^89^ for transposable elements (Ou et al., to be submitted to bioRxiv). We performed RepeatModeler v2.04^90^ annotation on the assemblies, which identifies all types of endogenous repeats, and found that repeats explained much of the variation observed in genome size within each major clade, and some of the differences across clades (**Fig. 3f**). K-mer analyses revealed a concordant phylogenetic signal, with amphibians, sharks/rays and the early deep branches of ray-finned fishes showing the highest 31-mer repeat content, and birds showing the least (**Fig. 3g**, **Supplementary Table 14**). Phylogenetic independent contrasts analysis revealed that this relationship is dependent on genome size, which has a stronger correlation in the VGP genomes relative to non-VGP assemblies (**Extended Data Fig. 6a**). There were outlier genomes with large deviations from the predicted value (**Extended Data Fig. 6a,b**), which we found have repeat structures that are substantially different from others from the same lineage. More details are in **Supplementary Information section 4**.

Satellite annotation yielded 36,216 high-confidence arrays grouped into 3,429 recurrent families across 577 species in the Cactus alignment (**Extended Data Fig. 6c**). These satellites were overwhelmingly lineage gap-flanking fields specific (94% restricted to a single species, the NCBI order being the deepest rank at which any complex satellite was shared beyond the telomeric repeat). Monomer lengths clustered at integer multiples of the nucleosome repeat (**Extended Data Fig. 6c, dashed lines**), with a predominant cluster at ∼171 bp. More details are in **Supplementary Information section 5**.

For telomere repeat analysis, PacBio CLR-based assemblies showed shorter telomeres than PacBio HiFi-based assemblies (**Fig. 2i**), likely reflecting technological limitations (Medico et al., to be submitted to bioRxiv). Thus, for telomere analyses, we focused on HiFi based assemblies and treated lengths as lower-bound estimates owing to the maximum length of HiFi reads (∼20 kbp). Considering the clades where we had 10 or more species, we noted a wide range of telomere sizes within each, with mammals showing the widest range (∼1-25 kbp; **Fig. 3h**, **Supplementary Table 15**). Nevertheless, there were some overall lineage-specific differences, where turtles had on average longer telomeres (14 kbp on average), whereas birds, lizards, ray-finned fishes and deuterostome invertebrates had on average the shortest (6-7 kbp on average).

Although ribosomal DNA loci are complex and difficult to assemble completely^20,91^, we were able to annotate rDNA clusters in most species, as detailed in a companion analysis^92^. Across the broad taxonomic scope of the VGP dataset, and in line with previous findings^93,94^, the 18S, 5.8S, and 28S rRNA-encoding genes exhibited relatively low sequence divergence and strong coupling between sequence and secondary structure compared with the internal transcribed spacers (**Fig. 3i**, **Supplementary Table 16**). The spacers showed elevated sequence divergence, pronounced heterogeneity across lineages, and substantially weaker sequence–structure constraint. The 18S and 5.8S rRNA gene lengths were highly conserved across vertebrate lineages, whereas 28S showed substantially greater variation, suggesting lineage-specific expansion and contraction of variable domains superimposed on an otherwise strongly constrained rRNA core. Interspecies differences extend to higher levels of organization, including extensive variation across vertebrates in rDNA chromosomal distribution, repeat copy number, and GC content^92^. More details are in **Supplementary Information section 6**.

### Regulatory elements across vertebrates

#### Cis-regulatory modules

Cis-regulatory elements represent critical functional components of genomes. These include promoters of genes, which are often GC-rich and missing from short-read based assemblies^1^. We applied a cross-vertebrate regulatory annotation pipeline, the <u>V</u>ertebrate <u>R</u>egulatory <u>MO</u>dule <u>D</u>etector (VRMOD^95^, Gonçalves et al., under review) to selected VGP Phase I genomes. VRMOD is designed to predict cis-regulatory modules (CRM) directly from genomic sequence without requiring any training data or parameter tuning for vertebrate genomes (Gonçalves et al., under review). VRMOD predicted 280,622 to 12,028,231 CRMs per genome, with an average CRM length ranging from 192 to 324 bp (**Extended Data Fig. 7a**, **Supplementary Table 17**). These elements collectively covered 33.5% to 68.2% of each genome (**Extended Data Fig. 7b**). Although these estimates are substantially larger than the fraction of the genome occupied by protein-coding exons (4-8%), they are consistent with that occupied by introns and UTRs (20-60%) and with previous studies suggesting that regulatory sequences constitute a substantial portion (43%) of a human genome^96^, based on a comprehensive CRM prediction framework. Our finding indicates that regulatory elements might be more widespread throughout the genome than previously recognized. There were clear lineage and species differences, and this was best explained by a strong positive correlation between total CRM length and genome size (**Extended Data Fig. 7c**; Pearson’s correlation coefficient r = 0.95; P-value <0.01). However, whether this reflects increased regulatory complexity or is driven primarily by genome expansion through repetitive and other non-coding elements remains an open question. Collectively, VRMOD-predicted CRMs provide a high-resolution regulatory annotation set for each genome, establishing a universal framework for comparative regulatory genomics across vertebrates.

### Epigenetic promoter regulation

We also leveraged HiFi read-based methylation profiling to examine DNA methylation patterns at promoters. We studied the subset of 88 species that were sequenced with HiFi long-read technology, had methylation calls made during sequencing, and had RefSeq-annotated genomes in the data freeze as of April 2025 (**Supplementary Table 18**), encompassing representatives of all major vertebrate clades^97^. In the methylation profiles, we found a canonical promoter signal of hypomethylation starting around 1,000-3,000 bp on either side of the transcription start site (**Fig. 4a**). There were lineage differences, where lobe-finned fishes and amphibians had the highest methylation levels, and birds and sharks, the lowest, around promoters. Non-avian reptiles had the widest promoter profiles, whereas amphibians had the narrowest. These results provide a comparative baseline of epigenetic promoter regulation across vertebrates.

**Fig. 4.**
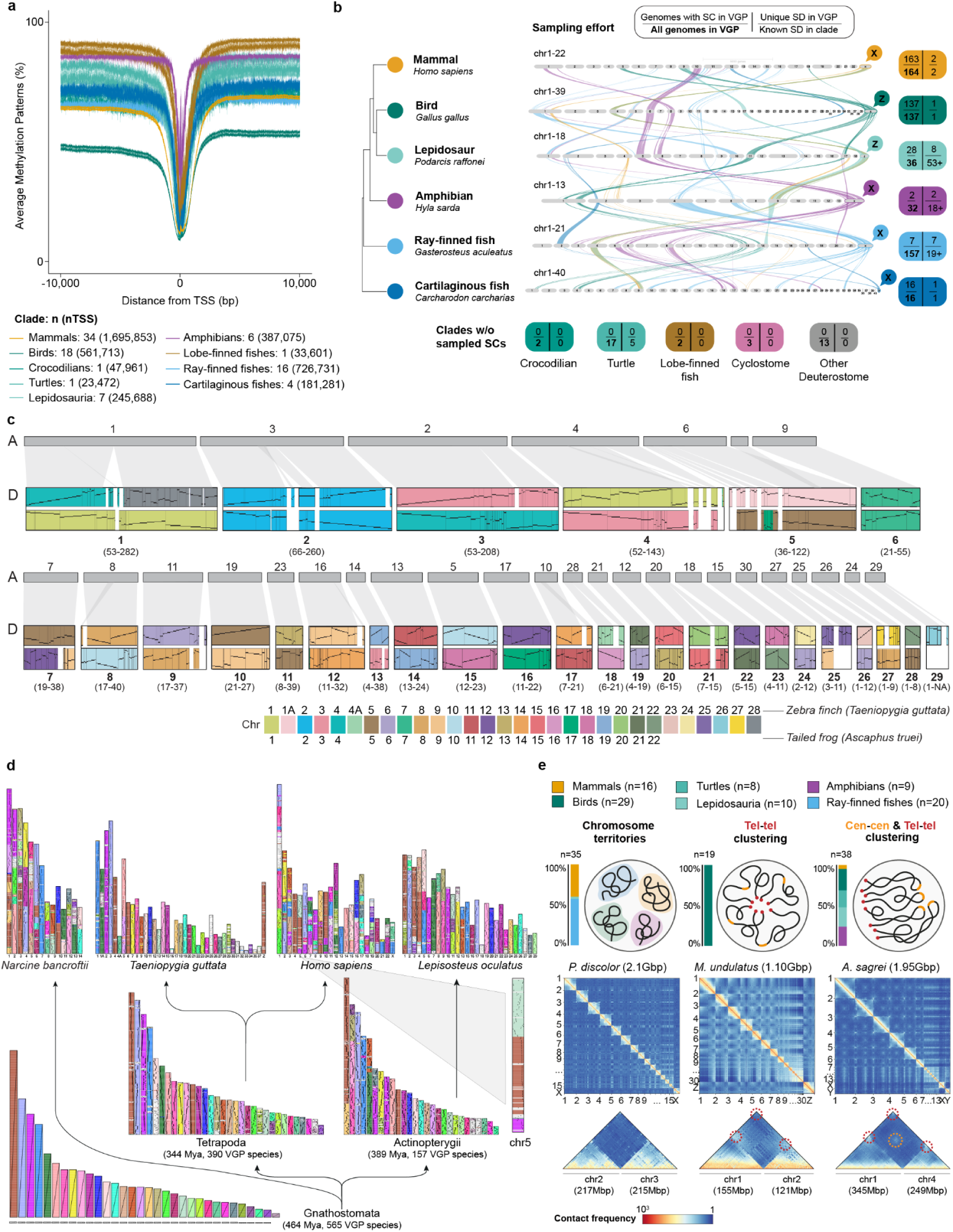
Chromosome organization and evolution. **a**, DNA methylation patterns on potential promoters assessed at all protein coding genes, centralized at transcript start sites, across vertebrate clades. Values are the total number of species (n) and total transcription start sites (nTSS) for each calde. **b**, Synteny of sex chromosomes (SC) across representative species from six extended lineages demonstrated that sex chromosomes are independently recruited from multiple independent syntenic blocks across the genome. The colors of the river plot track the genomic location of syntenic sequence from the sex chromosome in one representative of an extended lineage across the genomes of the other representatives. The boxes show four values as indicated in the upper figure panel legend, for each clade (color). **c**, Consensus reconstruction of the ancestral tetrapod karyotype between (A) ancestral blocks reconstructed by AGORA and (D) by DESCHRAMBLER. Grey ribbons connect AGORA blocks to DESCHRAMBLER reconstructed ancestral chromosome fragments (RACFs). Merger of DESCHRAMBLER RACFs and AGORA blocks yield 29 ancestral Tetrapoda chromosomes based on synteny between two or more of the three reconstructions. The two independent colored tracks of reconstructed Tetrapoda chromosomes are painted according to zebra finch (top) and Coastal tailed frog (bottom) reference chromosomes used in DESCHRAMBLER-based reconstructions. The numerical range in () indicates chromosome sizes (Mbp) based on the zebra finch and Coastal tailed frog DESCHRAMBLER reconstructions. Vertical lines within blocks designate homologous synteny with two divergent reference genomes; horizontal lines show orientation. White intervals between colored blocks demarcate boundaries between RACFs. The color code at the bottom was developed to represent chromosomes that have the same number in the zebra finch and the Coastal tailed frog. In the absence of one-to-one correspondence, additional colors are used. **d**, The Gnathostomata vertebrate ancestor inferred chromosomes reconstructed by AGORA using BUSCO genes and their fates in several descendant vertebrate lineages, including two intermediate descendant genomes (Tetrapoda, Actinopterygii) and four genomes representing major extant vertebrate classes: cartilaginous fishes (electric ray), birds (zebra finch), mammals (human), and ray-finned fishes (spotted gar). Colors show ancestral Gnathostomata chromosomes. Light gray, genes that belong to non-assembled fragments of the Gnathostomata reconstruction. Black dots, positions of BUSCO genes, with diagonals denoting stretches of genes whose order was conserved since the most-recent common ancestor (i.e., syntenic blocks). Chromosomes are ordered by decreasing size. **e**, Hi-C contact maps of chromosomal interactions. Top panel, cartoon of chromosome interaction types and their proportions (left bar plots) across mammals, birds, turtles, lepidosauria, amphibians, and ray-finned fishes. Middle and lower panels, Hi-C genome wide interactions (middle panels) and pair of chromosomes (lower panels) for three species representing their vertebrate lineages: *Phyllostomus discolor* (bats), *Melopsittacus undulatus* (birds), and *Anolis sagrei* (anolis lizards). Red circles represent interactions between telomeres, whereas the orange line segment indicates interactions between centromeres (Rabl-like configurations^116^). Genome and chromosome sizes are specified.

### Genome evolution across vertebrates

#### Sex chromosome diversity

The VGP phase I dataset represents the largest collection of assembled vertebrate sex chromosomes to date, with broad sampling across many vertebrate lineages that have known sex determination systems (**Supplementary Table 1, Columns T and F**). This sampling includes all mammalian and avian orders, representing conserved XX/XY and ZZ/ZW systems, respectively, as well as representatives of other sex determination systems including turtles (ancestral temperature dependent system), cartilaginous fishes (XX/XY system)^98,99^, and lacertid lizards (a shared ZZ/ZW system)^100^. Further, we assembled the genomes of many species with one or more additional sex chromosomes, including platypus and echidna (5X’s and 5Y’s or 4Y’s, respectively), *Artibeus* bats (X, Y1 and Y2), European common lizard (Z1, Z2, and W), the trahira fish *Hoplias malabaricus* (X1, X2, Y) and the Atlantic stingray (X1, X2, and Y) (**Supplementary Table 1, Column T**). To validate sex chromosome assignments and identify new sex chromosomes, we implemented multiple complementary approaches on VGP the VGP Phase I dataset. One tool was HalfDeep, which uses genome-wide sequencing-depth to infer regions divergent enough within the assembly to preclude long-read mapping from both haplotypes, resulting in a haploid-level (or approximately one-half) sequencing depth (Gable et al., to be submitted to bioRxiv). Under this prediction, heterogametic individuals (XY or ZW) should show haploid-level sequencing depth for both of their sex chromosomes, whereas homogametic individuals (XX or ZZ) should retain diploid-level sequencing depth. This is useful for both confirming genotypes of the reference individual in species with known sex chromosome systems and for implicating previously uncharacterized sex chromosomes in additional taxa.

We find that many VGP genomes with chromosomal sex determination fit these predictions (**Extended Data Fig. 8a-g**) and that assembled sex chromosomes were more complete and larger when using HiFi data and parental trio or Hi-C phasing relative to CLR data and/or partial phasing (**Extended Data Fig. 8h,i**). For sex chromosomes that are not highly diverged, typical expectations about depth may not apply, so we further applied SCINKD, which uses differences in k-mers between assembled haplotypes to identify sex chromosomes(<u>Pinto et al. 2026</u>). We found two cases of potentially novel sex chromosomes (**Supplementary Table 1**): the spotted gar (*Lepisosteus oculatus*) and the West Atlantic trumpetfish (*Aulostomus maculatus*). These genomes also contributed to testing the origin of genetic sex determination in Chelidae (see ^101^ Across all approaches, we corrected seven cases of sample sex mislabeling, six cases of incorrect sex chromosome assignment, and ten instances of unassigned sex chromosomes where the system was known from existing literature; we also detected 48 instances of incomplete or incorrect sex chromosome assembly (collapsed ampliconic signals between Y/W chromosomes; **Extended Data Fig. 8a**).

We next characterized orthology of conserved linkage groups found in any sex chromosome with any chromosome of any other species in the VGP dataset. Using synteny analyses, we found that the VGP 579 species dataset has 21 inferred independently evolved sex chromosome systems across five major vertebrate clades (mammals, birds, within lizards, within turtles, within ray-finned fishes, and cartilaginous fishes), recruited from multiple unique autosomal regions, using chicken chromosomes as the reference (**Fig 4b**; **Extended Data Fig. 8i, Supplementary Table 19**). Historically, analyses of vertebrate sex chromosome orthology were conducted relative to chicken chromosomes as a proxy for ancestral linkage groups^102^. Comparing the 21 independent sex determination systems identified in the VGP dataset, we observed regions within 29 of 39 chicken chromosomes are homologs of sex chromosomes in at least one of the VGP-sampled chromosomal sex determination systems (**Extended Data Figure 8i**). We chose representative species from each extended lineage with chromosomal sex determination in the VGP dataset to exemplify this across whole genomes, tracking the autosomal location of sex chromosomes across species, and highlighting the lability of the genome to be recruited as a sex chromosome (**Fig.4b**). More expansive analyses of sex chromosome evolution and best practices for sex chromosome assembly and annotation are presented in a companion study (Gable et al., to be submitted to bioRxiv). Additional companion studies use the VGP Phase I assemblies to investigate sex chromosome organization, evolution, and neo formation in birds (Shakya et al., submitted; Jackson et al. to be submitted to bioRxiv), an anuran^103^, a species of North Atlantic rockfish (*Sebastes*)^104^, and in cartilaginous fishes^98^. The lessons learned from the VGP Phase I genome sequencing effort highlight the need for more thorough interrogation of sex chromosomes during the assembly process, using multiple lines of evidence, facilitating novel insights in sex chromosome evolution.

### Ancestral vertebrate genome reconstructions

The combination of high-quality assemblies and species breadth across the vertebrate tree for VGP Phase I enables inference of chromosome evolution in deep evolutionary time. We reconstructed the chromosomal organization of vertebrate ancestors using two complementary approaches, AGORA^105^ and DESCHRAMBLER^106^ (**Supplementary Table 20**). AGORA reconstructs ancestral gene order along a phylogeny by inferring ancestral gene adjacencies (syntenies) from the distribution of extant gene adjacencies across modern genomes of extant species. DESCHRAMBLER adds finer spatial resolution for conserved coding and non-coding sequences by using multiple pairwise whole-genome alignments of extant genomes to a reference genome, first defining conserved syntenic blocks and then inferring their ancestral arrangement in the common ancestor.

Independent AGORA and DESCHRAMBLER reconstructions of the ancestral tetrapod genome (344 MYA) yielded largely concordant findings with 29 consensus chromosomes in the common ancestor of all vertebrates (**Fig. 4c**), providing high confidence in the karyotypes of the deep reconstructed ancestors. It was not possible to reconstruct the gnathostome (jawed vertebrate) ancestor with DESCHRAMBLER due to the extensive sequence divergence between fishes and other vertebrate clades. Resolving the karyotype of more ancient vertebrate ancestors will likely require additional highly conserved non-vertebrate outgroups.

To maximize AGORA’s power, we mapped a common set of 3,695 BUSCO genes (odb12.1, including all 3,390 genes of the vertebrata_odb12 BUSCO set plus 305 genes of the mammalia_odb12 BUSCO set so as to cover avian dot chromosomes (which are absent from the vertebrata BUSCO set) on all 579 genomes across the VGP phylogeny. Using these genes as homologous genomic landmarks, we were able to reconstruct ancestral genomes for nearly all vertebrate ancestors, including Tetrapoda (344 MYA), Euteleostomi (424 MYA) and Gnathostomata (464 MYA) using AGORA (**Fig. 4d**; **Supplementary Table 24** used as time-calibrated tree), providing firm support for ancestral karyotypes which had previously been difficult to ascertain^107,108^. Gnathostomata represent 99.8% of all vertebrates. In addition, AGORA reconstructed the most likely gene order along these ancestral chromosomes, enabled by the unprecedented breadth and quality of VGP genomes.

Analysis of ancestral and extant karyotypes revealed that birds and some early-diverging ray-finned fish lineages (e.g., spotted gar, *Lepisosteus oculatus*) have retained more of the ancestral karyotype, whereas teleosts, mammals, and amphibians have experienced more rearrangements, both intra- and inter-chromosomal (**Fig. 4d** and data not shown), confirming previous observations^105,108,109^. Notably, the newly resolved ‘dot’ chromosomes of birds^23,110^ were likely present in very ancient ancestors, with some traced as far back as the ancestor of all jawed vertebrates (Gnathostomata, 464 MYA). All AGORA reconstructions are available for download and analyses at a dedicated Genomicus server^111,112^. Further analyses of AGORA- and DESCHRAMBLER-reconstructed ancestral genomes are reported in a companion study (Giudicelli et al., in preparation).

### 3D genome architecture

A major advantage of the VGP data set is that Hi-C data, which delineate 3D sequence contacts, are available for all species. Besides enabling both scaffolding and haplotype phasing for genome assembly, they enable the investigation of 3D chromosomal structure. We analyzed the Hi-C data in a subset of 92 species that had sufficient sequence coverage and were representative of the major phylogroups (**Supplementary Table 21**), using HiCExplorer v3.7^113^ to visualize normalized contact maps with genomewide interactions. We found three broad, distinct chromosome arrangements that were clade enriched: (i) within-chromosome clustering as chromosome territories, mainly found in placental mammals (n = 14), ray-finned fishes (n = 20) and one non-avian reptiles within Lepidosauria (n=1); (ii) clustering of telomeres, as found in birds (n = 19); and (iii) chromosomal clustering at both centromeres and telomeres, as found in marsupials (n = 2), some birds (n = 10), non-avian reptiles including turtles (n = 8) and Lepidosauria (n = 9), and amphibians (n = 9) (**Fig. 4e**, **Supplementary Table 21**). The present results expand on patterns previously reported for representative species of vertebrates^114,115^, and appear to be independent of tissue type (**Supplementary Table 21**). Our findings suggest that the 3D-genome architecture of vertebrates is an evolvable trait that exhibits not only conserved constraints but also plasticity across taxa. The lineage-specific architecture conservation provides compelling evidence that vertebrate genomes exhibit deeply divergent, lineage-dependent principles of nuclear organization, and establish a new evolutionary framework for understanding how chromosome architecture has diversified in parallel with genome evolution. A more detailed exploration of these patterns is presented in a companion study (Marin-Gual et al., to be submitted to bioRxiv).

### Gene family evolution

#### Protein-coding gene family evolution

A major driver of vertebrate molecular evolution, adaptation, and innovation is gene duplication – a process fuelled both by single gene duplication events and by whole genome duplication (WGD). Early vertebrate evolution was characterised by two rounds of WGD, *i.e.* the 2R hypothesis^117^ (one shared by all vertebrates and one shared by all gnathostomes) and a third round in the ancestor of teleost fish (3R), a large subgroup of ray-finned fish^118^. In total, we found 6,189 retained protein coding gene duplicates (formed by either WGD or single gene duplication events) that map to the branch where the 3R event occurred (**Fig. 5a**). However, retained paralogs were not uniquely concentrated to branches where the WGD occurred, but rather were a pervasive feature throughout vertebrate evolution (gene duplications per branch: median 562, lower quartile 158, upper quartile 1,380; **Fig. 5a**, **Supplementary Table 22**). The stem bird lineage had a high number (1,468) of duplications that were retained in extant species (a pattern that cannot be attributed to a WGD event as there is no WGD event hypothesized on this branch) and their functions were enriched in various biological processes, including antimicrobial immunity, oxygen transport, and chemotaxis (**Supplementary Table 23**). In contrast, loss of gene families across vertebrates was less variable, with no notable outliers associated with evolutionary innovations or radiations (gene duplications per branch: median 314, lower quartile 199, upper quartile 548) (**Fig. 5a vs 5b**).

**Fig. 5.**
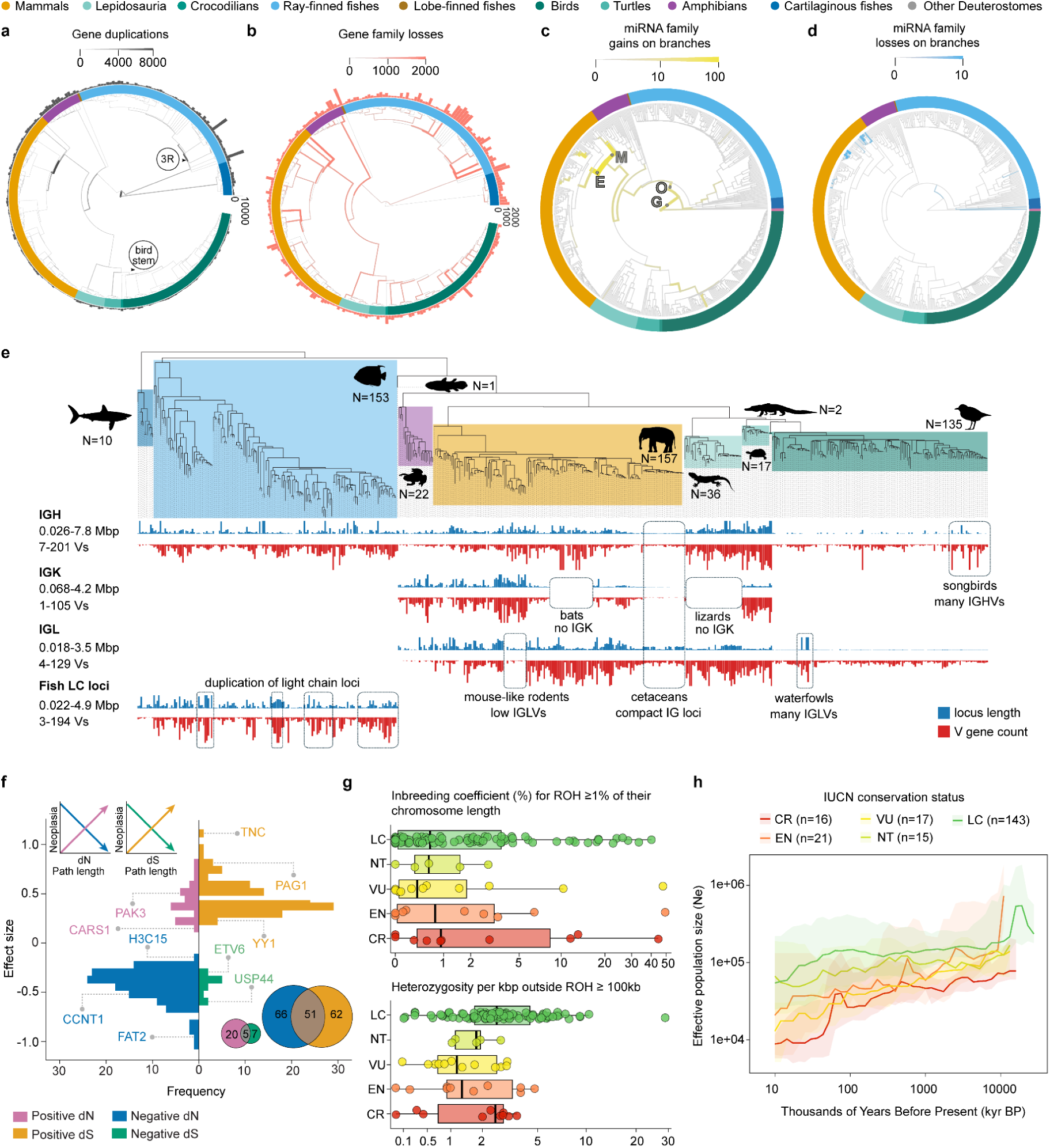
Gene family evolution and conservation. **a**, Protein-coding gene family duplications across 243 RefSeq annotated vertebrate genomes. The outer ring plots duplications on terminal branches only; duplications are indicated by the thickness of the branch (Tree based on ROADIES phylogeny). Position of the 3R WGD event is labelled. **b**, Protein-coding gene family losses across the same 243 genomes. Outer ring, losses on terminal branches only, displayed by the thickness of the branch. For gene family losses, data are not included for the two basal branches (stem Chondrichthyes and stem Osteichthyes). **c**, miRNA gene family gains across 554 vertebrate genomes. Branches where major expansions occur are labelled accordingly: “G” = Gnathostomata, “O” = Osteichthyes, “E” = Eutheria, and “M” = Metatheria. **d**, miRNA gene family losses per branch across the same 554 genomes. **e**, Comparative analysis of immunoglobulin loci in jawed vertebrate species. For each species in the ROADIES tree, locus lengths and counts of immunoglobulin (IG) variable (V) genes are shown as blue and red bars, respectively, for IGH, IGK, IGL, and fish light chain loci (from the top to the bottom). Left, ranges (2.5 and 97.5 percentiles) of locus lengths and V gene counts for each locus type. Dashed rectangles, highlighted lineage-specific immunogenomics features. **f**, The number and effect sizes of genes with significant associations between neoplasia prevalence and either dN or dS, with colors indicating direction of effect. **g**, Box plots of ROH and heterozygosity values for each species, categorized by IUCN status. **h**, PMSC plots of effective population sizes over time, for the median and interquartile range of species in each IUCN category: LC, least concern; NT, near threatened; VU, vulnerable; EN, endangered; CR, critically endangered.

### MicroRNA gene family gains and losses

We reconstructed the birth and death rates of microRNA (miRNA) families across 553 VGP Phase I assemblies which identified lineage-specific expansions and losses (**Supplementary Tables 24-26);** 28 species were omitted from this analysis due to technical difficulty in miRNA annotation (**Supplementary Table 25**). A total of 404 unique miRNA families were delineated across vertebrates, 37 of which were retained by all extant vertebrate species (for entire miRNA gene family presence/absence matrix see **Supplementary Table 25)**. Strikingly, many miRNA gain events coincided with major evolutionary transitions in vertebrates (**Fig. 5c**), which highlights their role as potential drivers of innovation: e.g., 12 miRNA families arose in the most recent common ancestor of Gnathostomata and an additional nine in the ancestor of Osteichthyes. However, as reported previously^119–122^, the largest expansion of the miRNA gene repertoire occurred in the ancestors of Metatheria and Eutheria, with 52 and 93 novel miRNA families, respectively. Substantial miRNA gene expansion also occurred within primates, with 31 novel miRNA families arising in the ancestor of Simiiformes (Simian primates), in agreement with previous reports^123^, and 21 additional families arising in the ancestor of catarrhines (Old World monkeys and apes)^124^. Finally, we observed nine novel miRNA gene families that arose in the ancestor of Murinae (Old-World rats and mice), concordant with previous reports^125^. Death of miRNA gene families was also observed: 290 miRNA families were completely lost in at least one taxon, with the average number of loss events per miRNA gene family being 7.91. As expected, miRNA loss events were concentrated at terminal branches (**Fig. 5d**). Further analyses that demonstrate the role of miRNAs in the convergent evolution of diverse placental phenotypes in mammals are presented in a companion manuscript (Fenn et al., to be submitted to bioRxiv).

### Immunoglobulin loci across vertebrates

A striking example of the gene birth-and-death dynamics is found in antibody encoding immunoglobulin (IG) loci, representing one of the major branches of adaptive immunity common for vertebrate species^126^. These loci include immunoglobulin heavy chain (IGH) and light chain (IGK: kappa, IGL: lambda, IGI: iota, IGS: sigma) loci that were previously reported among the most dynamic and complex regions in vertebrate genomes^127^, likely as a consequence of their coevolution with pathogens^128^. IG loci generate enormous antibody diversity through somatic recombination of variable (V), diversity (D), joining (J), and constant (C) genes. The highly repetitive nature of immunoglobulin loci made them difficult to analyse comprehensively using short-read assemblies. We used the VGP Phase I data to perform the first comparative analysis of complete sequences of immunoglobulin loci across all families of jawed vertebrates. We focused on immunoglobulin variable genes, as they provided the clearest signal of locus span and organization.

The IgDetective tool^129^ was modified such that newly identified immunoglobulin variable genes could be used to discover even more new genes in an iterative manner (**Extended Data Fig. 9a**). Across the VGP 579 species dataset, we detected 495 intact IGH loci that were specific to one or more vertebrate clades (**Fig. 5e**, **Supplementary Table 27**). We also detected 164 intact IGK and 354 intact IGL loci in 364 species. Due to incomplete and conflicting chain labels of known variable genes, light chain types were not distinguished in both ray-finned fishes and cartilaginous fishes and are thus collectively referred to as fish light chain loci. The IGK locus was lost independently in bats, lizards, and birds (**Fig. 5e**). In contrast, the IGL locus was missing or highly reduced in amphibians and mouse-like rodents (Myomorpha). There may be some compensatory mechanism for these losses, given that all chains are essential for viable antibody production in humans^130^. Among mammals, cetaceans (whales and dolphins) possessed exceptionally compact immunoglobulin loci with a reduced number of immunoglobulin genes across all chain types. This reduction may reflect decreased exposure to the pathogens that their terrestrial ancestors faced. While most birds have compact immunoglobulin loci as well, consistent with their generally compact genomes^131^, waterfowl (Anseriformes) and songbirds (Passeriformes) have a greater number of IGL and IGH variable genes, respectively (**Fig. 5e**). On average, waterfowls encode 43 IGLV genes per species compared with 18 in other birds, whereas songbirds encode 86 IGHV genes on average compared with 32 in other birds. We speculate that the expansion of the germline gene set in songbirds and waterfowl may be related to the fact that they are often hosts/vectors for pathogens and parasites^132^.

Consistent with previous studies^133,134^, duplications of IGH loci were detected for 14 Yangochiroptera bat species and light chain loci for 56 ray-finned fishes scattered across 36 taxonomic families (**Extended Data Fig. 9b**). Most IG loci were found on single contigs (**Extended Data Fig. 9b**), indicative of complete locus assemblies. In birds, however, IGH loci were found on short contigs (**Fig. 5e, Extended Data Fig. 9b**), likely due to the location of IGH loci on bird dot chromosomes, which are associated with known PacBio HiFi sequencing dropout (**Extended Data Fig. 9c**; **Supplementary Information section 7**)^23^. We observed mosaic patterns of duplications of light chains across ray-finned fish lineages (**Extended Data Fig. 9b**). Pairwise alignment of subsampled variable genes clarifies the evolution of these loci. For example, some light chain loci of ray-finned fishes exhibited similarity to both IGK or IGL, suggesting that the modern light chain loci emerged from distinct ancestral copies in the early evolution of vertebrates (**Extended Data Fig. 9d**). Although the broad pattern is consistent with previous studies^135,136^, the structure of variable gene similarity points to the presence of additional, previously unrecognized light chain subtypes. Finally, IG loci were not detected for salamander species; our iterative IgDetective procedure reported more than 10,000 V gene candidates, but these did not pass manual curation and did not form continuous loci. This finding will be addressed in follow up studies using transcriptomic data collected from expressed antibody repertoires. Overall, the higher quality assemblies allowed a greater analysis of more full-length immunoglobulin loci, for reconstructing the evolutionary history of these genes. Further comparative genomic studies of duplicated gene families are in companion papers, including olfactory receptor genes in birds^137^.

### Comparative genomics of traits and diseases

#### Specialized and fundamental vertebrate traits

Relatively complete assemblies, broad phylogenetic sampling, and whole-genome alignments in the VGP Phase I dataset provide an opportunity to revisit candidate genetic changes associated with specialized or fundamental vertebrate traits. As examples, we focused on one specialized trait, vocal learning, used for speech in humans, and a fundamental trait, the vertebrate body-plan specification. We used both the 577-way Progressive Cactus alignment and 579 species all-vs-all alignment-depth profiles.

For vocal learning, we first revisited FOXP2, a transcription factor extensively studied in relation to speech in humans^138^ and song in vocal learning birds^139^. FOXP2 has two specialized amino acid substitutions, originally proposed to be unique to humans^140^. In the reference-free 577-way Progressive Cactus alignment, the complete gapless FOXP2 locus, including exons, introns, and surrounding regulatory regions, was present for nearly all Phase I vertebrate species. For the first FOXP2 substitution, a threonine to asparagine (T303N), we confirmed that the human N amino-acid state was not found in any other primate, mammal, bird, amphibian, Cartilaginous fish, or non-avian reptile (except for one lizard, blue-tongue_skink, *Tiliqua scincoides*). However, the N substitution was found in most ray-finned fishes, excluding salmonids and close relatives (**Extended Data Fig. 10a**, **Supplementary Table 28**). For the second FOXP2 substitution, asparagine to serine (N325S), the human S substitution was found, as expected in some bats, mainly all Yinpterochiroptera, and all Canivora sequenced, and unexpectedly, in three amphibian species and three ray-finned fishes (**Extended Data Fig. 10a, Supplementary Table 28**). When we examined the surrounding four amino acids for the two sites, they were also highly conserved across vertebrates, except for ray-finned fishes (**Extended Data Fig. 10b, Supplementary Table 29**), indicating relaxation in this clad for this region. No species had both human substitutions, and thus the combination of the two substitutions together remains unique to humans. Analysis of protein coding genes in the all-vs-all alignment across 579 species, reveals that FOXP2 amino acid sequence conservation is among the top 5% of conserved proteins, and the top two genes on human chromosome 7 (measured by alignment depth; **Extended Data Fig. 10c**). In contrast, the surrounding regulatory regions and intervening introns which have LINE elements are more permissive to transposon insertion (**Extended Data Fig. 10e**).

The NOVA1 splicing regulator contains an isoleucine-to-valine substitution (I197V) reported to be unique to humans in a highly conserved region of the protein and proposed to be associated with language evolution; mice humanized for the V substitution showed differential splicing of genes involved in vocalization behavior and produced ultrasonic courtship songs with more varied syllable modulation^141^. The V substitution has also not been found in archaic humans (Neanderthal or Denisovan genomes)^141^. Like FOXP2, in the VGP Phase I progressive cactus alignment of the complete gene, we confirmed that no other primates, mammals, birds, or non-avian reptiles had the human V amino-acid substitution. However, the V substitution was present in all frogs and all teleost ray-finned fishes but not other amphibians or other fishes (**Extended Data Fig. 10a, Supplementary Table 29**); the teleost V substitution arose from a different codon (GTG) than in frogs and humans (GTA), suggesting convergence between teleost and humans. The surrounding four amino acid sequence was 100% conserved across all vertebrate species examined (**Extended Data Fig. 10b, Supplementary Table 30**). Together, the FOXP2 and NOVA1 results show that although the individual variants are not unique to humans, their combination is unique and in regions with resistance to change. Future investigations can determine if the rare FOXP2 variants in Canivora (which contain vocal learning pinnipeds) and Yinpterochiroptera bats (also vocal learners), and NOVA1 in frogs and teleost fishes, have some influence on their vocal communication behavior.

Beyond nucleotide level variation, we also applied the long read VGP genomes to study the large set of regulatory elements found in human parvalbumin positive (PV+) inhibitory interneurons, whose predicted open chromatin levels show some convergence among vocal learning mammals (humans, bats, cetaceans, and pinnipeds )^142^. Given their functional association with vocal learning, like the single nucleotide findings in FOXP1 and NOVA1, we predicted that these PV-interneuron associated enhancers might be in genomic regions of high synteny conservation to maintain the enhancer-gene relationship. We found the PV+ interneuron open chromatin regions were in regions of higher synteny in mammals, birds and fish (**Extended Data Figure 10f,g Supplementary Table 31**), suggesting that despite their enrichment in vocal learners, these enhancers maintain important gene regulatory functions across distantly-related species. Further details are in **Supplementary Information section 8**.

The other highly conserved locus on human chromosome 7 was the HOXA cluster (**Extended Data Fig. 10c,d**), a cluster of homeobox genes that regulate anterior-posterior axis and limb patterning during embryonic development^143^. The anterior-posterior axis of the vertebral column is a defining feature of vertebrates, and HOX genes play central roles in body-plan specification. The HOXA cluster was nearly completely assembled across all 579 species. Across the cluster, we observed pronounced depletion of transposable elements, including SINEs, LINEs, DNA transposons, and LTR classes (**Extended Data Fig. 10c,d**). This pattern is consistent with strong purifying selection acting on both the coding and cis-regulatory architecture of the cluster, likely reflecting the requirement for precise spatiotemporal collinear expression during development. Together, these analyses illustrate the value of the VGP Phase I assemblies and alignments for investigating candidate loci for specialized and fundamental traits across deeply diverged vertebrate lineages.

### Cancer genetic dynamics in vertebrates

We next asked whether the VGP genomes could be leveraged to investigate disease-relevant evolutionary dynamics. As an example, we examined possible mechanisms of cancer suppression across vertebrates. We integrated three curated datasets of cancer associated genes, COSMIC^144^, IntOGen^145^, and OncoKB^146^ (all downloaded in June 2025) to produce a list of 1,169 unique genes that are known to be implicated in human disease. For each gene, we constructed multiple sequence alignments (MSAs) across 43 bird and 66 mammal species with paired cancer prevalence data for each species. Using the MSAs, we applied the adaptive branch-site random effect likelihood aBSREL^147^ model in HyPhy to estimate the total amount of synonymous (dS) and non-synonymous (dN) evolution for each gene in each species, measured as the distance from root-to-tip. We then fitted a phylogenetic Poisson regression in which the neoplasia prevalence^148^ (the sum of benign and malignant tumours) was modelled as a function of the number of necropsies, body size, dN, and dS for each species. Across the 1,169 genes analysed, 149 showed a significant association between neoplasia prevalence and dN, while another 149 showed a significant association with dS (**Fig. 5f**, **Supplementary Table 32**). As expected, most genes (125 out of 149) exhibited a negative association between neoplasia prevalence and dN. That is, natural selection acted on a larger number of genes to decrease tumour prevalence. In contrast, the majority of genes (145 out of 149) had a positive association between neoplasia prevalence and dS. That is, high rates of ‘nearly neutral’ evolution^149^ through time were associated with increased tumour prevalence. Given the overlap, we noted many genes with a significant positive association between neoplasia prevalence and dS also showed a significant negative association with dN (**Fig. 5f**). This pattern suggests a ‘tug-of-war’ in which natural selection counteracts background molecular evolution to suppress tumour development across the vertebrate tree of life. Further analyses of cancer-associated evolutionary dynamics are presented in a companion manuscript (Butler et al., to be submitted to bioRxiv). Complementary analyses of longevity are presented in companion studies, including identifying a specific regulatory region in the GDF11 gene associated with the repeated evolution of low heart rate and long lifespan in mammals (Li et al., in preparation); and extreme longevity and potential cancer-resistance mechanisms in the Greenland shark and little sleeper shark^150^.

### Species conservation

#### Single nucleotide and structural diversity

Reference genomes are valuable resources in conservation biology^151^. Of the 579 data freeze species, 24% are classified as threatened by the IUCN Red List (**Supplementary Table 2, Column AJ**). The highly contiguous haplotype-resolved assemblies allow for the accurate identification of both Single Nucleotide Variants (SNVs), enabling inference of demographic history using Sequentially Markovian Coalescent (SMC) methods, and Structural Variants (SVs), which are challenging to ascertain with short reads. Most variance in SNV heterozygosity was found within taxonomic groups, with no statistically significant differences among groups (using a phylogenetic generalized least square model with taxonomic group and IUCN conservation status as explanatory variables followed by a Tukey’s post-hoc test; a Bonferroni-corrected p-value threshold of 0.05 was used to determine statistical significance; **Extended Data Fig. 11a left panel**, **Supplementary Table 33**). By contrast, SV heterozygosity was significantly higher in jawless fishes, ray-finned fishes, amphibians, and squamates, than in birds and mammals (**Extended Data Fig. 11b**). Both SNV and SV heterozygosity had a weak correlation with conservation status according to IUCN categories, with species of least concern having significantly higher heterozygosity than critically endangered species (**Extended Data Fig. 11a right panel, 11b**). Taken together, these results highlight the utility of high-quality genome assemblies in characterizing previously intractable genetic diversity and providing insight into the diversity of threatened taxa. More detailed analyses are presented in a companion manuscript (Lou et al., to be submitted to bioRxiv).

### Runs of homozygosity and demographic inferences

We further calculated the fractions of the genome that are in long runs of homozygosity (ROH; ≥1% the length of the chromosome), as these are indicative of recent inbreeding within an individual’s lineage^152–154^. We found that species with higher extinction risk (as defined by IUCN Red List status) had higher proportions of long ROH in their autosomes (**Fig. 5g**, **Supplementary Table 34**; β = 0.029, Odds Ratio = 1.03, SE = 0.013, *p*-value = 0.027). Some Critically Endangered species had much higher proportions of long ROH, supporting that recent inbreeding plays an important role in extinction risk. There was a positive skew of outliers for Least Concern species that had high proportions of long ROH; this could be due to the result of life history traits for certain species, or a currently unrecognized level of species extinction threat^155^. Heterozygosity outside of these ROH tracks was lower in threatened species (β = –0.479, Odds Ratio = 0.52, SE = 0.119, *p*-value < 0.001). After correcting for phylogenetic relationships, only heterozygosity outside ROH retained a significant relationship with IUCN status (posterior mean = -0.328, CI = -0.54 to -0.125, *p*-value = 0.002), while the inbreeding coefficient (F_ROH_) was not significant (posterior mean = 0.025, CI = 0.002 to 0.052, *p*-value = 0.054). This is consistent with what has previously been reported by Zoonomia^156,157^. Interestingly, a companion manuscript reports no correlation between ROHs and hatching failure rates or other fitness components, likely due to historically small population size, in the Critically Endangered Hawaiian crow (*Corvus hawaiiensis*)^158^, highlighting the role of demographic history in shaping inbreeding load and species’ susceptibility to inbreeding depression^154^.

In keeping with these analyses, demographic reconstruction using PSMC^159^ shows that threatened species tend to have had lower historical effective population sizes as far back as >10 MYA (**Fig. 5h**), indicating that lower long-term historical genetic diversity may predispose species to higher extinction risk. The median effective population sizes of species across all IUCN categories have gradually declined over time, with Critically Endangered species exhibiting a sharp decline around 60 thousand years ago, coinciding with human demic expansion^160^, again supportive of some species having a predisposition to human impact^157^. Additional companion studies look at these factors in further detail (Gardiner et al., in preparation), and apply VGP reference resources to population- and conservation-genomic analyses of leatherback turtles (*Dermochelys coriacea*)^161^, captive mainland clouded leopards (*Neofelis nebulosa*) (Breech et al., in preparation), and southern black rhinoceroses (*Diceros bicornis*) (Antonets et al., in preparation).

## Discussion

This study reflects a 10-year effort of developing methods to produce high-quality reference genomes, applying them over 8 of those years on species representing a wide breadth of vertebrate biodiversity at the ordinal level and divergence times between 50-100 MYA, and utilizing those genomes to address principled questions that were otherwise not be possible or more difficult with draft-quality genomes and less phylogenetic breadth. With this genomic resource, we gained a better understanding of vertebrate protein-coding and regulatory region evolution, repeats, specialized and fundamental traits, disease, and conservation.

### Reflections

Our species selection criterion tried to balance standard taxonomic classification hierarchies usually based on morphology, with phylogeny and divergence times derived from fossil and genomic sequence data^26^. These two frameworks are not always concordant and there is ongoing debate regarding which better reflects species relationships^162^. Applying both approaches provided a practical compromise while achieving a more even coverage of species divergences in the same time window and the diverse Earth climate conditions at the time. Amphibian species were sampled less densely within the 50-100 MYA divergence window, due to the large genome sizes that would cause additional technical limitations and sequencing cost. Nevertheless, we sampled within the two major frog groups (Archaeobatrachia and Neobatrachia), the salamanders, and caecilians sufficiently to allow for more balanced analyses with other lineages. Unlike past studies^15,37^, future efforts will be able to ask questions about similarities and differences in species divergences that occurred soon before and soon after the last mass extinction, around the K-Pg boundary, using this more uniform genomic dataset.

Our finding that analytical power improved as genome quality increased over the 8-year course of the project, for example, through more accurate detection of telomeric sequences in PacBio HiFi assemblies compared with earlier CLR-based assemblies, highlights the need to consider potential technical biases arising from differences in assembly quality within the Phase I dataset^1,28^. Nevertheless, because most assemblies achieved the high-quality standards established by the VGP, the resulting gains in analytical accuracy and biological insight are orders of magnitude greater than those attainable from large-scale efforts based primarily on draft-quality short-read-based genomes^17,18,156,163^. As genome assemblies continue to become more complete and error-free, technical biases will be further reduced, enabling increasingly robust comparative genomic analyses^3,20,164^.

One of the most enabling advances made possible by the structural quality of the VGP Phase I dataset was the reconstruction of the chromosome organization of the vertebrate common ancestor, as well as the tracing of how those chromosomes evolved in subsequent diverse lineages^105,165^. It is intriguing that bird genomes retain the closest organization to the common ancestor of vertebrates, given that birds are essentially a highly derived dinosaur lineage of reptiles^131,165^. This pattern suggests that the common ancestor of birds and other reptiles had such an ancestral organization. The inferred common ancestor vertebrate genome also provides a framework for tracing the chromosomal history of most genes and genomic regions among vertebrates.

At the other end of the genomic scale, the nucleotide quality and completeness of the VGP Phase I dataset enable higher resolution and accuracy of gene variants that may have contributed to human-specific traits, such as FOXP2 and NOVA1 associated with speech evolution^141,166^. They also allow detection of variants inferred to be resistant to cancer and other diseases. Such findings generate more informed hypotheses for functional testing in species with these rare variants.

From a conservation perspective, the VGP Phase I resource helps preserve the genomic legacy of about 500 million years of vertebrate evolution. Some currently threatened species carry genomic signatures of long-term vulnerability that predate human impacts by millions of years^157^. Others, especially monotypic and ditypic species, may harbor genomic features associated with long-term persistence through climatic instability, glacial-interglacial cycles, and even mass extinctions. The Living Planet Report 2024 documented an average 73% decline in monitored vertebrate population sizes between 1970 and 2020, which our PMSC genomic analyses supports, indicating that it is imperative to proceed with sequencing future high-quality genomes before further biodiversity is lost.

### Phase II

The lessons from Phase I are both scientific and organizational (Paez et al., in preparation). What worked well was the grassroots, global, collaborative model that allowed many institutions, taxonomic experts, sequencing centers, and analysis groups to contribute to a shared vertebrate genomics resource^1,8,27^. At the same time, several challenges became also apparent, including difficulties in obtaining samples from some key lineages with limited numbers of species, incomplete resolution of certain genomic regions in the non-T2T assemblies, uneven annotation data, large and repetitive genomes, and the challenge of scaling the highest assembly standards across the full breadth of vertebrate diversity. These lessons provide a foundation for improving both the scope and execution of Phase II.

The Phase I resource represents ∼1% of named vertebrate species (∼800 of ∼70,000). For Phase II, the current VGP target family list includes approximately 1,100 vertebrate families based on the NCBI taxonomy. Of these, 263 family-level lineages are already represented by completed Phase I ordinal lineage genomes, and deeper within orders another 136 family-level lineages have been completed. Thus, the 816 completed Phase I species currently cover approximately 393 Phase II family-level lineages, or ∼36% of the Phase II target. The target number will likely increase when selecting family level divergences, expected to be at the pre 30 MYA time frame, if using bird and mammal family divergences as a guide^15,26,37^. At the same time, the 50–100 MYA divergence interval remains incompletely sampled. In addition, some extant lineages still lack the nuclear genomic data necessary for divergence-time estimation and phylogenetic placement, while published estimates often differ depending on taxon sampling, sequence content, and inference method^15,167^. The VGP phylogeny group will therefore need to refine which lineages diverged before, during, or after this interval, and to identify branches of the vertebrate tree that remain underrepresented.

We suggest that VGP Phase II should also strive to oversample acquisition and quality bottlenecks, develop scalable T2T and near-T2T protocols at scale, improving transcriptomic and regulatory annotation, integrating trait and conservation databases, and connecting vertebrate resources more tightly with invertebrate efforts and the broader Earth BioGenome Project^5,27^.

The most persistent assembly gaps remain concentrated in repetitive regions, including centromeres, telomeres, and rDNA arrays^20,91,168,169^. Other repetitive regions, such as immune loci, transposable elements, and segmental duplications, are substantially better assembled in the VGP assemblies than in previous resources^17,18,170^. The current T2T genome protocols can fill and correct many of the remaining repetitive regions, with the important exception of rDNA arrays, but they are currently most tractable for genomes at the smaller end of the vertebrate range, roughly 0.5–3.0 Gbp^20,59,60,171^. Larger and more repetitive genomes, such as salamanders (10-60 Gbp), will require further methodological development to improve quality, to generate polyploid assemblies, T2T, or near-T2T assemblies at scale^172^. We therefore recommend that Phase II develop or adapt semi-automated, lineage-aware protocols capable of producing T2T or near-T2T assemblies for thousands of genomes per year.

Annotation remains a future challenge. The VGP Phase I effort did not enforce generation of RNA transcriptome data for all species, an essential ingredient for high-quality annotations^4^. For Phase II, we recommend that greater emphasis and investment be put into generating long-read transcriptome data of each species sequence, for high-quality gene annotation^173^. Additional epigenetic methylation data that now comes from the genome sequence long reads^97,174^ we believe should be used for annotation routinely. Greater emphasis should be placed on non-coding annotation, because many major vertebrate differences are likely to lie in introns, intergenic regions, repetitive regions, and regulatory elements rather than protein-coding sequence alone^166,175,176^.

As Phase II progressively adds family-level representatives, and as population-level resequencing accumulates, alignment graphs can become the natural scaffold for a vertebrate pangenome^177–179^. This will be especially powerful if coupled to a curated trait database that links genomic variation to morphology, physiology, behavior, ecology, disease resistance, and conservation status^157,176,180^.

More complete assemblies and broader phylogenetic sampling reduce ambiguity relative to previous comparative genomic studies, by improving orthology, resolving repetitive and structurally complex regions, and distinguishing human-specific, lineage-specific, and convergent changes^1,17,18,20,23,66^.

In conclusion, the overall value of the VGP resource is that it makes biological inferences more precise, and better contextualized. The VGP Phase I shows how high-quality genome assemblies generated at a broad phylogenetic scale can strengthen comparative biology by improving confidence in genome-wide analyses of vertebrate evolution, trait innovation, genome organization, and biodiversity conservation. VGP Phase 1 assemblies have already been used in over 200 studies in the past (**Supplementary Table 35**), and the entire collection here for an expected over 100 studies^181^. Phase I is rapidly allowing novel biological findings not possible with prior draft-quality assemblies and more limited phylogenetic breadth. Our study also highlights the lineages, genomic regions, annotations, and analytical frameworks that should be prioritized next. Phase II should therefore not only add more species, but also build on the Phase I foundation to develop a scalable, increasingly complete, and biologically integrated vertebrate pangenomic resource for understanding, preserving, and restoring the diversity of vertebrate life.

## Methods

### Sampling

The BioSample page in NCBI for each genome accession (**Supplementary Table 1**) contains information on source of tissue, tissue or cell culture type, preservation method, and persons responsible for collecting the sample. Only samples collected, preserved, and transferred with procedures that ensure limited cellular and nucleic acid degradation were used. Most samples were flash frozen with liquid nitrogen or dry ice immediately after collection, transported on dry ice, and stored at -80°C when possible. For some samples, in addition to long term -80°C storage, 95-100% ethanol preservation was used, especially for cases of nucleated blood sampling. When possible, we avoided samples preserved in RNA (RNA later) or DNA (DNA guard) preservative agents, as we found that this causes degradation of chromatin contacts for Hi-C sequencing and leads to partial degradation of DNA, negatively impacting the advantage of long reads^182^. Samples were collected, shipped, and received in compliance with local and global permitting.

### Genome sequencing and assembly

Genome assemblies included in the VGP Phase I resource were generated over an 8-year period during which sequencing technologies and assembly algorithms improved substantially. Accordingly, individual assemblies were produced using combinations of sequencing data types and assembly workflows appropriate to the available technologies and the characteristics of each species’ genome. The principal assembly categories and their progression from partially phased long-read assemblies to fully phased and telomere-to-telomere (T2T) diploid assemblies are described in the Results and summarized for individual species in **Supplementary Tables 1 and 2**. Sequencing failures (**Supplementary Table 5**) often resulted from improperly frozen samples, unintentional thawing during transport or border-control permit processing, preservation in RNAlater or other nucleic acid stabilization cell lysis reagents^182^, or insufficient tissue or cellular material to yield the required quantity of DNA. Across assembly types, long reads provided the primary sequence information for contig construction, while Hi-C data were used for chromosome-scale scaffolding^56,57^ and, in later workflows, for haplotype phasing. Earlier VGP assemblies were generated primarily from PacBio continuous long reads (CLR), using FALCON/FALCON-Unzip^183^ to generate contigs, Hi-C with the SALSA algorithm and Bionano optical maps^184^ to scaffold the contigs using, and Illumina short reads for polishing and/or phasing^1^. With the adoption of PacBio High Fidelity (HiFi) reads^185^, most subsequent assemblies were generated using the hifiasm algorithm^29,64^, with haplotypes resolved using either parental short-read data^61^ or Hi-C data. T2T or near-T2T assemblies used thes same data types and additionally incorporated Oxford Nanopore Technologies ultra-long reads, and were generated using workflows including Verkko or updated hifiasm-based approaches algorithms^20,22,23,59,67^.

Reproducible implementations of the principal VGP assembly and evaluation workflows are available through the Galaxy platform^28^ (https://vgp.usegalaxy.org/; Lariviere et al, to be submitted to bioRxiv) and in the Sanger NextFlow toolkits (https://pipelines.tol.sanger.ac.uk/pipelines). All assemblies were manually curated to identify and correct structural assembly errors, assign sequences to chromosomes (including sex chromosomes when present), and remove contaminating sequences of other species when present^186^ (Lariviere et al, to be submitted to bioRxiv). To select a reference genome for each species, we selected the highest-quality haplotype meeting or exceeding VGP quality criteria, moved in the sex chromosome from the other haplotype in the cases of a heterogametic sex and fully diploid assembly, and moved in the mitochondrial genome when available^72,187^; designated this genome set as the “main” haplotype. Accession numbers for the main and additional haplotypes, mitochondrial genome accessions, sex-chromosome representation, assembly technology, annotation status, and summary assembly metrics are provided in **Supplementary Table 1** for the 579 species data freeze and **Supplementary Table 2** for the entire 816 species dataset. Additional sequencing and assembly-associated metadata, including publicly deposited sequence records and assembly accessions, are available through NCBI under the VGP umbrella BioProject PRJNA489243 and affiliated project records. Raw sequencing data, intermediate assembly products and associated project resources are additionally available through GenomeArk (https://www.genomeark.org/) for a subset of genomes generated by members of VGP proper. For genome assemblies deposited in GenBank or RefSeq, NCBI Foreign Contamination Screen reports provide an additional public record of screening for foreign biological and synthetic sequence contamination^188^.

### Genome annotation

Genome assemblies were annotated using the NCBI RefSeq Eukaryotic Genome Annotation Pipeline (EGAP)^4^ or its publicly available implementation, EGAPx (Tvedte et al, in preparation). For each species, publicly available short and long RNA-seq datasets were retrieved from the Sequence Read Archive (SRA)^189^ and selected to maximize tissue representation, with the aim of capturing a broad range of transcriptional diversity. When RNA-seq data were not available at the species level, datasets from closely related species within the same genus or family were used. RNA-seq reads along with well-annotated proteins from related taxa were aligned to the corresponding genome assemblies and incorporated to support transcript reconstruction, definition of exon–intron boundaries, and refinement of gene structures. When applicable, EGAPx was executed using parameters consistent with the standard EGAP workflow to ensure comparability across annotations.

Our gene annotations cover ∼97.9% (567) of the VGP Phase I data freeze assemblies and were generated with available or newly generated transcriptome data including, at a minimum, RNA-seq datasets from congeneric or even confamilial species. Thirteen genomes were annotated using only proteome evidence. Annotation lift-over or other gene prediction methods based on related species are only a partial solution and thus not employed, because sequence divergence over 40–500 million years is too large to recover lineage-specific gene models, isoforms, and regulatory elements with sufficient accuracy. The type of annotation and annotation summary results is in the annotation report under each genome accession (**Supplementary Tables 1 and 2**).

### Repeat masking

RepeatModeler version 2.0.4/December 2022 was used to generate RepeatMasker family libraries. The assemblies were repeat-masked with RepeatMasker versions 4.1.4 family libraries for earlier obtained assemblies and version 4.1.7-p1 for later obtained assemblies.

### Genome quality comparisons

We downloaded one genome assembly per BioSample^190^ entry using the “BioSample Representative==primary” filter in the Genomes on a Tree (GoaT) ^191^ portal (https://goat.genomehubs.org/) to compare the VGP data-freeze genomes to all vertebrate genomes in GenBank, removing those tagged as “atypical” by GenBank (i.e., too large/short, highly contaminated, suppressed). For each genome, we calculated the gene completeness by mapping the proteins from the BUSCO vertebrata_odb12 database using miniprot v0.14-r265 with arguments --trans -u -I --outs=0.95 and then ran the “analyse” module of compleasm^192^ v0.2.7 with argument --retrocopy. We furthermore calculated the Nx values (size of smallest sequence such that the sum of all sequences larger and equal to this value make up x% of the genome) for contigs and scaffolds for each genome in 0.1% increments using the script “get_nx.py”. The mean and median Nx value for each increment was then plotted for each lineage within the VGP 579 data-freeze genomes and other genomes within GenBank. To create a list of genomes for direct comparison, used in the kmer spectra analysis below, we searched for genomes within the GenBank list with the same species taxID and not part of the VGP data-freeze. For instances where multiple genomes were available, we selected the genome with the largest contig N50 that was below 1 Mbp. Dedicated scripts are available in the associated Github repository (https://github.com/VGP/vgp-assembly-manuscript).

### Karyotype and scaffotype analyses

We compared cytogenetically derived chromosome numbers with chromosomes curated in VGP assemblies using a subset of species from the primary dataset. The subset included species for which both (i) curated chromosome-level assemblies were available at the time of the VGP 579 species data freeze and (ii) cytology-based chromosome counts with karyotype images in major databases or the literature. When multiple values or population-level variation were reported, records were verified through targeted literature review. All karyotype data was stored in the Genomes on a Tree (GoaT)^191^ database and used for the analysis. The curated dataset is provided in **Supplementary Table 6** and can also be found with a dedicated query for the corresponding subset in GoaT (https://goat.genomehubs.org/projects/vgp_phase1). For the assemblies, we utilized those that followed VGP and Earth BioGenome Project (EBP) assembly standards, with all identified sex chromosomes included in the primary haplotype. In several amphibian assemblies, large chromosomes were split into multiple scaffolds during submission. Assembly chromosome counts (scaffotypes) were normalized to diploid for our comparative analysis. The diploid number was calculated as twice the haploid chromosome count represented in the primary haplotype, adjusted for additional chromosome-scale scaffolds introduced during submission (for example, sex chromosomes in heterogametic individuals or chromosomes split into multiple scaffolds owing to size constraints). For assemblies in which sex chromosomes could not be identified, the diploid count was estimated as twice the number of chromosome-level curated scaffolds. For visualization, inferred diploid counts were compared with cytogenetic data. When multiple cytogenetic values were available, the value matching the assembly-derived count was used for plotting.

### Telomere annotation

Telomeres were annotated with Teloscope v0.1.3 (https://github.com/vgl-hub/teloscope; Medico et al., to be submitted to bioRxiv) using the parameters -k 50 -d 200 -l 500 -y 0.5 -x 1 -w 200 -s 200 -g -r -e -m -i. The annotation was restricted to assembled chromosome-level curated scaffolds, e.g. excluding unassigned contigs. Each assembly was scanned against a library of 38 hexamers, including the vertebrate canonical motif TTAGGG, its 18 single-substitution variants (allow to differ from canonical by 1 bp), and their 19 reverse complements (Medico et al i preparation). Species with additional canonical motifs were handled on a case-by-case basis. For each chromosome end, we retained the longest telomeric block on the expected strand (CCCTAA-rich on the p-arm, TTAGGG-rich on the q-arm) and classified it as terminal when its most-distal base lay within 1 kbp of the scaffold end. Per-assembly telomere completeness was computed as the observed-over-expected ratio: O/E% = 100 × (terminal ends passing filters) / (2 × haploid chromosome number). Telomere length was defined as the length of each retained terminal block. To reduce the influence of extreme outliers, each assembly was summarized by its median terminal telomere length across vertebrate lineages.

### Reconstruction of a preliminary species tree using ROADIES

To enable multiple whole-genome alignments using CACTUC that requires a guide phylogenetic tree and other downstream analyses, we used ROADIES^75^ version 0.1.10 to infer a preliminary species tree from a 579-species VGP dataset (comprising main assemblies only). Given the scale and deep timescale of the VGP species tree, we performed per-group gene subsampling, leading to 123,141 gene trees and filtering of inadequate loci (see Supplementary information). For better placements at deep nodes, the position of invertebrates and several other species was provided as a constraint tree. We note that this tree is designed to support preliminary analyses rather than present a comprehensive resolution of vertebrate relationships.

### Retrocopy identification

Retrocopies were identified using a customized implementation of the RCPedia framework^193^. mRNA sequences from annotated protein-coding genes were extracted using gffread^194^ and aligned to the corresponding genome assemblies with LAST^195^, an aligner selected for its sensitivity to divergent sequences, permitting long intronless matches consistent with retrotransposition events. Alignments shorter than 120 bp - approximately the exon length for some species - were discarded. Candidates located within 200 kbp of their parental gene locus were excluded to minimize contamination from segmental duplications. To confirm a reverse transcription origin, candidate retrocopies were required to exhibit at least one continuous alignment spanning two consecutive exons of the parental gene (i. e., ungapped alignment crossing an exon-exon junction from the 3′ end of the parental mRNA). Candidates in which ≥40% of the aligned sequence overlapped regions annotated by RepeatMasker, simple repeats, or WindowMasker were filtered to remove alignments with excessively low complexity or repetitive content. Retrocopies derived from the same parental gene were required to be separated by at least 500 kbp, and candidates overlapping three or more annotated exons (i.e., two or more annotated introns) were excluded. Retrocopies originating from highly redundant gene families (>5 paralogous copies) were removed to minimize ambiguity in parental gene assignment. For each retained retrocopy locus, the most likely source transcript was selected based on a combined evaluation of sequence identity and alignment coverage, with ties resolved by maximizing the relative alignment score. Fragmented alignments corresponding to the same retrocopy were merged if separated by gaps of up to 6 kbp, a threshold chosen to accommodate interruptions (gaps in alignment) by transposable element insertions, thereby enabling reconstruction of full-length retrocopy structures.

### Orthology inference of retrocopies

Orthology relationships among retrocopies were inferred using an all-by-all pairwise alignment strategy based on genomic sequence similarity and reciprocal best-hit criteria. For each species, genomic coordinates of the curated set of retrocopies were extracted, and each locus was extended by 3 kbp upstream and 3 kbp downstream to include flanking regions. These extended regions were retrieved from the corresponding reference genome assemblies using bedtools getfasta^196^. The inclusion of a flanking sequence improves alignment robustness while preserving a defined retrocopy core for downstream filtering. Pairwise sequence alignments were performed using LASTZ^197^ in general output format. Each retrocopy from the focal species was aligned against the complete set of retrocopy sequences from each downstream species in the species phylogenetic order. Alignments were run in parallel on a high-performance computing environment to maximize throughput. Raw LASTZ outputs were post-processed using a custom Python script. Briefly, alignment blocks were parsed, and block sizes were computed as the aligned length on the query sequence. Only alignment blocks overlapping the retrocopy core region (defined as the sequence excluding the 3 kbp flanks) by at least 50% in both query and target were retained. For each query-target pair, retained blocks were merged and summarized to obtain the total aligned length, weighted mean nucleotide identity, and coverage relative to both query and target sequences. Pairs were retained if they met minimum thresholds of 50% nucleotide identity and at least 50% coverage of both the query and target retrocopies and their flanking sequences. Coverage was computed independently for query and target as the fraction of the sequence spanned by retained alignment blocks. To assign putative orthologous retrocopies, a reciprocal best-hit strategy was applied. For each query retrocopy, the best target hit was defined as the alignment maximizing a composite score based on nucleotide identity and query coverage. Conversely, for each target retrocopy, the best query hit was selected based on nucleotide identity and target coverage. Retrocopy pairs that were mutual best hits under these criteria were retained as orthologous relationships. Filtered reciprocal best-hit pairs from all pairwise species comparisons were aggregated to generate a non-redundant list of orthologous retrocopy relationships across species.

### dN/dS analysis and tests of selection on retrocopies

Selective constraints acting on retrocopies were evaluated using pairwise dN/dS (ω) analyses between each retrocopy and its inferred parental gene. For each species, predicted retrocopy open reading frames (ORFs) were obtained from TransDecoder (https://github.com/TransDecoder/TransDecoder) analyses of retrocopy nucleotide sequences. Only the longest ORF per retrocopy was retained. Metadata tables were constructed linking each retrocopy ORF to its corresponding parental transcript and protein, based on previously curated retrocopy annotations and NCBI feature tables. Amino acid sequences for retrocopy ORFs and their parental proteins were extracted and paired into two-sequence FASTA files, with one entry corresponding to the retrocopy and the other to the parental gene. In parallel, the corresponding nucleotide sequences were retrieved for both retrocopies and parental transcripts. For each retrocopy-parental pair, matched amino acid and nucleotide FASTA files were generated to enable codon-aware alignment. Protein sequences were aligned using ClustalW^198^. Resulting amino acid alignments were converted into codon-based nucleotide alignments using PAL2NAL^199^, preserving reading frame information and excluding codons containing gaps. These codon alignments were formatted for compatibility with PAML and constituted the input for dN/dS estimation. Rates of nonsynonymous (dN) and synonymous (dS) substitutions, as well as their ratio (ω = dN/dS), were estimated using the codeml program from the PAML package^200^. For each retrocopy–parental pair, two models were fitted: (i) a free ω model allowing ω to vary, and (ii) a neutral model with ω fixed to 1. Log-likelihood values were extracted from codeml outputs, and likelihood ratio tests (LRTs) were performed by comparing the two models. Statistical significance was assessed using a chi-squared distribution with one degree of freedom. Retrocopies were classified as evolving under purifying selection when ω < 0.5, and under positive selection when ω > 1.2; the latter threshold was adopted conservatively to reduce false positives arising from stochastic variation in ω estimates for short or recently emerged ORFs. A retrocopy was considered to show a significant signature of selection when its LRT was significant (FDR-corrected p < 0.05) and its ω satisfied one of these thresholds. For each species, codeml outputs were parsed to extract ω, dN, dS, and log-likelihood values. Likelihood ratio statistics (ΔlnL) and corresponding P values were computed for all retrocopy-parental pairs. To ensure robust inference, results were filtered to exclude cases with unreliable synonymous divergence (dS < 0.05 or dS > 3) or extreme ω estimates (ω > 5). P values were corrected for multiple testing using the Benjamini–Hochberg false discovery rate (FDR) procedure. Filtered dN/dS results were integrated with orthology assignments to classify retrocopies as conserved (orthologous across species) or lineage-specific. When multiple ORFs were predicted for a given retrocopy, results were collapsed at the retrocopy level to avoid redundancy, retaining evidence of selection if any ORF showed a significant signal. Retrocopies were further annotated by clade using a curated species classification table. All analyses were performed using consistent parameter settings across species to ensure comparability of dN/dS estimates and selection tests.

### Cis-regulatory module annotation

Cis-regulatory modules (CRMs) were annotated using the Vertebrate Regulatory MOdule Detector (VRMOD^95^; Gonçalves et al., in revision), a sequence-based regulatory annotation pipeline. CRMs were defined broadly as DNA elements with potential roles in regulating spatiotemporal gene expression, including enhancers, silencers, insulators, locus control regions, promoters, and RNA post-transcriptional regulatory modules. VRMOD was applied to selected VGP genomes from birds, Lepidosauria, ray-finned fishes, turtles, and mammals. Predicted CRM coordinates were summarized per genome to calculate the number of CRMs, mean CRM length, total CRM length, and the fraction of each genome covered by predicted CRMs. Correlations between total CRM length and genome size were assessed for the VGP genome sets using Pearson’s correlation coefficient. Summary statistics and comparative results are reported in **Extended Data Fig. 7** and **Supplementary Table 17**.

### miRNA gene family gains and losses

MirMachine 2.0^201^ (https://github.com/sinanugur/MirMachine) was used to annotate miRNA precursor sequences from 554 species genomic FASTA files^202^. A small number of species were omitted from the analysis due to technical issues (**Supplementary Table 25**). The covariance models (CMs) in MirMachine 2.0 were trained specifically to prevent bias from well-annotated genomes, such as human, thereby reducing the likelihood that inferred expansions are artifacts of the miRNA detection method. Bitscore-filtered annotation files were collated and converted to miRNA gene family presence/absence matrix in R (**Supplementary Tables 12,13**). To perform Dollo parsimony-based reconstruction of miRNA family gain and loss, a time-calibrated version of the VGP phylogeny was created. Divergence times at 20 nodes^203^ were used as calibration points for the chronos function of the R package ape^204^, supplemented by an additional 270 divergence time points taken from TimeTree^26^ providing a time-calibrated version of the VGP Phase I guide tree (**Supplementary Table 11**). The divergence estimate for Cyclostomi from other vertebrates was excluded due to overlap with predicted divergence times for Vertebrata causing issues with time calibration.

For each miRNA family, gain and loss across the vertebrate tree were estimated from the miRNA presence/absence matrix via Dollo parsimony using the *mapDolloChanges* function of the R package Claddis^205–207^, performing 100 runs per miRNA family. The node of gene birth and node(s) of gene loss (where appropriate) supported by the plurality of these 100 runs were extracted as the best-supported estimates for each miRNA family.

### Protein coding gene family evolution

The longest canonical protein sequence corresponding to each gene was selected from all RefSeq-annotated genomes in the VGP at the time of the 579 species data freeze (*i.e.* 243 genomes total annotated at the time). The generated FASTA files feature one protein per gene and were used as input to the OrthoFinder pipeline (version 2.5.5)^208^ with Diamond-Blast^209^ for sequence similarity searching and Dendroblast^210^ for tree generation. Duplications inferred by the reconciliation step of the pipeline were extracted by parsing the “Duplications.tsv” file and were plotted on the tree inferred for these species in this publication. The branch leading to modern birds displayed a particularly high level of paralog retention. To determine if there were gene functions enriched in this set we used the *Gallus gallus* homologs of genes that duplicated in the ancestor of all birds. The Duplications.tsv file was scoured for gene families duplicated on this branch, taking *G. gallus* genes found on either side of these duplications as the foreground set to test against the background of all *G. gallus* genes. The Gene Ontology resource was used to perform the enrichment analysis using PANTHER classification^211^, searching for biological process terms overrepresented in the foreground set, i.e. a PANTHER Overrepresentation Test (Released 2024-08-07). The annotation version and release data was GO Ontology database DOI: 10.5281/zenodo.15066566 Released 2025-03-16. The reference list was “*Gallus gallus”* (all genes in the database). The annotation dataset was “GO biological process complete”. The test type was Fisher’s Exact and the correction applied was False Discovery Rate.

To quantify protein coding gene family loss events across the vertebrate phylogeny, we parsed Orthofinder output in a Dollo-like parsimony approach where gene families could be gained once and lost multiple times. For each orthogroup (OG), the number of genes present in each species tree node was compared. The deepest node where the parent featured more genes than either of the child nodes was then considered the origination node for that OG. Gene family losses were then inferred by propagating presence downward. That is, at each branch, if the child count is zero the OG is lost and that lineage is not followed further for that OG. We note that without further outgroups, gene families lost on either of the branches emerging from the root are undetectable according to this method.

### Analyses of k-mer repeat spectrum

We used Jellyfish^212^ to compute 31-mer frequencies for each of the 568 ‘vertebrate’ genomes included in the ROADIES tree, and the 248 comparator existing genomes from the same species (see "Genome Quality Comparison" above). We used RESPECT^213^ to summarise the results. Let *r_i_* be the number of 31-mers observed 1 ≤ *i* ≤ *m* times in a genome, and note that the assembly length is 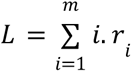. We compute the percentage of the genome composed of k-mers that appear at least twice (i.e., 1 − *r*_1_/*L*) and refer to it as the 31-mer repeat ratio (RR). We performed maximum likelihood ancestral state reconstructions of RR and assembly length using a Brownian motion model with the phytools R package v2.4-4^214^ based on the ROADIES tree. We used the same tree, restricted to the 248 species available with existing genomes, to compute the same quantities using existing and VGP genomes. We visualize the relative increase in RR from existing genomes to VGP genomes. Values higher than 0 indicate VGP genomes are more repetitive.

We also computed phylogenetic independent contrasts (PIC)^215^ using the ape R package v5.8-1^204^ to compare changes in RR and assembly length at internal nodes of the tree. A PIC value close to 0 indicates little difference between the two descendant lineages, while a large positive or negative value indicates a substantial change according to phylogenetic correction. We then compared PICs of RR to PICs of assembly and observed that nodes with a higher contrast in RR also tend to have a higher contrast in assembly length. We quantified the strength of this relationship using Spearman correlation, selected to be robust to the effect of several outliers. Outliers from this trend are indicated on the figure. Additionally, we use the ape package^204^ to compute a phylogenetic covariance matrix under a Brownian model^215,216^ and use it to fit a linear model by generalized least squares as implemented in the nlme R package v3.1-168 ^217^. Using the learned regression, we can assign an expected assembly length for each genome based solely on its RR and compare the expected length to the actual assembly length.

### Single nucleotide variant (SNV) and structural variant (SV) calling and PSMC analysis

To analyze SNVs and SVs segregating between two haplotypes within a single individual, we started from all VGP assemblies of the 579 species data set that are haplotype-resolved. We filtered the dataset for assembly contiguity by requiring both haplotypes to have N50 values above 100 kbp. We excluded invertebrate species from this analysis. In addition, we excluded potential incomplete/unbalanced assemblies by requiring the shorter haplotype to be at least 70% the length of the longer assembly. With a final dataset of 476 haplotype-resolved assemblies, we first aligned the alternate assembly to the primary assembly using minimap2-v2.29 with -a -x asm5 --cs -r2k^218^. When the designation of primary vs. alternate assemblies is not specified (e.g. when they are labeled paternal vs. maternal), we used the assembly with the higher contig N50 as the primary. We sorted and indexed the resulting sam files with samtools-v1.17^219^. We called structural variants with svim-asm-v1.0.3^220^. In addition, we converted the sam files to paf format and called SNVs, insertions, and deletions using the paftools script in minimap2 with -l 2000 -L 10000. SNV heterozygosity was calculated as the number of SNVs in each individual divided by the length of the primary assembly that paftools considers callable. Insertion and deletion heterozygosity is calculated as the number of insertions and deletions larger than or equal to 50 bp called by paftools divided by the callable length. Since paftools does not call inversions and duplications, their heterozygosity was calculated as the number of inversions and duplications larger than or equal to 50 bp called by svim-asm, divided by the total length of the primary assembly. SV heterozygosity was then obtained by taking the sum of the heterozygosity in insertions, deletions, inversions, and duplications.

The IUCN conservation status for each species was obtained with the IUCN API^221,222^, and the phylogenetic tree was obtained from the Open Tree of Life^223^. Comparison of SNV and SV heterozygosity between groups of species was performed under a phylogenetic generalized least square framework, with taxonomical group and conservation status as explanatory variables. This was followed by Tukey’s post-hoc tests to determine the statistical significance between a pair of groups. Lastly, we conducted demographic reconstruction using psmc-v0.6.5 with our SNV callset. Uncallable regions of the genome as defined by paftools were masked from the analysis using seqkit-v2.4.0. Mutation rate and generation time were obtained from Wang & Obbard 2023^224^ and used for scaling the PSMC results. When species-specific mutation rate and generation time were not available (which was most of the cases), the values from their closest relative who has an estimate were used. Only species with additional high-quality haplotypes that surpassed the VGP metrics and available IUCN conservation categories were included in the summary plot (n = 214 species); the results were similar when using the less stringent 476 species mentioned above, which included more draft quality haplotypes.

### Sex chromosome assignment validation and synteny analysis

When assembling genomes, identifying the sex chromosomes is a non-trivial task that requires multiple lines of evidence. Sex chromosomes were initially assigned during manual curation using a combination of genome-to-genome alignment to a closely related chromosome-level assembly, Hi-C/Pretext evidence including pseudoautosomal region structure where visible, and reduced read depth, typically approximately half autosomal depth, on sex chromosomes in heterogametic individuals (Lariviere et al., to be submitted to bioRxiv). To validate initial sex chromosome identifications and implicate previously unknown sex chromosomes, we implemented multiple approaches including HalfDeep (Gable et al., to be submitted to bioRxiv), SCINKD^225^, GENSPACE^226^, and, where possible, short read alignment. Briefly, we ran SCINKD on 136 species in the data freeze with haplotype-resolved assemblies that were scaffolded to the chromosome-level (Gable et al, to be submitted to bioRxiv). Then, we aligned the two haplotypes using minimap2^218^ and generated visualizations of outlier loci with SVbyEye^227^ to further corroborate sex chromosome assignments. In parallel, we applied HalfDeep, a sequencing depth–based approach, to 578 species from the VGP Phase I dataset. For each species, chromosome-wise HalfDeep proportions were calculated, representing the fraction of regions exhibiting haploid-level depth. These results served as independent lines of evidence to validate and refine sex chromosome assignments.

### rDNA identification

Assembled genomes were scanned for nuclear rDNA with barrnap^228^ (kingdom=eukaryote) to annotate 45S rRNA transcription-unit components, including 18S, 5.8S and 28S rRNA genes. Partial predictions were excluded. For each species and each rRNA type, all full-length features were collected and the corresponding nucleotide sequences were extracted from the genome assembly. To identify internal transcribed spacers (ITS), each 5.8S rRNA hit was used to extract a flanking 30 kbp genomic window, and sequences were retrieved with BEDTools (getfasta)^196^. ITS regions were then identified with ITSx^229^ (taxon = Metazoa; -only_full; -save_regions ITS1,ITS2), and only full predictions delimited by SSU-5.8S (ITS1) and 5.8S-LSU (ITS2) were retained. Quality control computed, per ITS, the fraction of ambiguous bases (N) and the maximum homopolymer length; sequences with N-fraction > 0.05 or any homopolymer > 15 nt were excluded to minimize the influence of low-quality assembly segments and residual long-read sequencing artifacts on downstream analyses.

### Runs of Homozygosity analysis

Genome alignment was done using FastGA^80^, and then paftools, a script written to accompany minimap2^218^, was used to find SNVs in uniquely aligned regions of the genome. To calculate ROH on a given chromosome, we measured the distance between variants in bases, only counting bases that were aligned:

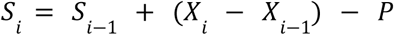

where *S_i_* is the length ‘score’ for ROH at any given variant, *S_i_*_−1_ is the score at the previous variant, *X _i_*− *X_i_*_−1_ represents the distance in aligned bases between the current variant and previous variant, and *P* is the penalty threshold that must be passed to be considered an ROH. We set the penalty *P* at 100,000, so that a homozygous stretch had to be at least 100 kbp in length to be considered a ROH. While we measure the distance in aligned bases, this equation means that ROH can cover non-aligned regions of the genome. To avoid this interfering with results, we masked the ROH using the genome alignment to only measure their length in aligned regions of the genome.

ROH are commonly categorised by their length in bases, with large ROH being 1 Mbp or longer (e.g. <u>Ceballos et al., 2018</u>; <u>Pegollo et al., 2025</u>; <u>Selli et al., 2021</u>).To account for the wide variation in genome size in our dataset, we instead categorised ROH by the proportion of the chromosome that they take up: short (0, 0.5%], medium (0.5%, 1%], or long (1%, 100%]. The cutoff for long ROH was decided as it is approximately the length of a 1 Mbp ROH across human chromosomes. We used the aligned regions of the genomes to calculate both the length of the ROH and the length of the chromosomes and autosomes as a whole. From these we calculated the inbreeding coefficient, F_ROH_, which is simply the proportion of the autosome contained in ROH:

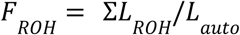

Heterozygosity was also calculated to compare to ROH results. Heterozygosity was calculated in 1 Mbp sliding windows moving along each chromosome in a genome, with the windows overlapping by 500 kbp. If a chromosome was shorter than 1 Mbp in length, the windows were reduced to 100 kbp in length, overlapping by 50 kbp. If any chromosomes or chromosome fragments were shorter than 100 kbp, they were excluded from the analysis. The number of variants between the primary and alternate haplotype in each window was summed, while excluding any part of the window which was contained in a ROH, including any variants within the ROH. To get heterozygosity per kbp in each window, the number of variants was divided by the window length excluding any ROH within the window, and multiplied by 1000:

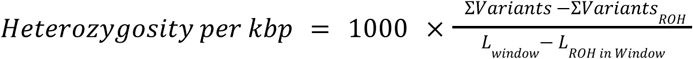

The mean heterozygosity across all windows on a given chromosome was taken to calculate the mean for that chromosome. The average heterozygosity across the whole genome was calculated as the mean of the heterozygosity across all windows on all autosomal chromosomes.

Both F_ROH_ and heterozygosity outside ROH were calculated using a custom snakemake pipeline with python scripts, and the packages *numpy*, *pandas*, and *cyvcf2*^230–233^. They were subsequently visualized using a custom script R with the *tidyverse* and *ggplot2* packages^234,235^. Ordinal logistic regression was done using MASS to test for statistical significance of ROH, heterozygosity, and both together correlating with IUCN status, ranked with Least Concern (LC) as the lowest to Critically Endangered (CR) as the highest^236^. To test for whether the results were influenced by taxonomic relationships, we performed phylogenetic correction and a regression to test whether it influenced the likelihood of a species being threatened (VU to CR) versus non-threatened (LC or Near Threatened, NT). We tested this using Phylolm and MCMCglmm packages in R, with the phylogeny generated for this study using ROADIES^237,238^. Both Phylolm and MCMCglmm showed the same result, with heterozygosity having a statistically significant negative correlation with IUCN status, while the relationship with F_ROH_ was no longer significant.

### Neoplasia prevalence across species

We used publicly available neoplasia tumour prevalence data^148^ and body mass data^239^ for each species. A chordate phylogenetic tree from timetree.org^26^ (downloaded February 2024) was used to quantify molecular rate variation. An existing phylogenetic tree^148^ was used to model tumour prevalence. Among the 169 species of birds and mammals with paired tumour prevalence and body size data, 117 had corresponding genome assemblies as part of the VGP Phase I data freeze (22 species), other long read-based assemblies available in NCBI Datasets (22 species), and short-read chromosomal-level short-read based assemblies submitted 2015 onward in NCBI (73 species)^240^. When multiple assemblies were available for a given species, we selected the most recent genome, prioritizing long-read-based assemblies when available. A further 163 bird and mammal VGP genomes were also included to improve dN and dS rate estimations. Similarly, a subset of amphibian genomes were also included to act as an outgroup for each alignment. The final multiple sequence alignment (MSA) dataset consisted of 280 species of birds and mammals.

By combining 3 known cancer datasets (COSMIC^144^, downloaded June 2025; IntOGen^144,145^, downloaded June 2025; and OncoKB^146^, downloaded June 2025), we curated 1,374 reviewed human genes as candidate orthologs for cross-species genomic analysis. Gene models were lifted over to the selected genomes using Miniprot^74^, and custom scripts were applied to retain the best model per species based on alignment coverage and sequence identity. Models with ⩾75% identity and coverage were kept, with more relaxed thresholds (⩾40%) applied to the outgroup taxa. The resulting one-to-one orthologs were processed through a MSA and a filtering pipeline adapted from OrthoMAM v12^241^ to ensure alignment quality and phylogenetic suitability. Genes represented in fewer than 50 species or lacking an amphibian outgroup were excluded from downstream analyses, leaving a total of 1,194 individual gene MSAs.

To detect variation in the rate of molecular evolution, an adaptive branch-site random effect model (aBSREL^147^) was fitted in Hyphy for each gene. For these Hyphy analyses, each alignment was further filtered to remove species in which over 75% of the sites were missing. If over 75% of the sites were missing within the amphibian outgroup, the gene was removed from all further analyses. Next, for each gene, the aBSREL model was fitted using the chordate time tree of life, with species removed within the alignment if they did not have a match in the time tree of life database, and allowing for double and triple hits within a given site. Once fitted, two trees were produced where the branch lengths were scaled to the branch specific rate of synonymous and non-synonymous evolution respectively. The two trees were rooted on the amphibian outgroup which was then removed. Finally, branches were summed from root to tip for each species to obtain the total amount of synonymous (dS) and non-synonymous (dN) evolution. This process was repeated independently for each gene.

Neoplasia prevalence was modelled throughout using phylogenetic generalized linear mixed models (PGLMMs)^242^ fitted in a Bayesian Markov Chain Monte Carlo (MCMC) framework using the MCMCglmm R package^237^. Neoplasia tumour prevalence was estimated for each gene with separate intercepts for birds and mammals. For genes where only a single class of species was present, e.g. only mammals, a single intercept was fitted. Phylogeny was included as a random effect in every model to account for the shared ancestry. The log_e_ total number of necropsies, log_e_ body mass, log_e_ dN, and log_e_ dS were included as standardized fixed effects, henceforth referred to as the full model (outlined below). That is, each independent variable was transformed to have a mean of 0 and a standard deviation of 1. For genes where both birds and mammals were present, covariates were standardized on a per class basis. At least 5 species needed to be present for a given class to be included in the model. A single slope was estimated for each dependent variable across all species. Marsupials are known to have a divergent immune system from most mammals and thus were excluded from neoplasia modelling. The final paired dataset consisted of 1,169 converged genes spanning up to 109 species (66 mammals and 42 birds).

Full model:

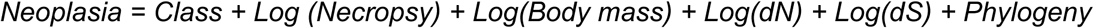

All MCMC chains were run for 1x10^7^ iterations. The first 9x10^6^ iterations were discarded as burn-in and the chain was sampled at every 1,000^th^ iteration. Models were considered to be converged if each of the fixed effects in the model had an effective sample size > 500. MPGLMMS were fitted with a Poisson link to account for the error structure in the benign and malignancy count data. MCMCglmm automatically accounts for overdispersion in count data. We used default priors (µ = 0*_n_* and V = **I** x 10^8^, where 0 is the zero vector and **I** is the identity matrix in which n is equal to the number of fixed effects in the model) for the fixed effects and multivariate parameter-expanded priors (V = 1, n = 1, ⍺µ = 0, and ⍺V = 25^2^) for the phylogenetic random effects (https://devillemereuil.legtux.org).

Regression parameter significance was assessed by the proportion of the posterior distribution that crosses zero (P_x_), where P_x_ < 0.05 is considered to be significantly different from 0.

### Identification of 3D genome folding patterns

Hi-C data were also used to determine different patterns of genome folding ^243^ as previously described ^115,244^. First, Hi-C reads were mapped against their reference genome using GEM v3.6.1 mapper ^245^ from TADbit v1.0.1^246^, with windows from 15 bp to 75 bp in 5 bp steps and reads classified as “self-circle", "dangling-end", "error", "extra dangling-end", "too short", "too large", "duplicated" and "random breaks" were filtered out. HiCExplorer v3.7^113^ was then used to bin the mapped reads into square matrices of 500 kbp resolution, which were subsequently corrected and normalized by scaling the sum of all interactions into the same sequencing coverage (40 million interactions/Gbp of genome size). This normalization method ensures that matrices from all species have the same density of interactions regardless of genome size. Normalized matrices were plotted at a 500 kbp resolution using HiCExplorer v3.7^113^ to identify each chromosome folding pattern. Species with a coverage higher than 3x and a well-defined pattern were included in the analysis, with coverage being calculated considering the number of valid reads and genome size per species.

### Analysis of immunoglobulin loci

#### Detection of immunoglobulin variable genes across vertebrates

The original V gene database of the IgDetective tool was compiled from mammalian species and tested on only mammalian IG loci^22,127,129^. To adapt this strategy to species from other vertebrate classes (ray-finned and cartilaginous fishes, birds, non-avian reptiles, and amphibians) which have distinct sets of V gene lineages, new databases were established by combining V genes available at the International ImMunoGeneTics information system (IMGT^247^) and GenBank databases. Multiple iterations of IgDetective then were further performed to identify V genes. After each iteration, short (≤250 nt), low complexity (one or two nucleotides compose more than *LowComp_MAX_*% of the sequence), and sequences with a premature stop codon in the first reading frame were discarded. The remaining putative V genes (referred to as productive) were incorporated into the database and used in the following searches.

The number of iterations was determined using several indicators. The first indicator was the point at which the number of newly discovered productive V genes plateaued. In birds, this stabilization occurred after the 4th iteration. In some other classes, we observed a sudden increase in gene counts after several iterations; however, most of these additional sequences displayed low complexity. Using a *LowComp_MAX_*=70% threshold removed the majority of false positives across vertebrate classes, although amphibians, ray-finned and cartilaginous fishes required a stricter *LowComp_MAX_*=60% cutoff. A sudden increase in the number of contigs corresponding to the detected V gene candidates was another indicator that the iterative search recruits false positive candidates. In non-avian reptiles, the number of IGHV genes continued to increase across iterations, but the 5th iteration produced an abrupt surge of false positives as well as an increase of contigs containing genes. Therefore, the genes identified by the 4th iteration were retained. IGKV and IGLV gene counts in non-avian reptiles reached saturation before the 4th iteration. In amphibians, gene counts rose steadily, but the accumulation of false positives after the 7th iteration prompted us to stop at the 8th iteration. After the 4th iteration in amphibians and lobe-finned fishes, the same sudden increase in contigs containing V genes in IGK and IGL was observed. In ray-finned fishes, numbers of light chain V genes stabilized after the 3rd iteration, while IGHV gene counts continued to rise. Cartilaginous fishes, analyzed separately because the ray-finned fish database did not perform adequately, showed a similar pattern. For both ray-finned and cartilaginous fishes, the prevalence of false positives in IGH at the 4th iteration led us to finalize the V gene set at that stage.

#### Definition of immunoglobulin loci

An immunoglobulin locus was defined as a run of productive V genes found on the same contig such that distance between two closest V genes do not exceed 1.5 Mbp (productive V genes are defined as in-frame genes without premature stop codons at the end of the first section). If two or more loci were computed for a given chain type, only loci with at least 15 productive V genes were used. In previous work^127^, a smaller threshold (0.3 Mbp) was used to define IG loci which resulted in multiple distal IG loci: situations when two or more IG loci were located on the same contig and separated by more than 0.3 Mbp. While the new 1.5 Mbp threshold masked such cases (partially due to the higher contiguity of VGP genomes), it allowed us to increase confidence in the detected IG loci by increasing the number of V genes in them. The choice of threshold was adjusted using manual curation that included alignment of V genes using the highly sensitive IgBlast tool^248^ and assessing whether the final V genes have full-length alignments to the known V genes. For loci with more than one haplotype per species, the longest locus was used for the analysis. The implemented IG locus refinement procedure has several limitations. First, IG loci were previously identified as IG light chain loci of different subtypes could be merged if they are located close to each other. This decision was made intentionally because minimization of false positive V genes was prioritized in this study. This limitation will be addressed in follow up studies focused on evolutionary development of immunoglobulin chain types. Second, highly diverged V genes were likely missed by the iterative search. Because there is no benchmarked way to detect such IG genes across diverse vertebrate species in a standardized manner, they were outside of the scope of this analysis.

#### Similarity analysis of immunoglobulin V genes

To analyze gene similarities across different types of immunoglobulin chains, 300 V genes were selected from each abundant vertebrate class (birds, squamates, mammals, amphibians, ray-finned fishes, cartilaginous fishes) and each type of immunoglobulin chain (IGH, IGK, IGL, fish light chain loci). If less than 300 V genes were found for a class and chain type, then all of the V genes were used for the analysis. In total, 3,892 V genes were collected. These V genes were aligned using Clustal Omega^248,249^, and the guide tree corresponding to the multiple alignment was computed.

## Supporting information

VGP Phase I Manuscript Extended Author List

## Data availability

All the assemblies in the data sets presented here have been grouped under the VGP umbrella (NCBI BioProject PRJNA489243), with subsets also under other affiliated umbrella projects, including: Darwin Tree of Life (DToL; PRJEB40665), European Reference Genome Atlas (ERGA; PRJEB43510), Bat1K (PRJNA489245), Cetacean Genomes Project (PRJNA1020146), Denmark Yggdrasil (PRJNA955268), Genomics of Brazilian BioDiversity (GBB; PRJNA1180976), Oceanomics (PRJNA1046164), ATLASea (PRJEB64126), AfricaBP (PRJNA811786), and California Conservation Genomes Project (CCGP; PRJNA720569), among others. A subset of the resources generated for this data freeze under the VGP are accessible through a variety of other repositories and genome browsers, including GenomeArk^250^, the GenArk^251^ UCSC genome browser^252^, Ensembl^253^, GoaT^191^, and Genomicus^111,112^. Raw data are also stored in GenomeArk (https://www.genomeark.org) along with intermediate files, annotations, and downstream analyses. GenomeArk2 connects VGP data, intermediate products, and public infrastructure to support end-user access and analysis (**Supplementary Information section 9**).

## Acknowledgements

We acknowledge that this VGP Phase I Flagship study was made possible by the contributions of the coauthors listed here, as well as by the expertise, help, and support of additional individuals and institutions, including field workers, permitting authorities, local collaborators, collection managers, and others who facilitated sample acquisition, access, and data generation.

In particular, we thank the following individuals and institutions for their invaluable contributions: Scott Hamilton (*Lycodopsis pacificus*, *Melanostigma gelatinosum*), Aimee Lang (*Balaenoptera musculus*), Carlos Luis DoNascimiento Montoya (*Trichomycterus rosablanca*), Jeff Jacobsen (*Balaenoptera musculus*), Michelle Smith (long read sequencing at Wellcome Sanger Institute), Chris Conroy (*Ctenodactylus gundi*, *Thomomys bottae* samples from the Museum of Vertebrate Zoology), Adam Ferguson (*Lestoros inca* FMNH 174459, *Amblysomus hottentotus* FMNH 165583, *Anomalurus derbianus* FMNH 222639, *Tarsius syrichta* FMNH 142007 samples from the Field Museum of Natural History), Jane Merkel (*Dermatemys mawii* 17988, *Chitra chitra* 982013, *Platysternon megocephalum* 130681 samples from the Saint Louis Zoo), Jon Fjeldså (*Alca torda* sample from the Natural History Museum of Denmark), Robert Bonde (*Eubalaena glacialis* sample), Jeff Kneebone (*Amblyraja radiata* sample), Rebecca Pugh (*Balaenoptera ricei* sample), Patricia E. Rosel (funding acquisition and sampling for cetacean genomes), Gail Armstrong (*Nyctalus leisleri* sample), Robin Stobbs (*Latimeria chalumnae* sample), Wildlife Conservation Society, Bronx Zoo, Saint Louis Zoo, the government of Malawi (Malawi bat samples) and the Turner Endangered Species Fund (*Gopherus flavomarginatus*). Some data came from cetacean biospecimens that were archived in the National Marine Mammal Tissue Bank as a part of the NIST Biorepository. The Field Museum provided specimens for the VGP and contributed funding for sequencing through its Grainger Bioinformatics Center and Felix Grewe. We also thank the following individuals who assisted the Vertebrate Genome Lab with sample permitting and shipment: David Pasnik, Darla J. Brown-Farnham, Paul R. Stringer, Thomas Wittig, Zachary Ladin, Natchanon Ketram, Chrystie Cucura, Jason Stern, John Boyd, and Matt Price.

This research received general VGP support from the Howard Hughes Medical Institute and Rockefeller University start-up funds to EDJ. Much of the computational work conducted at the Rockefeller University, including genome assembly and annotation and data processing by the Vertebrate Genome Lab and ROADIES analysis performed by C.L. and A.G., utilized the computational resources from the Rockefeller University High Performance Computing Resource Center (RRID: SCR_025889). Over 2.5 million CPU hours that enabled this effort were provided by the Texas Advanced Computing Center (TACC) and the Jetstream2 cloud at Indiana University to support genome assembly and annotation in Galaxy. The VGP consortium is thankful to the directors of these resources, Dan Stanzione and David Hancock. This work was supported in part by the National Center for Biotechnology Information of the National Library of Medicine (NLM), National Institutes of Health (NIH), including the Intramural Research Program of the NIH. The contributions of the NIH authors are considered Works of the United States Government. The findings and conclusions presented in this paper are those of the authors and do not necessarily reflect the views of the NIH or the U.S. Department of Health and Human Services. This work utilized the computational resources of the NIH HPC Biowulf cluster (https://hpc.nih.gov). HalfDeep computations were performed on Delta at the National Center for Supercomputing Applications through allocation BIO250090 from the Advanced Cyberinfrastructure Coordination Ecosystem: Services & Support (ACCESS) program, which is supported by U.S. National Science Foundation grants #2138259, #2138286, #2138307, #2137603, and #2138296. We thank the HPC Service of ZEDAT, Freie Universität Berlin, for computing time. A.N. acknowledges the Penn State Institute for Computational and Data Sciences (RRID:SCR_025154) for providing access to computational research infrastructure within the Roar Core Facility (RRID:SCR_026424). J.F., N.F., A.J.S.B, C.H.T and M.J.O’C would like to thank the Biotechnology and Biological Sciences Research Council UK (award numbers BB/X007332/1 and BB/X003086/1) and Wellcome Trust (grant number 227178/Z/23/Z) for funding and University of Nottingham HPC facilities. M.J.O’C thanks the Leverhulme Trust for her personal fellowship (RF-2024-492). E.G., P.S., S.C. acknowledge funding from NIH R01HG013017 and U01HG013760. P.A.F.G. acknowledge funding from Sao Paulo Research Foundation #18/15579-8 and #25/18246-3.

