## Supplementary Information for "The Vertebrate Genomes Project Phase I: A global reference genome resource"

1. **Species selection criteria**
2. **ROADIES phylogeny**
3. **Retrocopies annotation**
4. **Analyses of k-mer repeat spectra**
5. **Satellite annotation**
6. **Ribosomal RNA (rRNA) genes**
7. **Assembly challenges of bird IGH loci**
8. **Conservation of enhancer-gene associations in vocal learning genes**
9. **GenomeArk2**

### 1. Species selection criteria

We spaced ordinal representatives at approximately even divergence intervals. For mammals and birds, which underwent rapid diversification around the end-Cretaceous, this translated to roughly 50–70 million years between chosen lineages. Notably, molecular clock analyses show that most modern bird orders arose just after the Cretaceous–Paleogene (K–Pg) mass extinction ~66 million years ago[1], and similarly that mammalian orders radiated in the early Paleogene following that event[2]. In contrast, other vertebrate groups have deeper evolutionary splits; for example, the three living amphibian orders (Anura, Caudata, Gymnophiona) diverged in the Paleozoic–early Mesozoic, prior to the breakup of Pangaea[3–5]. Likewise, the common ancestor of birds and crocodilians dates to the Triassic (> 250 MYA). Accordingly, Phase I selections for reptiles, amphibians, and fishes were calibrated to cover these older divergence windows. We incorporated time-calibrated phylogenies from the literature (e.g., for mammals, birds, non-avian reptiles, amphibians, and ray-finned fishes) to guide our choices, ensuring that even ancient and long-branched clades are represented. In addition, a few non-vertebrate chordates (e.g., tunicates or cephalochordates) were included as outgroup references to anchor the vertebrate tree and root the analyses.

The final Phase I roster of species was determined through an expert-driven nomination process. Taxonomic working group chairs for each major lineage (e.g., mammals, birds, etc.) proposed candidate species, which were then consolidated and evaluated by the G10K executive committee. This collaborative vetting ensured that key lineages identified by specialists were not overlooked. In assembling the species list, we gave priority to cases where adding a genome could resolve contested phylogenetic relationships or break up long branches. For instance, inclusion of intermediate lineages helps mitigate long-branch attraction artifacts by providing additional nodes for comparison. Each selected species thus serves a strategic role in the evolving vertebrate tree of life—either by filling a phylogenetic gap or by shortening an otherwise exceptionally long branch in the tree, thereby improving the robustness of downstream evolutionary inferences. We also favored species that could act as genomic "bridges" for annotation, meaning their genomes are similar enough to well-annotated model organisms to allow gene identification by synteny. By sequencing such bridging species, we facilitate comparative genome annotation across distant taxa, since conserved gene order (synteny) can help transfer annotations from one reference genome to another.

Multiple practical considerations were applied alongside these scientific criteria. We preferentially selected species for which some genomic data were already available (short-read drafts or ongoing sequencing efforts), reasoning that existing data could accelerate the assembly of a high-quality reference. Technical feasibility was also critical: we limited selection to species with sufficient body mass (≥0.5 g) to yield high-molecular-weight DNA for long-read sequencing. We saved taxa with extremely large genomes (estimated genome size >6 Gbp) towards the end of the Phase I project, to ensure time for sufficient development of technologies, algorithms, and cost reductions. Among candidates meeting these thresholds, those of high conservation concern received precedence. Critically endangered species that are likely to become extinct in the next 5–10 years were prioritized so that their genetic blueprints can be preserved. This emphasis is both ethical and practical, as the VGP is racing against the ongoing sixth mass extinction to capture genomic data for imperiled taxa. In parallel, we included a number of "iconic" species, taxa of broad public interest or cultural significance, on the rationale that their inclusion would help engage the public and attract funding support for the project. Examples range from charismatic megafauna to emblematic birds and fishes that resonate with conservation efforts. While public appeal was not the sole factor for any selection, it often aligned with scientific merit (many iconic species are themselves evolutionarily unique or endangered).

Finally, the Phase I dataset features species with specialized or convergent traits that are especially informative for comparative genomics. By sequencing such organisms (for example, cave-dwelling fish, hibernating mammals, or vocal-learning mammals and birds and their closest vocal non-learning relatives), we create opportunities to examine the genomic underpinnings of these traits in independent lineages. The insight gained can shed light on the genetic basis of convergent evolution and trait innovation. We also ensured that species important to human medicine and biology are represented. This includes traditional model organisms and others with physiological traits relevant to human health (e.g. disease resistance, longevity, or unique metabolisms).

### 2. ROADIES phylogeny

ROADIES is a fully automated pipeline for species tree inference that starts by randomly sampling loci of fixed length (default: 500 bp) from the input genomes, then infers homologous regions for each locus in the remaining genomes using a whole-genome aligner, followed by inferring multi-copy gene trees, and then summarizes them using a discordance-aware method (ASTRAL-Pro[6]) into the final species tree. To address the wide diversity of vertebrates, we used ROADIES primarily to collect gene trees for four different partitions of the input dataset: (1) *Full dataset (579 species),* where we ran ROADIES in its deep mode using 128,000 sampled loci across all species which produced 29,329 gene trees post-filtration; (2) *Bird-focused subset (139 species: 137 birds + 2 crocodiles as outgroups)*. Since avian phylogeny is particularly complex and we found that it required a larger number of gene trees to be correctly resolved, we increased the sampling depth, generating 54,288 gene trees after filtration steps, from 64,000 sampled loci. (3) *Shark-focused run (partitioned dataset)*. To confidently resolve the relative placement of sharks compared to other fishes and other vertebrates, we partitioned the dataset into three clades (sharks, all fishes, and all remaining species) and enabled ROADIES to sample 8,000 loci evenly across the three groups, resulting in 1,848 gene trees post-filtration. (4) *Fish-focused subset (181 species: 178 fishes + 3 Gymnophiona as outgroups).* Given the challenges in resolving the deep divergence within fishes, we sampled 160,000 loci from this subset, which yielded 37,766 gene trees. Among filtration parameters in ROADIES, we set the MIN_ALIGN (minimum number of species) to 10% of the number of input genomes for all partitions except for the bird-focused subset, where we set this parameter to 4. All other parameters were set to their default settings. The final species tree (**Extended Data Figure 4**) was inferred by combining all resulting gene trees (a total of 123,141) using ASTRAL-Pro3. Since ASTRAL-Pro3 produces unrooted trees by default, we constrained invertebrate species to their known placement in the species tree, re-estimated branch lengths and support values to produce the final tree.

**3. Pseudogene investigation, with focus on retrocopies.**

To extend beyond the capabilities of conventional NCBI EGAP/EGAPx pipelines, we systematically identified and annotated retrocopies (processed pseudogenes) across VGP species using RCPedia algorithm (Galante et al., in preparation; see Methods). Retrocopies originate from the reverse transcription and genomic integration of mature mRNAs and constitute a significant source of de novo genetic novelties, yet are frequently underrepresented in standard annotation resources [7,8].

Our analysis revealed pronounced lineage-specific variation in retrocopy counts per genome (**Extended Data Fig. 5c; Supplementary Table 11**). Mammals exhibited the highest counts (median 5,395; range 289-16,346), though with notable exceptions at both extremes: monotremes (Ornithorhynchus anatinus, 289; Tachyglossus aculeatus, 1,207) harboured markedly fewer retrocopies than other mammals, while species exceeded 10,000 - including Choloepus didactylus (16,346), Dasypus novemcinctus (14,903), and Nycticebus coucang (14,721). Amphibians (median 943; range 210-3,772) and non-avian reptiles (median 702) showed intermediate levels, whereas birds (median 119; range 52–256) and fishes (median 322) generally harboured fewer retrocopies.

Analysis of retrocopy conservation revealed a striking partition across vertebrate lineages (**Extended Data Fig. 5c**). Mammals, birds, and turtles showed the highest proportions of conserved retrocopies (mammals 52.3% [259,803/478,469]; birds 48.7% [3,537/7,263]; turtles 52.6% [2,075/3,945]0, consistent with long-term retrogene retention. In contrast, crocodilians (8.1% [20/247]), lepidosaurs (9.4% [774/8,200]), cartilaginous fishes (4.3% [603/14,081]), lobe-finned fishes (0.3% [2/660]), and amphibians (1.4% [298/21,861]) were overwhelmingly species-specific, and ray-finned fishes occupied an intermediate position (28.3% [6,266/22,128]). These patterns reflect lineage-specific differences in retrotransposition activity: lineages with high activity (e.g., mammals) accumulate abundant young, species-specific retrocopies, whereas others with reduced activity retain proportionally more conserved, older copies, a distinction corroborated by further analysis and presented in the companion manuscript (Galante et al., in preparation).

A lineage-dependent fraction of retrocopies (25-55%) lacked intact ORFs, consistent with the rapid mutational decay expected following retroduplication [9]; Extended Fig. 5d). Among those retaining coding potential, a fraction displayed significant signatures of purifying ($dN/dS < 0.5$) or positive ($dN/dS > 1.2$) selection ($p$-value $< 0.05$; **Extended Data Fig. 5d**).

Ray-finned fishes, birds, and amphibians showed the highest proportions of retrocopies under detectable selection, consistent with active retrogene recruitment in these lineages. Strikingly, species-specific retrocopies in non-mammalian lineages, particularly amphibians, birds, lepidosaurs, and fishes, were strongly enriched for low dN/dS values (**Extended Data Fig. 5e**), indicating that purifying selection is already acting on recently originated retrogenes that have acquired functional roles. In contrast, mammals showed comparable dN/dS distributions between conserved and species-specific retrocopies (**Extended Data Fig. 5e**), consistent with evidence that mammalian retrocopies recruitment is a continuous process distributed across evolutionary time [8,10,11].

**4. Analyses of k-mer repeat spectra**

The percentage of the genome consisting of 31-mers that appear at least twice in the genome, referred to as the repeat ratio (RR) hereafter, increased substantially in VGP assemblies compared to 248 existing assemblies that we identified (**Fig. 2b**). Thus, while the corresponding 248 assemblies in VGP are on average (median) only 15% (9%) larger than existing ones, their RR is 180% (69%) higher, confirming that the repetitive regions dominate missing parts of existing genomes. Across the tree, the biggest relative increase was observed among birds. While birds have among the least repetitive genomes, the draft-quality bird genomes (mostly from B10K) are far less repetitive than high-quality VGP genomes, making this clade stand out in terms of its increase in RR. The number of existing genomes that are more repetitive than VGP genomes was relatively small (e.g., only 39 existing genomes had at least 5% higher RR than VGP compared to 190 genomes in the reverse direction).

Comparing the phylogenetic independent contrast (PIC) of RR and assembly length showed that generally, phylogenetic nodes with higher contrast in RR between their children have a higher contrast in length as well (**Extended Data Fig. 6a**). However, Spearman's correlation between PIC of RR and assembly length was stronger for VGP genomes compared to existing genomes ($\rho$=0.65 vs 0.77; **Extended Data Fig. 6a**). The increase in correlation is even stronger for “cherry” nodes with only two sister species under them ($\rho$=0.72 for existing vs 0.92 for VGP). This is because changes in the genome length and 31-mer RR tend to track each other closely among closely related species, but assembly incompleteness can mask this strong correlation. For example, in VGP samples, changes in RR and assembly size of chimpanzee and bonobo are consistent with other species (roughly, 2% decrease in genome size and 8% decrease in RR going from chimp to bonobo); however, the abnormally low repeat content of the previous draft bonobo genome creates an outlier in PIC correlations among existing genomes.

To further investigate the correlation between 31-mer RR and genome length, we performed phylogenetic regression for each extended lineage separately, including all 568 ‘vertebrate’ genomes of Phase I (**Extended Data Fig. 6b**). Results confirm that RR and assembly length generally correlate, with notable strong deviations from the general trend. Among amphibians, five genomes were at least 20 Gbp and their RR ratio, while indicative of highly repetitive genomes, would not predict such extremely large genomes. A similar pattern was observed for unusually large genomes in the elephant shrew among mammals and goby and Reedfish among fish.

### 5. Satellite annotation

VGP assemblies remain contiguous across many repetitive regions, allowing us to catalogue satellite DNA, Satellites are large tandem-repeat fields characteristic of centromeric, pericentromeric, and subtelomeric heterochromatin. From each genome of the VGP Phase I data freeze, we extracted the satellite-DNA-proper fraction, defined as the longest arrays, arrays at contig ends, and arrays flanking assembly gaps. We then validated each monomer consensus against the canonical 23-mer spectrum of its respective genome, retaining only sequences with counts >100. This removed 17% of arrays as possible low-copy assembly artefacts and corrected a further 15% with miscalled consensuses, leaving 36,216 validated arrays grouped into 3,429 recurrent satellite families across 577 species. Per-clade median satellite-field lengths ranged from 69 kbp in ray-finned fishes to 1.04 Mbp in primates, while the longest individual arrays reached 11.5 Mbp in a cartilaginous fish (*Scyliorhinus*) and 26.1 Mbp in primate α-satellite.

Of the 3,429 satellite families, 94% were restricted to a single species (**Extended Data Fig 6c**), and no complex satellite was shared between vertebrate classes beyond the telomeric repeat $(CCCTAA)_n$. Neither 23-mer cleaning nor relaxing the cross-species identity threshold to the alignment floor (~62%) altered this result. At the NCBI order rank, however, each well-sampled order was dominated by a characteristic satellite. Re-clustering at class, superorder, and order ranks showed that order is the deepest level at which a satellite is shared: expanding an order label to its superorder added no further species to the dominant family (**Extended Data Fig. 6c**). These findings suggest that satellites were strongly lineage-specific.

Satellite-field location, length, and characteristic sequence varied by clade (**Extended Data Fig. 6c**). In mammals, which contained 943 satellite families, 45% of fields occurred at contig ends and 11% flanked internal gaps. Dominant satellites included the artiodactyl macrosatellite (1,760 bp, 23 species), primate α-satellite (171 bp, 7 species; the largest fields, up to 26 Mbp, 87% interstitial), the bat satellite (Chiroptera, 418 bp), and the carnivore satellite (738 bp). Birds, with 607 families, had the highest proportion of contig-end fields (51%) and the highest median complex-array period (485 bp). Songbirds (Passeriformes) carried a 696-884 bp satellite in 12 species, together with a 191-bp pericentromeric satellite in 11 species, while Charadriiformes (180 bp, 8 species), Accipitriformes (173 bp, 7 species), and Falconiformes (174 bp, 6 species) each carried distinct order-specific satellites. Ray-finned fishes, with 1,023 families, had the highest proportion of interstitial satellites (53%), which resolved into many order-specific satellites, led by a carp satellite (Cypriniformes, 1,022 bp, 9 species). No other ray-finned order recurred beyond three species; even the well-sampled Perciformes (16 species) capped at three, the fastest satellite turnover in the panel. Reptiles, with 312 families, separated cleanly into a squamate satellite (Squamata, 185 bp, 15 species) and a distinct, GC-richer turtle satellite (Testudines, 222 bp, 5 species). Amphibians, with 292 satellite families, had the highest fraction of gap-flanking fields (24%) and included a salamander specific satellite (Caudata, 311 bp). Cartilaginous fishes, with 128 satellite families, carried the longest fields relative to their number but did not have an order-specific satellite across ≥ 2 species; they also had the shortest periodic quantum, with a median of 207 bp. Lack of an order specific satellite could be due too deeply divided into multiple orders.

Across clades, monomer lengths clustered at integer multiples of the nucleosome repeat (**Extended Data Fig. 6c**, dashed lines). The primate α-satellite (171 bp), Squamata satellite (185 bp), and three different bird-order satellites (Accipitriformes 173 bp, Charadriiformes 180 bp, and Falconoformes 174 bp) each corresponded to approximately one nucleosome, whereas the Artiodactyla satellite and other longer satellites fell near higher-order multiples (~10×). These patterns suggest that nucleosome organization constrains the unit length at which satellites expand.

Although most VGP assemblies are chromosome-scale rather than telomere-to-telomere, the satellite arrays flanking many centromeric gaps were nonetheless assembled. As a result, a metacentric centromere appeared as a satellite–gap–satellite configuration, as seen for example in the zebra finch 191-bp pericentromeric satellite flanking a 696-bp core repeat. We identified 9,254 such configurations in 540 assemblies, including 1,550 in which both flanks abutted an internal gap. A karyotype control confirmed that this signal reflects centromere architecture rather than satellite abundance. For instance, in mouse chromosomes, which are acrocentric[12], the major (234 bp) and minor (120 bp) satellites were recovered, but 97% of major-satellite arrays occurred at contig ends, and no flanked-gap configuration was found. By contrast, other rodent suborders, including those represented by the ground squirrel, beaver, and marmot, each yielded 8–12 such configurations. VGP assemblies that are HiFi-based therefore locate centromeres and recover their satellite sequence, while VGP telomere-to-telomere assemblies that incorporate both HiFi and ONT are needed to close the centromeric cores (Komissarov et al., in preparation).

#### 6. Ribosomal RNA (rRNA) genes

Ribosomal RNA (rRNA) genes occupy a central position in both cellular metabolism as a biological process and evolutionary biology as a field of study. As the structural and catalytic core of the ribosome, rRNAs are indispensable to protein synthesis and among the most abundant transcripts in all living cells, accounting for up to 80–90% of total RNA in the vertebrate transcriptome. Their evolutionary roots trace back to the last universal common ancestor (LUCA), making them some of the most ancient and highly conserved genetic sequences on Earth[13]. This extreme conservation, particularly in the 18S, 5.8S, and 28S rRNAs, has made them foundational markers for reconstructing deep phylogenetic relationships across all domains of life, including the major lineages of vertebrates[14]. In striking contrast, the internal transcribed spacers (ITS1 and ITS2) that separate these rRNA genes evolve rapidly under minimal functional constraint. These highly divergent non-coding regions, shaped by concerted evolution and frequent sequence turnover, provide powerful resolution among closely related species and are widely used for molecular “barcoding”[15].

Analyses of VGP Phase I genomes across vertebrate orders reveal striking, intra- and intertaxonomic variation in ribosomal spacers (**Fig. 3i**). ITS1 length ranges from ~185 to 3,197 bp (≈17-fold difference), and ITS2 length ranges from ~217 to 3,045 bp (≈14-fold difference). GC content also spans remarkably broad ranges: ITS1 from ~31% to 85%, and ITS2 from ~48% to 85%, reflecting substantial shifts in nucleotide composition across vertebrates. Despite their immediate genomic adjacency, ITS1 and ITS2 have diverged largely independently—ITS1 reaching its greatest lengths in birds and its shortest in ray-finned fishes, whereas ITS2 is longest in mammals and shortest in amphibians. These extremes—rRNA gene conservation on one end and ITS hyperdivergence on the other—capture a dual tempo of genome evolution, spanning the nearly immutable molecular core of translation and the rapidly evolving spacers that encode the most recent branches of the vertebrate tree.

#### 7. Assembly challenges of bird immunoglobulin heavy chain loci

Almost all bird immunoglobulin heavy chain (IGH) loci were found on short contigs: 78% of contigs containing IGH loci were shorter than 750 kbp, indicating a systematic challenge in assembling IGH loci in bird genomes (**Extended Data Fig. 9b**). To better understand this challenge, we analyzed the T2T zebra finch reference genome constructed using both PacBio High Fidelity (HiFi) and Oxford Nanopore (ONT) reads. The zebra finch IGH locus was found in a highly repetitive region that contains multiple tandem repeats on dot chromosome 36 (accession CM109827.1), a total chromosome length of 4.8 Mbp (**Extended Data Fig. 9c**). While ONT reads retained sufficient coverage in this region, in line with known sequencing bias issues with PacBio HiFi sequencing of dot chromosomes[16], read coverage demonstrated a significant drop. This drop matches a 1.1 Mbp region of elevated homopolymer rate (~75% of consecutive positions contain the same nucleotide instead of expected ~25%). Considering that other bird genomes were assembled using HiFi reads only, it is possible that similar effects led to assembly fragmentation for chromosomes carrying IGH loci but did not substantially affect our ability to detect them in the fragmented assemblies.

#### 8. Conservation of enhancer-gene associations in vocal learning genes

Using the VGP 577-way Progressive Cactus whole-genome alignment, we assessed the synteny conservation of published predicted PVALB interneuron open chromatin regions that are convergent in vocal learning mammals[17] across 280 extant vertebrate genomes spanning mammals, birds, reptiles and fishes. To transfer peak coordinates across species, we used halLiftover, a tool that leverages the HAL (Hierarchical Alignment Format) graph structure of the Progressive Cactus alignment to map genomic intervals from the human reference to each target genome, preserving alignment context across the vertebrate phylogeny[18]. Genomes for which no peaks could be successfully lifted over, including reconstructed ancestral nodes, were excluded from downstream analyses, yielding 280 extant species with synteny data available. Synteny retention was then defined as the proportion of peaks that maintained their nearest-gene association in each target genome relative to the human reference, with liftover rate correlating positively with synteny retention across species, consistent with increasing sequence divergence at greater evolutionary distances. Per-peak analysis revealed substantial heterogeneity in conservation, with a subset of elements showing broad synteny retention across all clades and the majority conserved primarily within mammals (**Extended Data Fig. 10e**). PVALB interneuron peaks showed significantly higher synteny retention than background PVALB interneuron regulatory elements across all vertebrate clades (Wilcoxon rank-sum test, Benjamini-Hochberg corrected $p<0.001$ in all clades), with retention highest in mammals and decreasing progressively with evolutionary distance (**Extended Data Fig. 10f**). The consistent enrichment of synteny retention in vocal learning-associated regulatory elements relative to background regions across all clades suggests that these enhancer-gene associations have been maintained under purifying selection, and demonstrates that the

VGP whole-genome alignment enables genome-wide detection of functionally relevant regulatory conservation beyond nucleotide-level variant analysis.

#### 9. GenomeArk2

To make VGP assemblies and their underlying data discoverable and immediately analyzable, we developed GenomeArk2 (https://genomeark2.org), a web portal that adds an analytics layer over the same GenomeArk data repository (https://www.genomeark.org/) referenced throughout this work. The portal is built on the open-source infrastructure (https://github.com/galaxyproject/brc-analytics) and presents a curated, continuously updated catalog of genomes indexed against the NCBI taxonomy. Through a browser-based interface, users can search and filter the catalog by organism, taxonomic group, and assembly accession; inspect per-assembly metadata; and follow direct links to the underlying sequence, read, and annotation files. From the same interface, users can launch Galaxy workflows on a selected assembly. Reference genome identifiers, FASTA, and annotation files are substituted into workflow parameters automatically at runtime, connecting genome discovery directly to the no-cost ACCESS-CI compute backend described above. These functions allow a researcher to move from browsing to analysis without leaving the portal. GenomeArk2 is presently focused on single-assembly access and analysis. Future functions will support comparative and population-scale genomics including interrogation of multiple genome alignments, pangenome graphs, orthology relationships, and signatures of evolutionary selection.


1. Jarvis, E. D. *et al.* Whole-genome analyses resolve early branches in the tree of life of modern birds. *Science* **346**, 1320–1331 (2014).
2. Meredith, R. W. *et al.* Impacts of the Cretaceous Terrestrial Revolution and KPg extinction on mammal diversification. *Science* **334**, 521–524 (2011).
3. Hime, P. M. *et al.* Phylogenomics reveals ancient gene tree discordance in the amphibian tree of Life. *Syst. Biol.* **70**, 49–66 (2021).
4. Pyron, R. A. Divergence time estimation using fossils as terminal taxa and the origins of Lissamphibia. *Syst. Biol.* **60**, 466–481 (2011).
5. Feng, Y.-J. *et al.* Phylogenomics reveals rapid, simultaneous diversification of three major clades of Gondwanan frogs at the Cretaceous-Paleogene boundary. *Proc. Natl. Acad. Sci. U. S. A.* **114**, E5864–E5870 (2017).
6. Zhang, C. & Mirarab, S. ASTRAL-Pro 2: ultrafast species tree reconstruction from multi-copy gene family trees. *Bioinformatics* **38**, 4949–4950 (2022).
7. Conceição, H. B. *et al.* RCPedia: a global resource for studying and exploring

retrocopies in diverse species. *Bioinformatics* **40**, (2024).
