## Extended Data Figures for "The Vertebrate Genomes Project Phase I: A global reference genome resource"

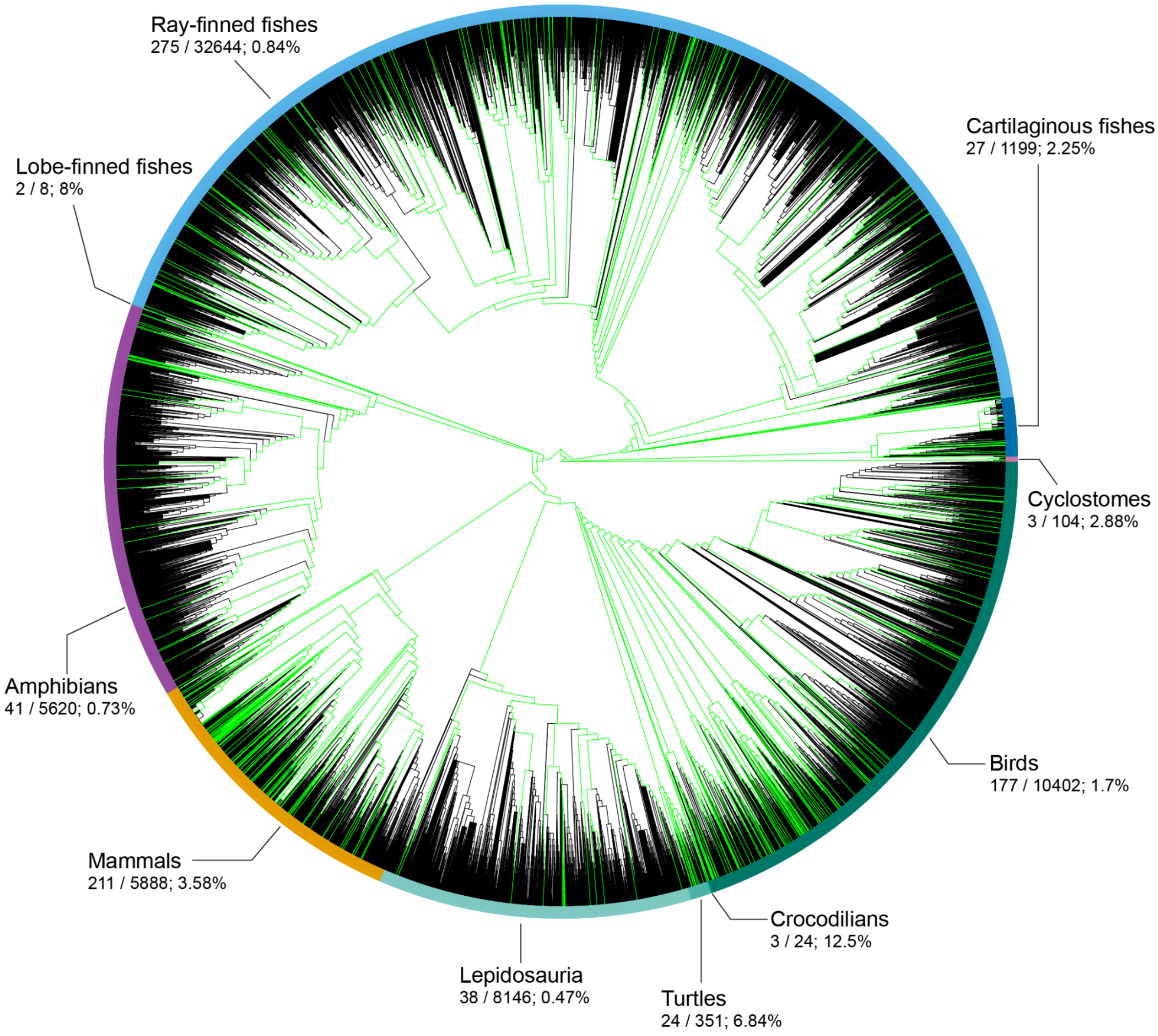


**Extended Data Fig. 1.** Phylogenetic distribution and taxonomic coverage of VGP Phase I genomes across vertebrate diversity. Circular phylogenetic tree of vertebrate species represented in the Open Tree of Life database[1]. All species with a main haplotype genome assembly in the expanded VGP Phase I dataset that can be mapped to the database are highlighted in green (801 species). For VGP species not represented as terminal tips in the plotted species-level OpenTree phylogeny, the nearest available plotted taxon identified from the OpenTree topology was highlighted as a visualization proxy. The colored outer ring indicates the major vertebrate Extended Lineages. For each Extended Lineage, the accompanying values report the number of VGP Phase I species with a genome assembly, the estimated total number of extant species represented by the corresponding ordinal lineages, and the resulting percentage of vertebrate diversity sampled in Phase I. Non-vertebrate outgroups were excluded from the tree and coverage calculations.

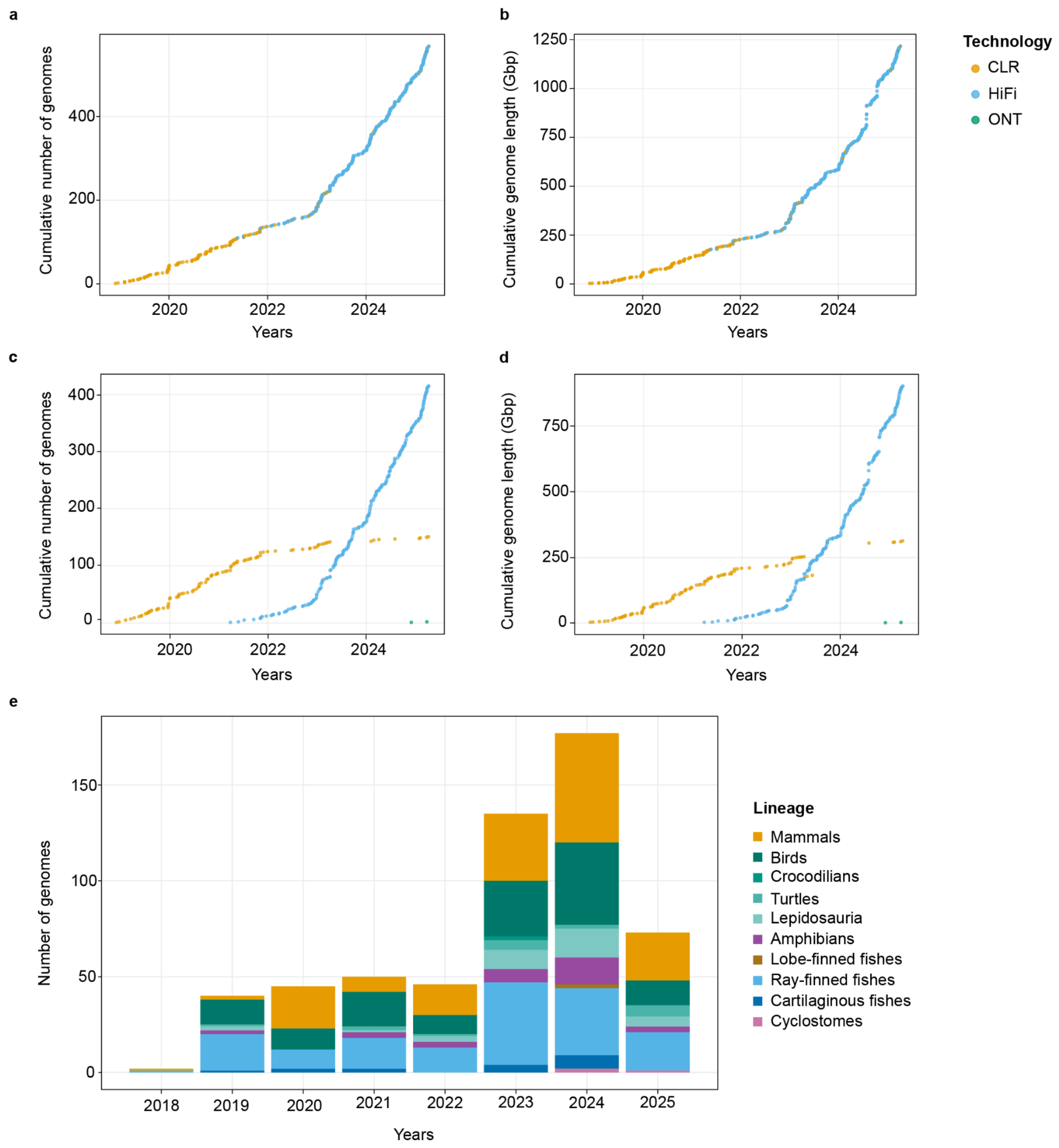


**Extended Data Figure 2. Number of VGP-associated genomes released in NCBI/ENA Genbank over an 8-year period.** Data is shown by genome release date in GenBank (x-axis) versus cumulative number of species sequenced (**a**) or Gbp sequenced (**b**) (y axis). The same data is also separated by sequencing technology (**c**, **d**). **e**, Number of species genomes completed in each major vertebrate group across the 8-year period.

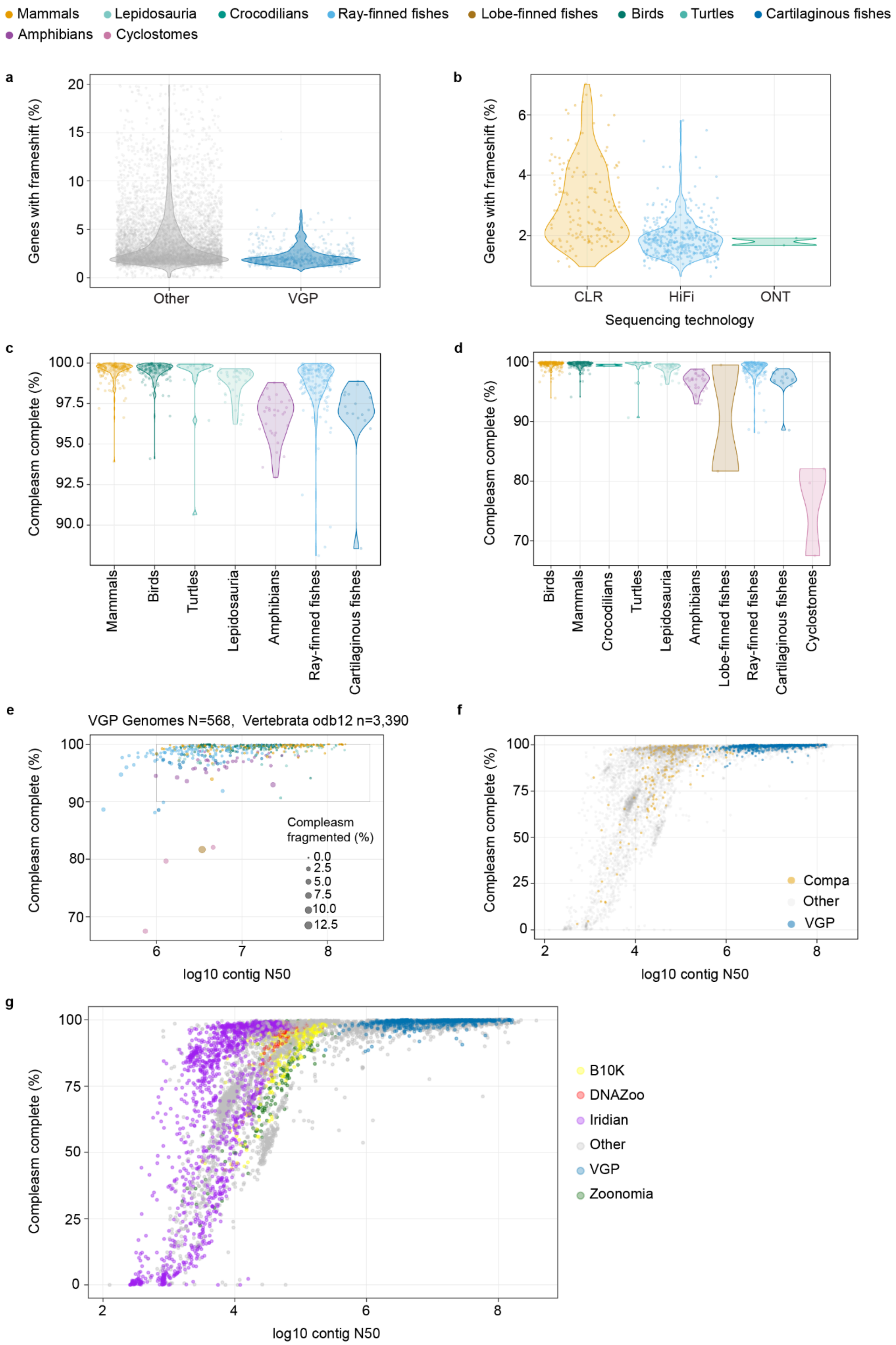

Mammals
Lepidosauria
Crocodilians
Ray-finned fishes
Lobe-finned fishes
Birds
Turtles
Cartilaginous fishes
Amphibians
Cyclostomes
a
Genes with frameshift (%)
Other
VGP
b
Genes with frameshift (%)
CLR
HiFi
ONT
Sequencing technology
c
Compleasm complete (%)
Mammals
Birds
Turtles
Lepidosauria
Amphibians
Ray-finned fishes
Cartilaginous fishes
d
Compleasm complete (%)
Birds
Mammals
Crocodilians
Turtles
Lepidosauria
Amphibians
Lobe-finned fishes
Ray-finned fishes
Cartilaginous fishes
Cyclostomes
e
VGP Genomes N=568, Vertebrata odb12 n=3,390
Compleasm complete (%)
Compleasm fragmented (%)
0.0
2.5
5.0
7.5
10.0
12.5
log10 contig N50
f
Compleasm complete (%)
Compa
Other
VGP
log10 contig N50
g
Compleasm complete (%)
B10K
DNAZoo
Iridian
Other
VGP
Zoonomia
log10 contig N50

**Extended Data Figure 3. Additional assembly QC.** **a**, Percentage of BUSCO genes from the vertebrata_odb12 set identified and containing frameshift mutations determined by compleasm for the VGP genomes (blue) and other genomes in GenBank (grey). **b**, Percentage of BUSCO genes containing frameshifts in the VGP genomes split by long-read sequencing technology. **c**, Percentage of BUSCO genomes from the vertebrata_odb12 dataset identified as complete by compleasm for the VGP genomes. **d**, Same as (**c**), but including lobe-finned fishes and Cyclostomes. Note that compleasm in genome mode often provides a lower bound for assembly completeness, as highly complete gene annotations for the same assemblies typically yield slightly higher completeness scores when compleasm is run in protein mode (**Supplementary Table 7**). **e**, Gene completeness (y-axis) vs Contig N50 (log10 x-axis) of the VGP genomes colored by lineage group (same colors as in **d**). The size of each bubble shows the percentage of fragmented BUSCO genes identified. **f**, Gene completeness vs contig N50 (log10) of all vertebrate genomes in GenBank with the VGP set coloured in blue and comparator set of genomes from the same species colored in orange. **g**, Same as in (f), but colored by each BioProject representative genome in GenBank. Those in grey dots are from other submitters.

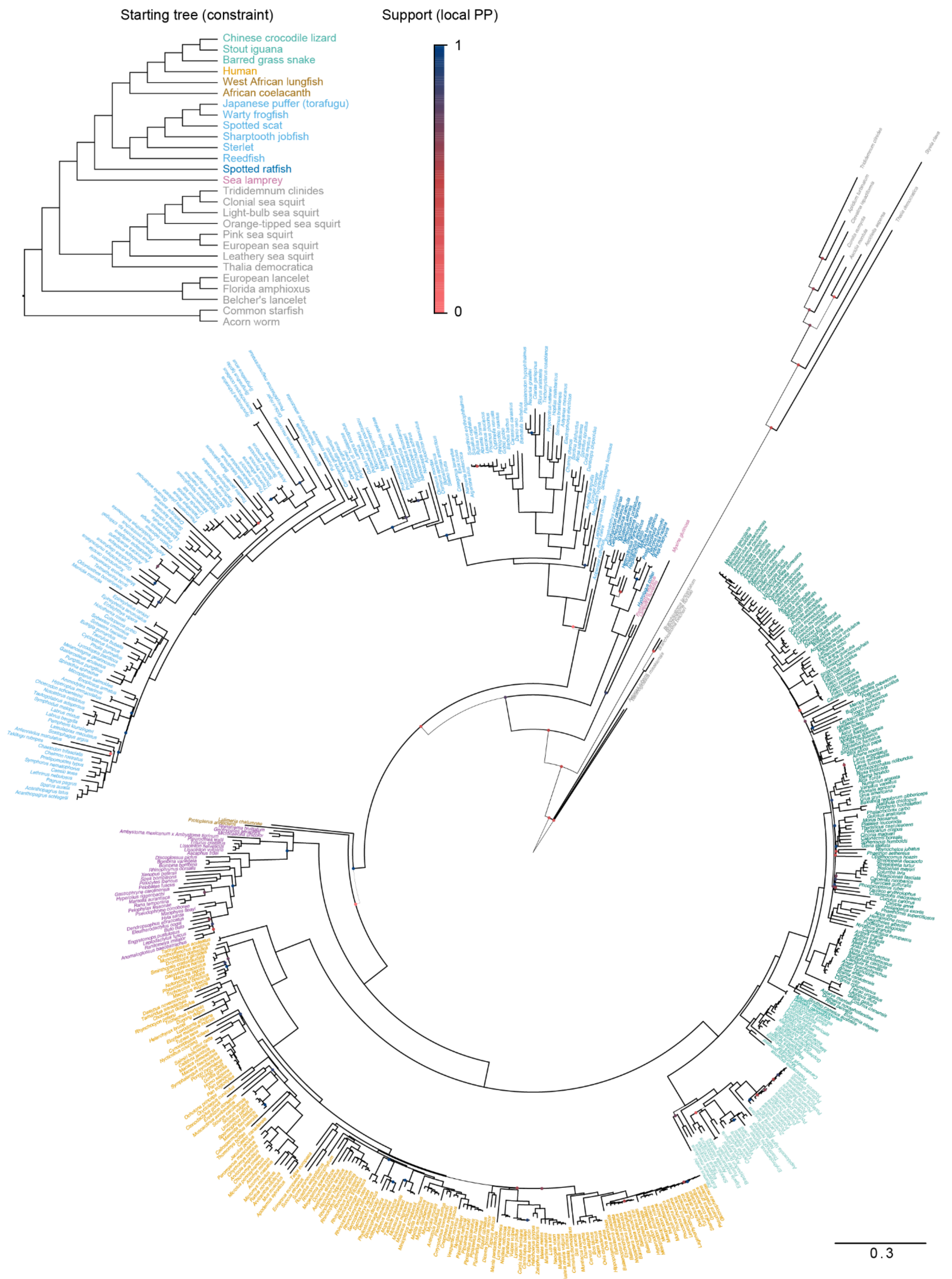


**Extended Data Fig. 4. The ROADIES[2] tree inferred using 123,141 loci sampled across 579 vertebrate species**. Node circle colors show local posterior probability support, as computed by ASTRAL-Pro[3], with nodes with no circle signifying 100% support. Note that relationships among many neoavian orders have relatively low support. Invertebrates had very sparse sampling among ROADIES locus trees, making their branch length less reliable. Similarly, several other branches were hard to resolve without a full genome alignment. Thus, we started the ASTRAL-Pro run within ROADIES from a starting tree (upper left), which ensured the correct placement of invertebrates, as well as the accepted placement of coelacanth, jawless fishes (with respect to sharks), snappers, and snakes.

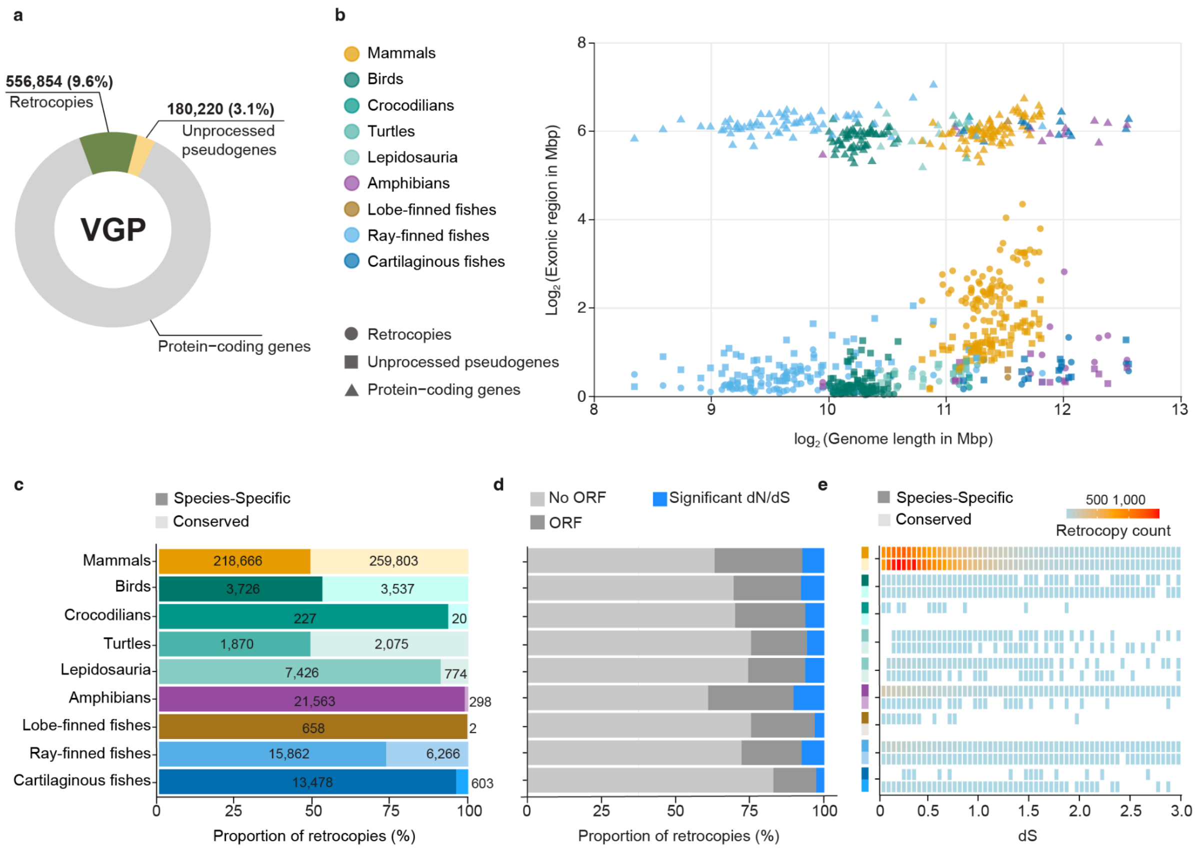


**Extended Data Figure 5. Diversity, conservation, and selective pressure on retrocopies and unprocessed pseudogenes across VGP vertebrate genomes. a**, Total counts of retrocopies and unprocessed (DNA) pseudogenes, relative to protein coding genes. **b**, Exonic content ($\log_2$ Mbp) plotted against genome length ($\log_2$ Mbp) for protein-coding genes (triangles), retrocopies (circles), and unprocessed pseudogenes (squares), for each species, color-coded by clade. **c**, Proportion of species-specific (dark) and conserved (light) retrocopies per clade. Absolute counts are indicated within each bar. **d**, Proportion of retrocopies per clade classified as: lacking intact ORFs, retaining an ORF without significant signal of selection, or retaining an ORF under significant selection. **e**, Distribution of dN/dS values for species-specific and conserved retrocopies across species. Cell color intensity reflects the number of retrocopies per dN/dS bin per species (0-4,000).

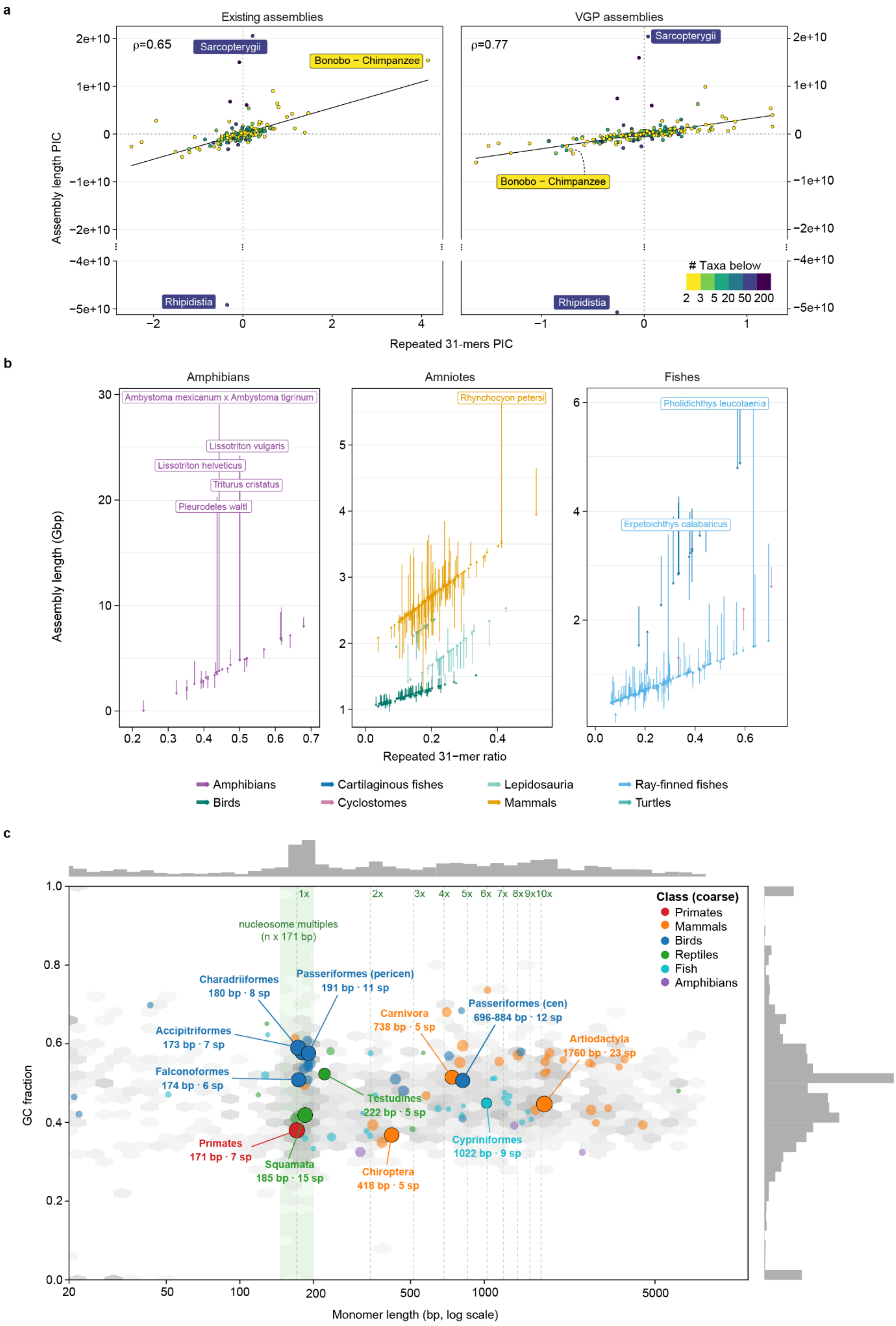

a
Existing assemblies
VGP assemblies
ρ=0.65
ρ=0.77
Sarcopterygii
Bonobo − Chimpanzee
Rhipidistia
### Taxa below
2 3 5 20 50 200
Assembly length PIC
Repeated 31-mers PIC
b
Amphibians
Amniotes
Fishes
Ambystoma mexicanum x Ambystoma tigrinum
Lissotriton vulgaris
Lissotriton helveticus
Triturus cristatus
Pleurodeles waltl
Rhynchocyon petersi
Pholidichthys leucotaenia
Erpetoichthys calabaricus
Assembly length (Gbp)
Repeated 31−mer ratio
Amphibians
Birds
Cartilaginous fishes
Cyclostomes
Lepidosauria
Mammals
Ray-finned fishes
Turtles
c
nucleosome multiples
(n x 171 bp)
1x 2x 3x 4x 5x 6x 7x 8x 9x 10x
Class (coarse)
Primates
Mammals
Birds
Reptiles
Fish
Amphibians
Charadriiformes
180 bp · 8 sp
Passeriformes (pericen)
191 bp · 11 sp
Accipitriformes
173 bp · 7 sp
Falconoformes
174 bp · 6 sp
Carnivora
738 bp · 5 sp
Passeriformes (cen)
696-884 bp · 12 sp
Artiodactyla
1760 bp · 23 sp
Testudines
222 bp · 5 sp
Cypriniformes
1022 bp · 9 sp
Primates
171 bp · 7 sp
Squamata
185 bp · 15 sp
Chiroptera
418 bp · 5 sp
GC fraction
Monomer length (bp, log scale)

**Extended Data Figure 6. Phylogenetic signals of 31-mer frequencies versus assembly length for the 568 vertebrate genomes.** We define the percentage of the genome composed of k-mers that appear at least twice as the 31-mer repeat ratio (RR). We performed maximum likelihood ancestral state reconstructions of RR and assembly length using a Brownian motion model with the phylotools based on the ROADIES tree. **a**, We show phylogenetic independent contrasts (PIC) to compare changes in RR and assembly length at internal nodes of the tree, focusing on 248 VGP genomes that had a comparable non-VGP (existing) assembly. The PIC value indicates the difference between the two descendant lineages of each node. We show that PICs of RR and assembly length correlate according to Spearman correlation (ρ shown), and the correlations are stronger with VGP genomes. The increase in correlation is even stronger for “cherry” nodes (yellow) with only two sister species under them ($\rho=0.72$ for existing vs 0.92 for VGP). Outliers are indicated on the figure. Some outliers when using non-VGP genomes (e.g., human/chimp) are due to differing genome qualities among existing assemblies, a problem solved in VGP. **b**, Using the entire tree (568 genomes), we fit a linear model correlating RR to assembly length using a phylogenetic covariance matrix computed under a Brownian model. We fit a model per lineage and removed outliers marked on the figure from model fitting. The learned regression assigns an “expected” assembly length for each genome based solely on its RR (arrow head), which we compare to the actual assembly length (arrow tail). The outlier genomes with large deviations from the predicted value (named in the figure) have repeat structures that are substantially different from other assemblies from the same lineage. **c**,GC fraction versus monomer length for vertebrate satellite families. We plot each validated satellite family by its consensus monomer length (bp, log scale) against its GC fraction, coloured by coarse taxonomic class. Small background points show all families in the dataset. Marginal histograms show the distributions of monomer length (top) and GC fraction (right). The green shaded band and vertical dashed lines indicate integer multiples of the nucleosome repeat unit (1×–10×).

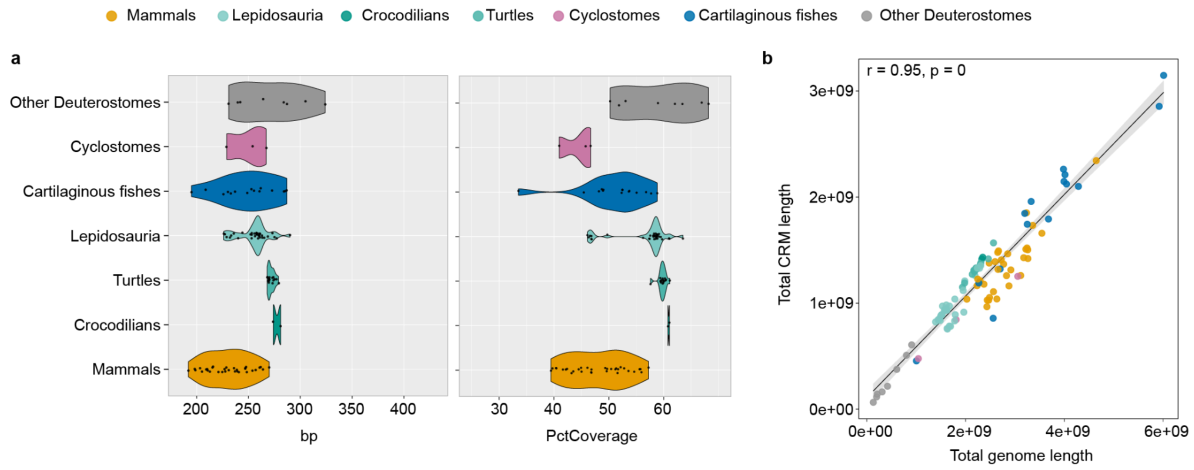


**Extended Data Figure 7. Annotation of regulatory elements. a**, Distribution of average cis-regulatory module (CRM) lengths across vertebrate clades (left). Violin plots show the variation in mean CRM size among species within each clade, with individual points representing species. Mammals, birds, reptiles, amphibians, cartilaginous fishes, ray-finned fishes, cyclostomes, and deuterostome invertebrates are shown separately. Percent of genome coverage by predicted CRMs across vertebrate orders (right). Violin plots represent the percentage of each genome annotated as regulatory sequence, with points representing individual species. **b**, Relationship between total genome length and total CRM length using linear regression analysis (Pearson's correlation). Each point represents a species, colored according to vertebrate order (left panel).

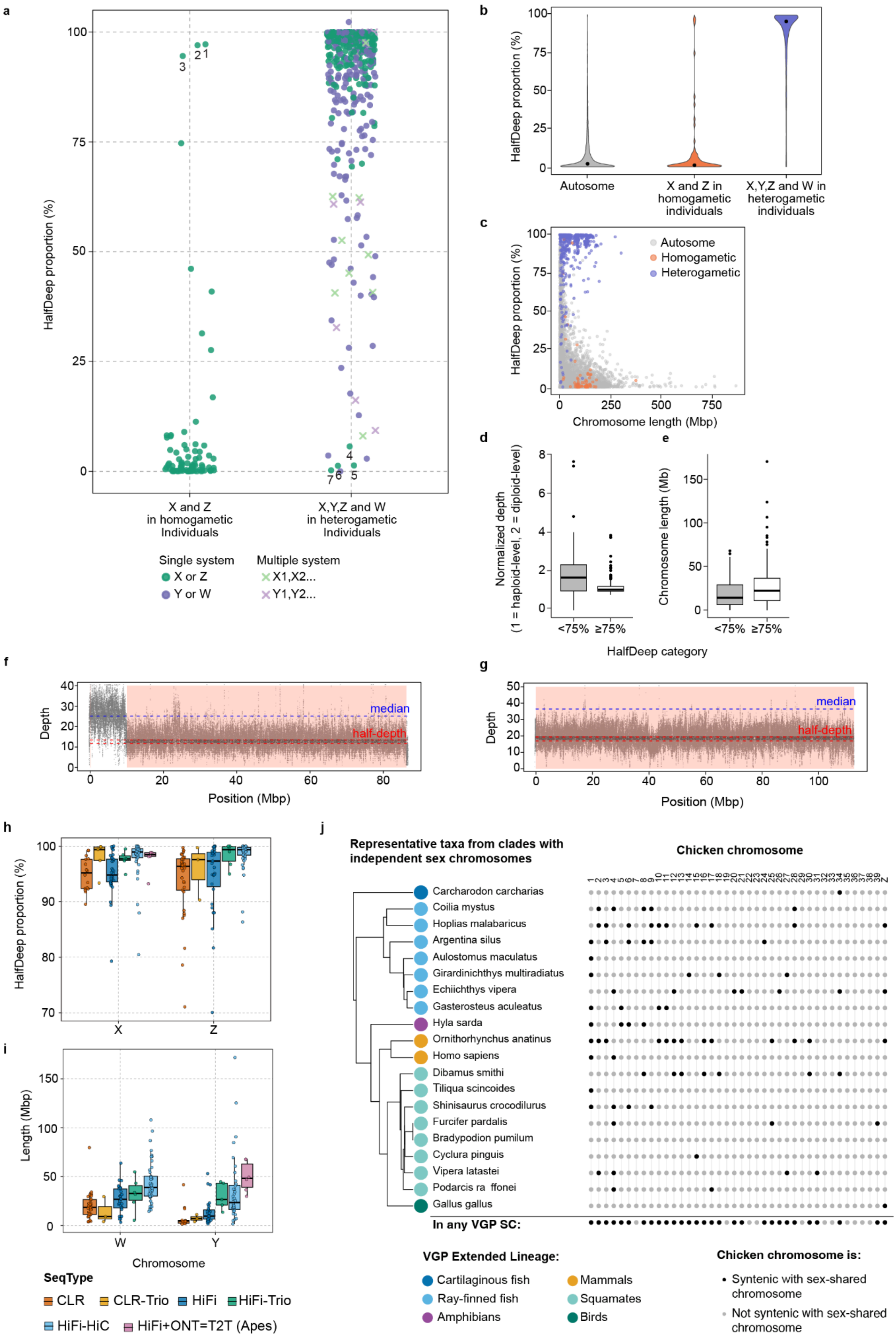
a
HalfDeep proportion (%)
X and Z in homogametic Individuals
X,Y,Z and W in heterogametic Individuals
Single system
X or Z
Y or W
Multiple system
X1,X2...
Y1,Y2...
b
HalfDeep proportion (%)
Autosome
X and Z in homogametic individuals
X,Y,Z and W in heterogametic individuals
c
HalfDeep proportion (%)
Autosome
Homogametic
Heterogametic
Chromosome length (Mbp)
d
Normalized depth (1 = haploid-level, 2 = diploid-level)
e
Chromosome length (Mb)
<75%
≥75%
HalfDeep category
f
Depth
median
half-depth
Position (Mbp)
g
Depth
median
half-depth
Position (Mbp)
h
HalfDeep proportion (%)
X
Z
i
Length (Mbp)
W
Y
Chromosome
SeqType
CLR
CLR-Trio
HiFi
HiFi-Trio
HiFi-HiC
HiFi+ONT=T2T (Apes)
j
Representative taxa from clades with independent sex chromosomes
Chicken chromosome
Carcharodon carcharias
Coilia mystus
Hoplias malabaricus
Argentina silus
Aulostomus maculatus
Girardinichthys multiradiatus
Echiichthys vipera
Gasterosteus aculeatus
Hyla sarda
Ornithorhynchus anatinus
Homo sapiens
Dibamus smithi
Tiliqua scincoides
Shinisaurus crocodilurus
Furcifer pardalis
Bradypodion pumilum
Cyclura pinguis
Vipera latastei
Podarcis raffonei
Gallus gallus
In any VGP SC:
VGP Extended Lineage:
Cartilaginous fish
Ray-finned fish
Amphibians
Mammals
Squamates
Birds
Chicken chromosome is:
Syntenic with sex-shared chromosome
Not syntenic with sex-shared chromosome

**Extended Data Figure 8. HalfDeep analysis for 566 sex chromosomes annotated in 324 species. a,** Chromosome-level HalfDeep proportions of annotated sex chromosomes in homogametic and heterogametic individuals. Each point represents the HalfDeep proportion for one annotated sex chromosome. Seven potential sex mislabelings (XX,ZZ) and partially collapsed ampliconic signals in 48 Y/W chromosomes (HalfDeep proportion < 75%) were detected. Numbers correspond to the following species: (1) monito del monte, (2) small-spotted catshark, (3) common brushtail possum, (4) common myna, (5) common buzzard, (6) lesser black-backed gull, and (7) Mauritius kestrel. **b**, HalfDeep proportion by chromosome type. **c**, HalfDeep proportion vs. sex chromosome length with points colored by chromosome type as in b. **d**, Normalized sequencing depth of Y and W grouped by HalfDeep proportion (<75% or ≥75%). **e**, Chromosome length of Y and W grouped by HalfDeep proportion (<75% or ≥75%). **f,g,** Chromosome-wide sequencing depth and HalfDeep profiles along a Z chromosome in one heterogametic individual (ZW) for Dalmatian pelican (**f**; PacBio HiFi-solo assembly without full phasing) and for Swift parrot (**g**; PacBio HiFi + Hi-C assembly with full phasing). Each grey dot represents sequencing depth in a 1 kbp window. Apricot shading marks HalfDeep regions. Half-depth thresholds were derived from the 40th and 60th percentile depths of the halved genome-wide dataset. **h,** HalfDeep proportion, and **i,** chromosome length, depending on the sequencing approach in 238 heterogametic individuals across multiple species. **j**, orthology of sex chromosomes across 21 species representing 21 independently derived sex chromosome systems, using the chicken (*Gallus gallus*) chromosomes as the reference. Dots represent the homologous sequence in chicken compared to the sequence assembled and annotated as the sex chromosome(s) in each species. The summary along the bottom is all chicken chromosomes that are identified as homologous to a sex chromosome in any species.

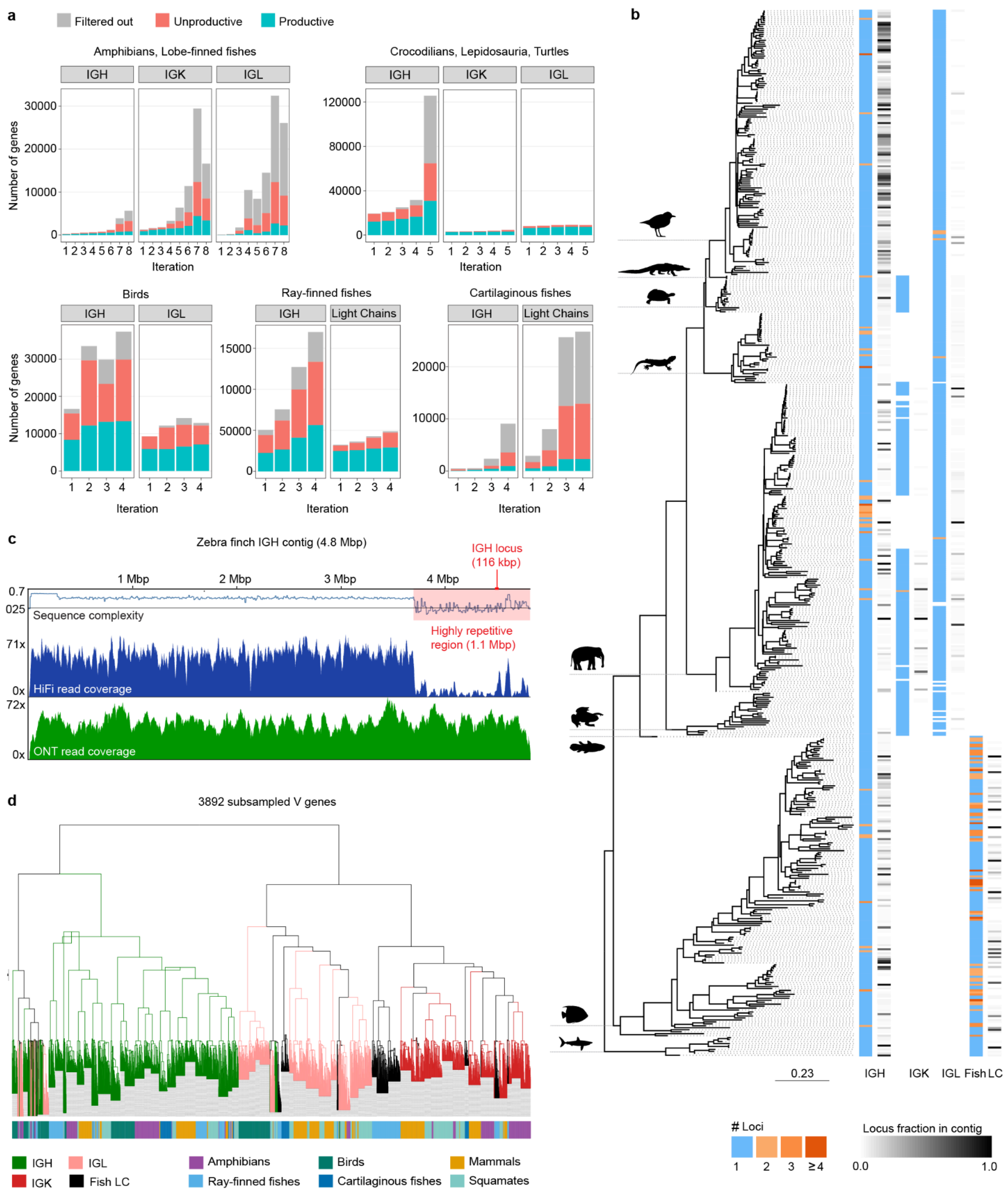


**Extended Data Figure 9. Annotation and analyses of vertebrate immunoglobulin loci. a**, Overview of the number of genes identified through the iterative IgDetective search. For each vertebrate class, the numbers of IG genes discovered at each iteration and across all loci are grouped into three categories: productive, non-productive, and short / low-complexity. The number of iterations used for each class was determined based on the saturation of productive genes and the point at which additional iterations yielded primarily short or low-complexity genes. **b**, A species tree showing the number of IG loci detected per chain type (IGH, IGK, IGL, fish light chain loci) as the blue-orange tracks: 1 contig (blue), 2 or more contigs (shades of orange). For cases when only one locus was detected per chain type, the fraction of the contig taken by the locus is shown in the gray tracks ranging from 0.0 (white) to 1.0 (black). **c**, Analysis of the 4.8 Mbp zebra finch contig containing a 116 kbp IGH locus (red shading). The top track displays the sequence complexity computed as the ratio of homopolymer-compressed length to original sequence length (i.e., fraction of nucleotide switches) plotted across sliding windows; lower values correspond to regions enriched in long or frequent homopolymers. The middle and bottom tracks show read-depth profiles from PacBio HiFi and ONT sequencing, respectively. **d**, The guide tree represents a multiple alignment of 3,892 V genes subsampled from six vertebrate clades and their IG chain types. Four types of immunoglobulin chains are shown by colors of branches: IGH (green), IGK (red), IGL (pink), and fish light chains (black).

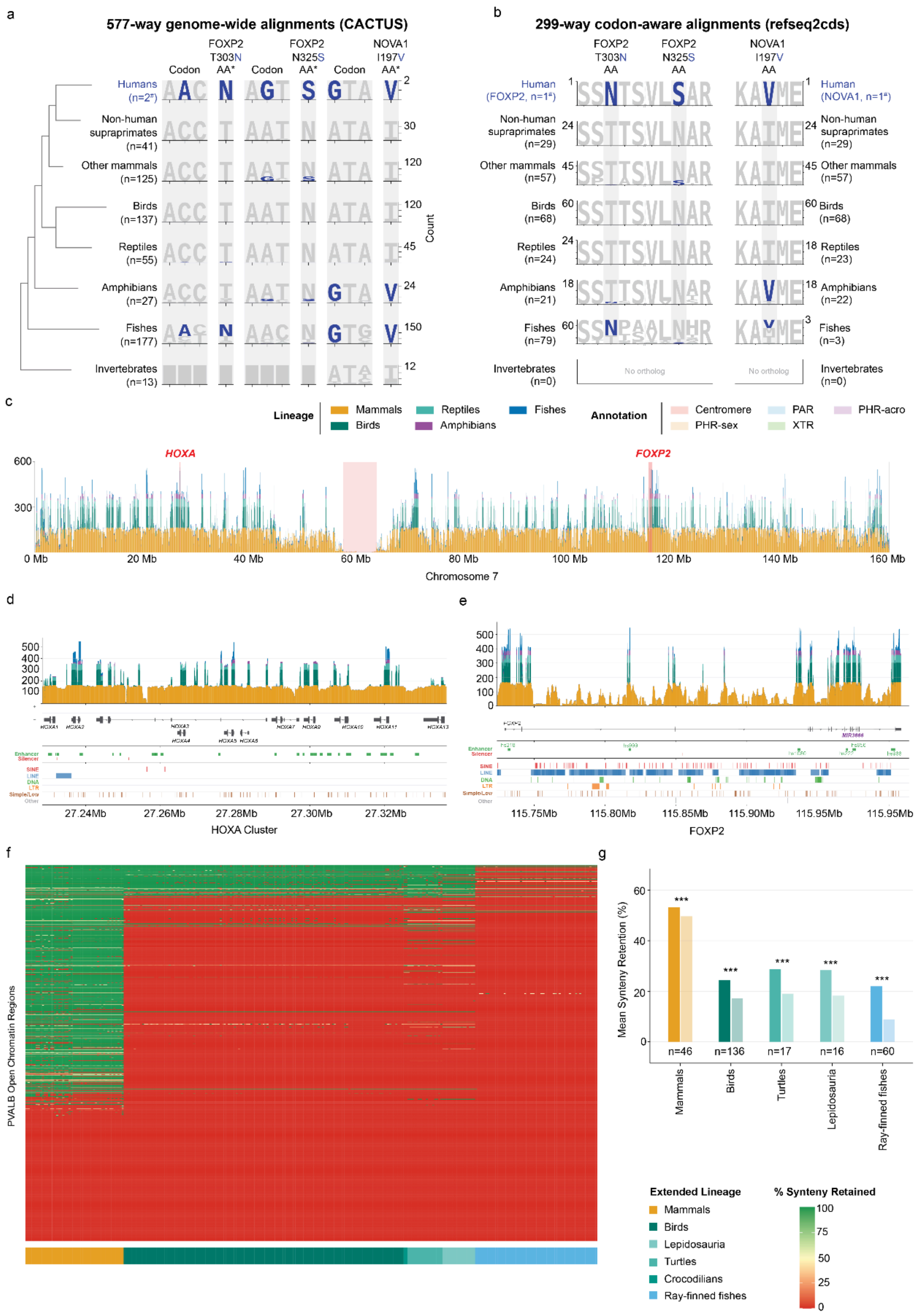


**Extended Data Fig. 10. Candidate amino-acid substitutions and enhancers specialized in humans in highly conserved loci. a**, Phylogenetic distribution of codon and amino acid sites in two genes (FOXP2 and NOVA1) that are specialized in humans. **b**, Amino acid logos for the surrounding 4

sites of the same genes, using a codon aware alignment of 299 of the 579 species that were annotated by EGAPx; NOVA1 in fish were not well annotated and thus have fewer species included. **b**, Genome-wide locus-scale coverage depth intervals containing *HOXA* and *FOXP2*, calculated from 578 assemblies aligned to the T2T CHM13 reference genome included in the VGP dataset. Coverage is shown as stacked vertical profiles, with colors indicating vertebrate lineage: mammals, birds, reptiles, amphibians and fishes. Shaded annotation tracks mark centromeric sequence, PHR-sex, PAR, XTR and PHR-acro regions. **c**, Enlarged view of the *HOXA* locus showing assembly coverage, CHM13 gene models and local sequence annotations. Lineage-specific variation in coverage reflects differences in assembly representation, alignment continuity and sequence divergence across the *HOXA* gene cluster. **d**, Enlarged view of the *FOXP2* locus showing the same lineage-resolved coverage, gene annotation and local marker tracks. Regions with reduced or uneven coverage indicate lineage- or assembly-dependent differences in mappability across this conserved regulatory locus. **e,** Per-peak synteny retention of PV+ interneuron open chromatin regions (n=1,011) across 280 annotated VGP vertebrate genomes, ordered by extended vertebrate lineage. Each row represents one peak sorted by mean synteny retention (lowest to highest), and color indicates retention from high (green) to low (red) synteny. **f,** Mean synteny retention of PV+ interneuron vocal learning-associated open chromatin regions (full color, n=1,011) compared to background PV+ open chromatin regions (reduced opacity, n=16,099) across vertebrate lineages represented in the VGP Phase I 577 species alignment. For each lineage, the signal bar (left) precedes the background bar (right), while n indicates the number of VGP genomes per lineage with signal. Significance assessed by Wilcoxon rank-sum test with Benjamini-Hochberg correction (*** indicates $p<0.001$).

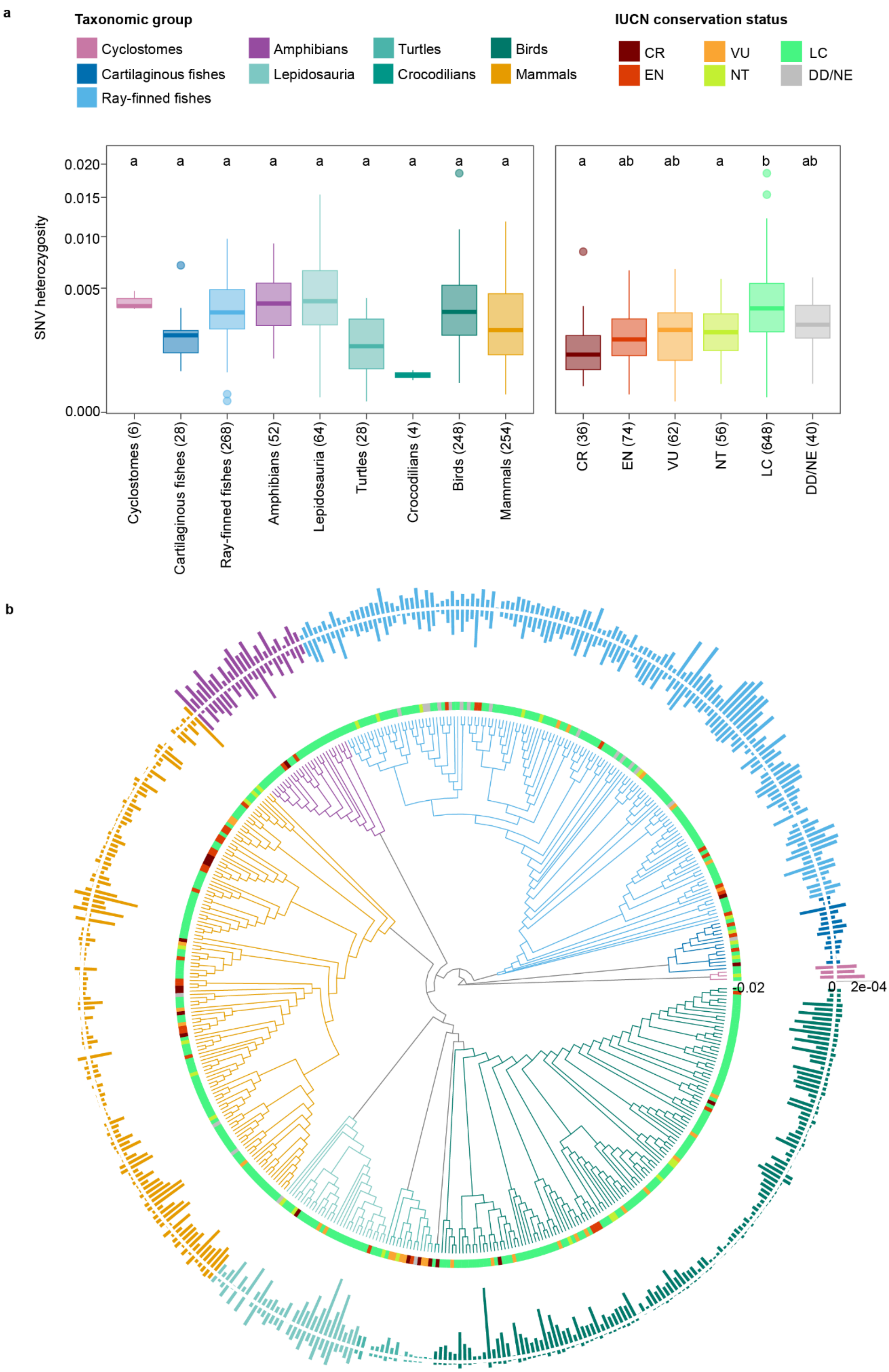
a
Taxonomic group
Cyclostomes
Cartilaginous fishes
Ray-finned fishes
Amphibians
Lepidosauria
Turtles
Crocodilians
Birds
Mammals
IUCN conservation status
CR
EN
VU
NT
LC
DD/NE
SNV heterozygosity
0.020
0.015
0.010
0.005
0.000
a a a a a a a a a
a ab ab a b ab
Cyclostomes (6)
Cartilaginous fishes (28)
Ray-finned fishes (268)
Amphibians (52)
Lepidosauria (64)
Turtles (28)
Crocodilians (4)
Birds (248)
Mammals (254)
CR (36)
EN (74)
VU (62)
NT (56)
LC (648)
DD/NE (40)
b
-0.02
0
2e-04

**Extended Data Fig. 11. SNV and SV diversity across the vertebrate tree of life. a**, Distribution of SNV heterozygosity for 476 high-quality haplotype-resolved VGP Phase I assemblies grouped by major vertebrate clade (left panel) and IUCN conservation status (right panel). CR: Critically Endangered; EN: Endangered; VU: Vulnerable; NT: Near Threatened; LC: Least Concern; DD: Data Deficient; NE: Not Evaluated. Letters indicate whether statistically significant differences were found between groups. **b**, Levels of SNV and SV heterozygosity in individual species shown as bars around the phylogeny (SNV in the inner circle; SV in the outer circle). Colors of the tree and the bars correspond to the major clade each species belongs to in (**a**). The IUCN conservation status of each species is indicated by the heatmap at the tips of the tree.


1. OpenTreeofLife *et al.* Open tree of life taxonomy. Preprint at https://doi.org/10.5281/ZENODO.3937750 (2019).
2. Gupta, A., Mirarab, S. & Turakhia, Y. Accurate, scalable, and fully automated inference of species trees from raw genome assemblies using ROADIES. *Proc. Natl. Acad. Sci. U. S. A.* **122**, e2500553122 (2025).
3. Zhang, C., Scornavacca, C., Molloy, E. K. & Mirarab, S. ASTRAL-Pro: Quartet-Based Species-Tree Inference despite Paralogy. *Mol. Biol. Evol.* **37**, 3292–3307 (2020).
